# EXOCYST70A3 controls root system depth in Arabidopsis via the dynamic modulation of auxin transport

**DOI:** 10.1101/559187

**Authors:** Takehiko Ogura, Christian Goeschl, Daniele Filiault, Madalina Mirea, Radka Slovak, Bonnie Wolhrab, Santosh B. Satbhai, Wolfgang Busch

## Abstract

Root system architecture (RSA), the distribution of roots in soil, plays a major role in plant survival. RSA is shaped by multiple developmental processes that are largely governed by the phytohormone auxin, suggesting that auxin regulates responses of roots that are important for local adaptation. However, auxin has a central role in numerous processes and it is unclear which molecular mechanisms contribute to the variation in RSA for environmental adaptation. Using natural variation in *Arabidopsis*, we identify *EXOCYST70A3* as a modulator of the auxin system that causes variation in RSA by acting on PIN4 protein distribution. Allelic variation and genetic perturbation of *EXOCYST70A3* leads to alteration of root gravitropic responses, resulting in a different RSA depth profile and drought resistance. Overall our findings suggest that the local modulation of the pleiotropic auxin pathway can gives rise to distinct root system architectures that can be adaptive in specific environments.

## INTRODUCTION

The root system enables plants to anchor themselves in the soil and forage their environment for nutrients and water. The evolution of this system 400 million years ago allowed plants to efficiently colonize and transform the land surface of our planet (Algeo and Scheckler, 1998; Gibling and Davies, 2012), paving the way for the highly diverse ecosystems that occupy its landmasses today. The modern root system of dicot species, such as the model plant *Arabidopsis thaliana*, consists of a single primary root that emerges from the seed coat, often remains active throughout the plant’s lifespan, and can develop several orders of lateral roots (Atkinson *et al*., 2014; Peret *et al*., 2009). The apical growth of each root in a root system is driven by cell divisions and subsequent cell expansions in a stem-cell containing structure called the root apical meristem that is located in each root tip (Benfey and Scheres, 2000). Depending on the direction of root growth, different areas in the soil are explored (i.e. a downward growing root will explore the soil vertically and a sideways growing root will explore it horizontally). The number and position of lateral roots add to this complexity and determine which soil layers are intensively explored. All these processes shape root system architecture (RSA) – the spatial distribution of the roots in the soil (Jung and McCouch, 2013). RSA determines important physiological properties of a root system including nutrient and water uptake and anchoring in the soil (Den Herder *et al*., 2010). Despite the importance of RSA, the genes and molecular mechanisms that govern RSA in natural environments remain largely unknown nor do we fully understand which RSA types confer fitness to a particular environment.

Various basic developmental processes that shape RSA including root elongation, gravitropism and lateral root development are regulated by the phytohormone auxin (Pierik and Testerink, 2014). Therefore, it is most likely that auxin plays an important role in the variation in RSA in different environments. However, auxin is a ubiquitous regulator of almost every aspect of plant growth and development at the molecular, cellular, tissue, and organ levels (Benjamins and Scheres, 2008; De Rybel *et al*., 2009). It is therefore not clear which genetic and molecular mechanisms modulate auxin signaling to give rise to the observed natural variation of RSA without disrupting general auxin pathway.

Here, using natural variation and genome wide association mapping in *A. thaliana*, we identify a gene, *EXOCYST70A3*, which regulates RSA by controlling the auxin pathway but independently from other general auxin-dependent growth processes. *EXOCYST70A3* affects the distribution of the PIN4 protein, suggesting that a PIN4-related auxin pathway can be specifically modulated by a component of the exocytosis pathway. We show that allelic variation and genetic perturbation of *EXOCYST70A3* lead to alterations in the orientation of root growth, resulting in a shift from shallow to deep root systems. Finally, guided by data showing that the distribution of *EXOCYST70A3* alleles is highly correlated with precipitation seasonality, a measure of rainfall patterns, we show that overexpression and allelic variation of *EXOCYST70A3* affect drought resistance and conclude that *EXOCYST70A3* variation may play an adaptive role in areas with variable rainfall patterns.

## RESULTS

### *EXOCYST70A3* is associated with natural variation of agravitropic root growth upon auxin transport perturbation

To identify components which regulate RSA related traits by the regulation of auxin pathway, we capitalized on the extensive natural variation of root growth among natural accessions of the model plant *A. thaliana* (Gifford *et al*., 2013; Slovak *et al*., 2014). In particular, we applied a chemical genetics approach using the auxin transport inhibitor 1-*N*-naphtylphthalamic acid (NPA, 10 μM) and assessed the sensitivity of 215 *A. thaliana* natural accessions (Table S1) to perturbation of auxin transport (Figure 1A). Notably, the response to NPA in terms of root growth direction (horizontality, an index of root agravitropism which captures the extent of root growth in the horizontal direction, see supplementary information for details of the trait), a key trait for RSA, was highly variable among accessions in comparison to other traits including total length (Figure 1B, S1). Using genome wide association (GWA) mapping to identify loci underlying the variation of horizontality upon NPA treatment, we detected a peak containing a highly significantly associated <u>S</u>ingle <u>N</u>ucleotide <u>P</u>olymorphisms (SNP) on chromosome 5 (Figure 1C-D, Table S2). This SNP showed considerable effect size (13.58%), suggesting a strong allele effect of the locus which was tagged by this SNP. Using haplotype analysis of the 40 KB region centered on this SNP, we classified the tested accessions into two haplogroups and observed that one haplogroup was significantly associated with greater horizontality (population structure corrected association test: *p*-value = 4.3 × 10^-3^, Figure S2A), indicating that accessions in this haplogroup show increased responses in root growth direction to NPA. Hereafter, this haplogroup is referred to as haplogroup H (<u>H</u>igh-response to NPA, containing 16 accessions) and the other as haplogroup C (<u>C</u>ommon-response to NPA, containing 199 accessions including Col-0).

**Figure 1.**
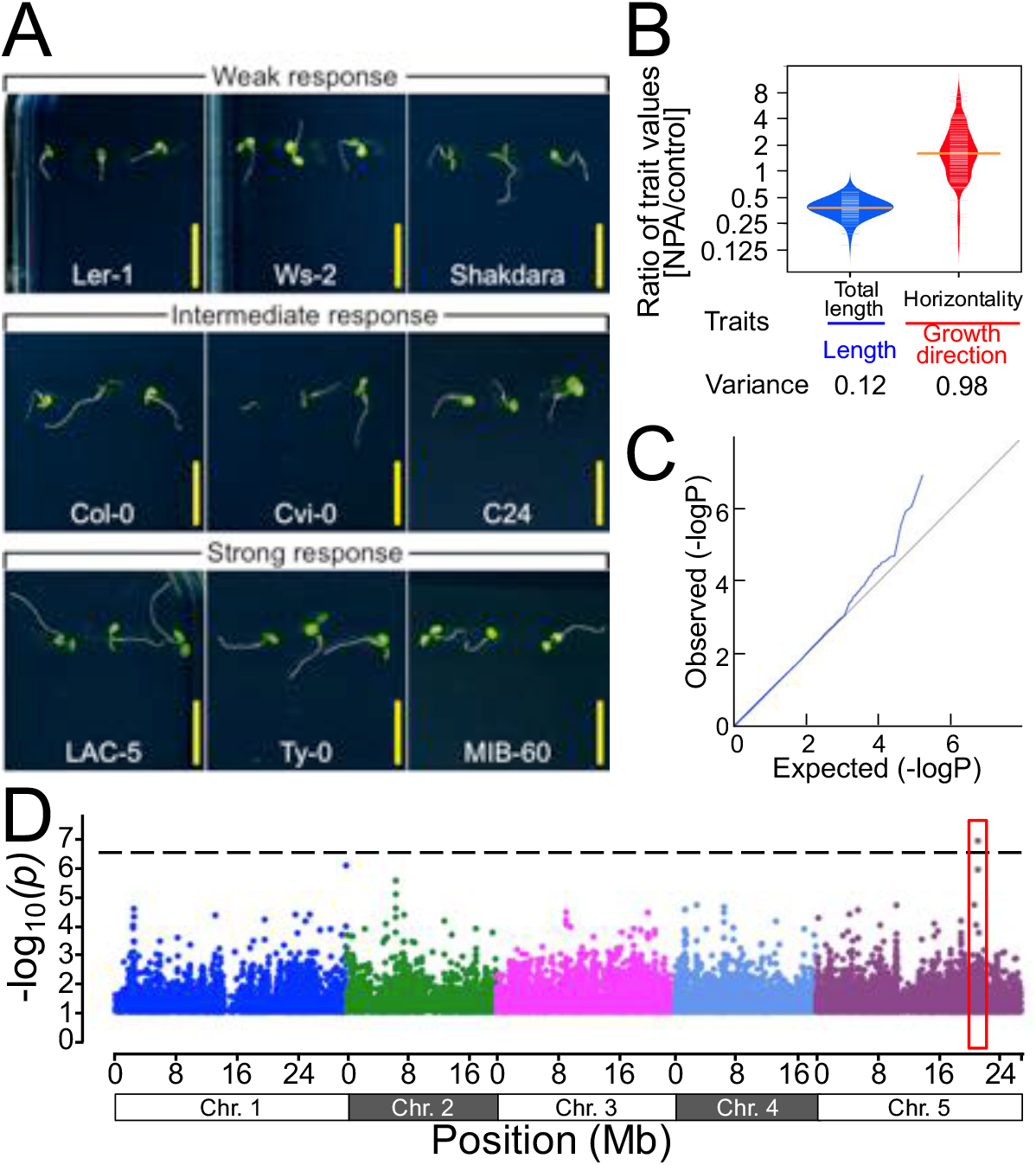
Natural variation of root growth responses upon NPA treatment. **(A)** Representative accessions of *A. thaliana* that show weak, intermediate, and strong agravitropic growth on medium containing 10 μM NPA at 6 days after germination (DAG). Yellow bars: 1 cm. **(B)** Distribution of phenotypes of NPA treated roots. Fold changes of trait values of NPA treated accessions in comparison to untreated accessions were compared between two traits (total length and horizontality, see supplementary information for details of traits.) x-axis: trait; y-axis: fold change of traits under NPA treatment. **(C)** Quantile-quantile plot of *p*-values that were calculated by GWAS. The line matching between x-y and x=y lines in the low *p*-value range suggests no significant cofounding in GWAS. x-axis: expected *p*-values of SNPs; y-axis: observed *p*-values of SNPs. **(D)** Manhattan plot for GWA mapping of the root angle phenotype. Manhattan plot for the SNP associations to horizontality were obtained from GWAPP (Seren *et al*., 2012). Chromosomes are depicted in different colors. The horizontal dash-dot line corresponds to a nominal 0.05 significance threshold after Bonferroni correction. Red box indicates the significantly associated locus. See also Figure S1 and S2 and Table S1 and S2.

Having identified a genomic region associated with control of root growth direction, we set out to identify the causal gene (Table S2, Figure S2B). The significantly-associated SNP identified by GWA mapping is located in the protein-coding region of the *EXOCYST70A3* gene (AT5G52350, hereafter called *EXO70A3*). EXO70A3 is a homologue of yeast and Human EXO70, which is one subunit of the octameric vesicle-tethering EXOCYST complex (Kee *et al*., 1997; TerBush *et al*., 1996). Human EXO70 directs vesicles to target locations on the plasma membrane (Matern *et al*., 2001). The association of the root growth direction phenotype with variation in a central component for exocytosis was striking, since its counterpart, endocytosis, is recognized as one of main regulatory processes for localization of the PIN-FORMED (PIN) auxin efflux carriers and therefore critical in regulating auxin signaling (Petrasek and Friml, 2009). Importantly, a microarray based high-resolution root expression atlas (Brady *et al*., 2007) showed that *EXO70A3* transcripts are specifically expressed in the root tip and the elongation zone (Figure 2A, S3A-D), which are the main sites for gravity perception and directional growth responses, respectively (Masson *et al*., 2002). In line with this, an independently generated RNA-Seq based atlas also showed columella cell-specific expression of *EXO70A3* (Figure S3E, Li *et al*., 2017). Consistent with both these studies, we could only detect expression of *EXO70A3* in the root tip and not in whole roots or shoots (Figure 2B). Taken together our data strongly suggested the involvement of *EXO70A3* in the determination of root growth direction via an auxin pathway (designated ARD: <u>A</u>uxin-dependent <u>R</u>oot <u>D</u>irectional growth), and led us to further analyze *EXO70A3* function.

**Figure 2.**
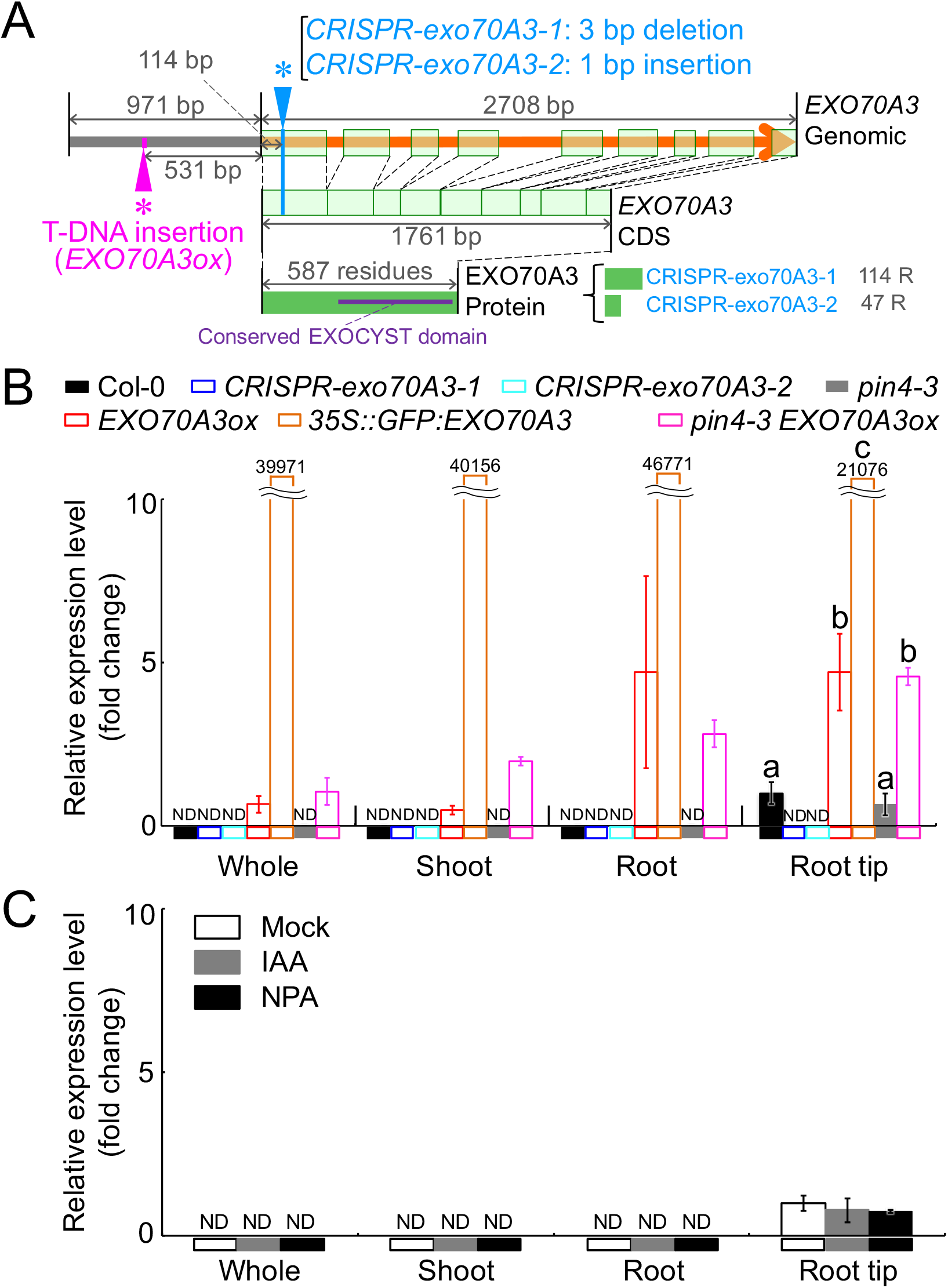
*EXO70A3* expression levels and gene models in *EXO70A3-related* lines. **(A)** Gene models of *EXO70A3* mutants. Orange arrow: genomic sequence of coding region; Light green boxes: exon; Gray bar: intergenic region; Green boxes: amino acid sequence; Lines indicated by arrowheads and asterisks: points of mutation. See also Figure S2 and S3 and Table S1 and S2. **(B)** Transcript levels of *EXO70A3* in different tissues of seven *A. thaliana* lines. x-axis: sample; y-axis: relative expression level (normalized to Col-0 root tip); Error bars: standard deviation; ND: not detected; Numbers above orange bars: fold change of *EXO70A3* transcript level in each tissue of *35S::GFP:EXO70A3* in comparison with that in Col-0 root tip; “a”, “b” and “c”: significance evaluated by ANOVA (*p*-value < 0.05, n = 3). **(C)** Transcript levels of *EXO70A3* in different tissues of chemically treated Col-0 plants. x-axis: sample; y-axis: relative expression level (normalized to mock-treated root tip); Error bars: standard deviation; ND: not detected.

### *EXO70A3* alleles cause natural variation of responses to NPA in roots

To characterize *EXO70A3*, we obtained a T-DNA line (SALK_075426; hereafter called *EXO70A3ox*) which has a promoter insertion resulting in the strong overexpression of *EXO70A3* (Figure 2A, B) but not of the surrounding genes (Figure. S3F, G). In addition, we generated another overexpression line that ubiquitously highly expresses *EXO70A3* using a *35S* promoter (*35S::GFP:EXO70A3*), and two *CRISPR/CAS9-based* knockout lines (*CRISPR-exo70A3-1* and *CRISPR-exo70A3-2*) each containing an early stop codon (Figure 2A, B). The effect of NPA on the direction of root growth of these four mutants was tested on MS medium supplemented with a low dose of NPA (0.5 μM) or control medium. The direction of root growth on medium containing NPA was significantly more agravitropic in the four mutant lines of *EXO70A3* (two-way ANOVA, genotype-treatment interaction *p*-value < 0.05, Figure 3A), suggesting that ARD is misregulated in both, knockout and overexpression lines of *EXO70A3* due to the loss of the appropriate spatio-temporal expression pattern of *EXO70A3*. We then tested whether *EXO70A3* itself is auxin regulated, but neither IAA nor NPA up/downregulated *EXO70A3* expression levels in Col-0 (Figure 2C), indicating that *EXO70A3* function itself is auxin-independent.

**Figure 3.**
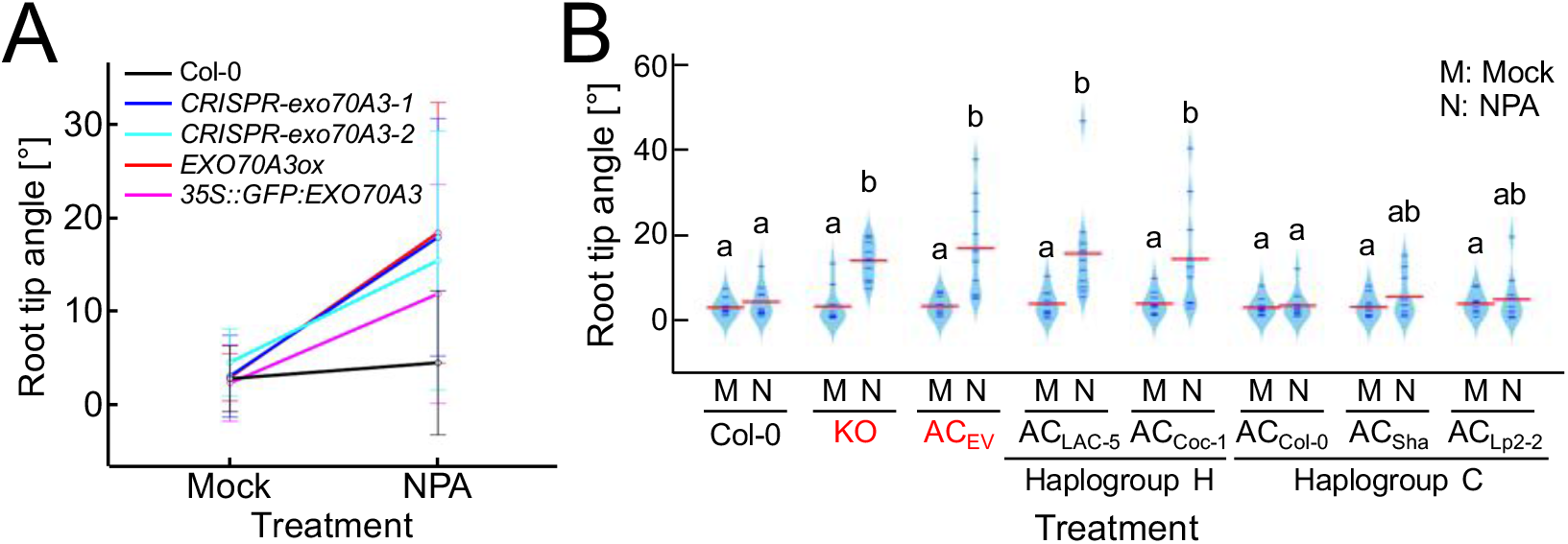
Natural *EXO70A3* variants determine high-response to NPA. **(A)** Root growth orientation response to NPA of Col-0 and EXO70A3-related mutant plants on mock and 0.5 μM NPA media. x-axis: treatment; y-axis: angle changes of root tips at 1 day after transfer (DAT) from 1 × MS plates to treatments medium at 5 DAG. **(B)** Beanplot showing root tip angles of allelic complementation lines on mock and 0.1 μM NPA media. KO: *CRISR-exo70A3-1;* AC: allelic complementation lines that were prepared by transforming *CRISR-exo70A3-1* with vector control (AC_EV_) or *EXO70A3* alleles from Col-0 (AC_Col-0_), Shakdara (AC_Sha_), Lp2-2 (AC_Lp2-2_), LAC-5 (AC_LAC-5_) and Coc-1 (AC_Coc-1_). Red characters indicate mutant-type alleles. x-axis: treatments; haplogroup of alleles is indicated below x-axis; y-axis: angle changes of root tips at 1 DAT from 1 × MS plates at 5 DAG. Red lines: mean; short lines: individual data points; shape: density of the data distribution; “a” and “b”: significant difference evaluated by post hoc Tukey test after ANOVA (*p*-value < 0.05, n ≥ 10). See also Figure S2.

Next, we tested the causality of alleles that we had identified in our GWAS by allelic complementation analysis. We therefore fused *EXO70A3* promoter sequence from Col-0 to the protein coding region of *EXO70A3* of five natural accessions (Haplogroup C: Col-0, Shakdara (Sha), Lp2-2. Haplogroup H: LAC-5, Coc-1. Figure S2C) and transformed these constructs into the *CRISPR-exo70A3-1* line. We reasoned that if the *EXO70A3* alleles of the two haplogroups were underlying the observed phenotypes, they should complement the knockout mutant phenotype to a different extent.

While we could not observe any complementation of the knockout mutant phenotype with an empty vector control, we achieved full complementation with the Col-0 allele belonging to haplogroup C (ANOVA: *p*-value < 0.05, Post hoc Tukey test: *p*-value < 0.05, Figure 3B). In contrast to the C haplogroup Col-0 *EXO70A3* allele, all complementations with haplogroup H alleles (Lac-5, Coc-1) resulted in similar phenotypes with *EXO70A3* mutants and a natural accession from haplogroup C, Coc-1 (Figure S2D), and did not lead to a significant complementation of the NPA-induced root growth direction phenotype (Figure 3B). Complemention with further haplogroup C accession alleles (Sha, Lp2-2) were in between the Col-0 complementation and the haplogroup H non-complementation (Figure 3B). Consequently, when testing whether haplogroup H and haplogroup C were different in their ability to complement, the 2-way ANOVA resulted in a high significance (haplogroup - treatment interaction *p*-value = 9.36×10^-7^), demonstrating that allelic variation of *EXO70A3* determines natural variation of ARD. Since the two constructs that were created with *EXO70A3* sequences from haplogroup H contained single non-synonymous SNP which exists only in haplogroup H in our analysis (Chromosome 5, 21257230, Figure S2C), this coding change is the prime candidate for conferring this effect.

### *EXO70A3* specifically regulates gravitropic responses of PIN4 in columella cells

Since allelic variation of *EXO70A3* was causal for the natural variation of the root response to NPA, we set out to investigate the mechanism for the involvement of *EXO70A3* in the regulation of auxin signaling. We first assessed the effects of *EXO70A3* on the localization patterns of the PIN auxin efflux carriers by crossing *EXO70A3ox* with widely used *PIN::PIN:GFP* plants. Of the eight members of the Arabidopsis PIN family, five PINs (PIN1, PIN2, PIN3, PIN4 and PIN7) that are widely recognized for their fuctions in roots were tested. While PIN1, PIN2, PIN3, and PIN7 localization were not substantially affected (Figure S4A), PIN4:GFP was superabundant in columella cells of primary roots but not in stele cells in *EXO70A3ox PIN4::PIN4:GFP* plants (Figure 4A, B). In these plants, almost all cells in the columella tissue showed PIN4:GFP signal while only a fraction of cells in the columella tissue showed signal in the control *PIN4::PIN4:GFP* plants. In both *PIN4::PIN4:GFP* and *EXO70A3ox PIN4::PIN4:GFP* plants, PIN4:GFP signal localizes ubiquitously on the plasma membrane of the cells in which PIN4:GFP signal is present, indicating that the regulation of PIN4:GFP by *EXO70A3* does not lead to a difference at the sub-cellular level. This increase of the PIN4:GFP signal in root tips was also observed in lateral roots although PIN4:GFP signal was absent in early stage lateral roots (Figure S4B). In contrast to the *EXO70A3ox* line, *PIN4:GFP* level was significantly decreased in columella cells but not in the stele of *CRISPR-exo70A3-1 PIN4::PIN4:GFP* plants (Figure 4A, B), supporting the idea that *EXO70A3* specifically regulates PIN4 localization in columella cells.

**Figure 4.**
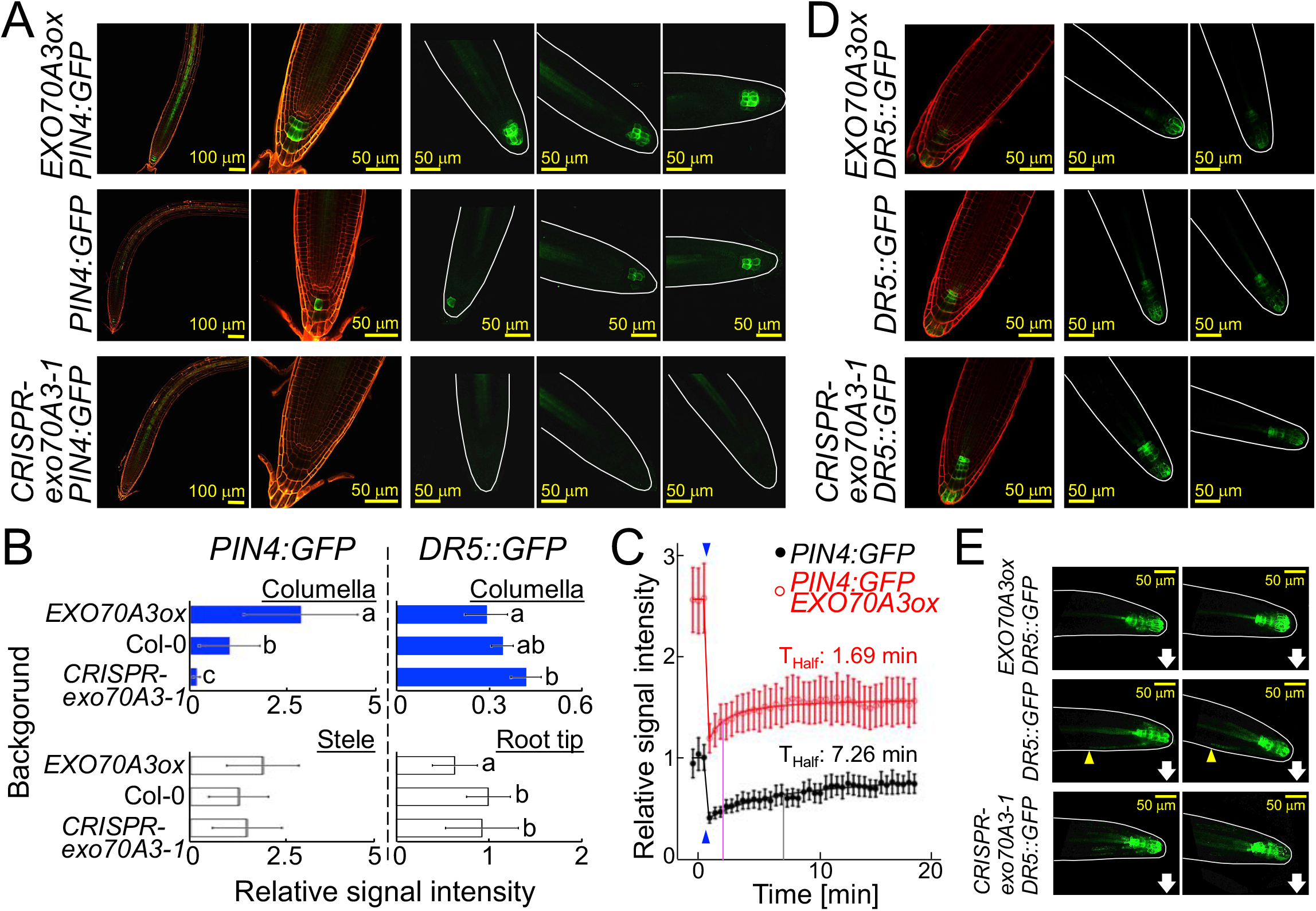
Perturbation of *EXO70A3* expression regulates PIN4 levels and patterns. Confocal images of representative roots of *PIN4::PIN4:GFP, DR5::GFP* or crosses with EXO70A3-related lines grown on 1 × MS plates (5 DAG) and quantitative comparison of fluorescent signal intensities. *PIN4:GFP* indicates *PIN4::PIN4:GFP*. Green: GFP signal; Red: cell wall stained with propidium iodide (PI); White lines: outline of root tip; Yellow bars: 50 μm or 100 μm (indicated in each diagram). **(A)** Median longitudinal optical section of root tips (left panel), a magnification of the left panel (second left panel) and maximum intensity projection (MIP) images (three right panels) of *PIN4::PIN4-GFP, PIN4::PIN4-GFP CRISPR-exo70A3-1* and *PIN4::PIN4-GFP EXO70A3ox*. **(B)** Quantitative comparison of GFP signal intensities among EXO70A3-related mutants. x-axis: average relative signal intensity normalized by that in plants with Col-0 background; y-axis: background line of crosses with *PIN4::PIN4:GFP* (left panels) or DR5::GFP (right panels) that are tested in this diagram; Error bars: standard deviation; “a”, “b” and “c”: significance evaluated by ANOVA (*p*-value < 0.05, n ≥ 10). **(C)** FRAP analysis of the EXO70A3 effects on cellular *PIN4::PIN4:GFP* transport. GFP signal in a columella cell was bleached after three pre-scans and the recovery of signal intensity in the cell was measured. More than 10 cells in different roots were analyzed per line. Signal intensities at all time points in all cells were normalized by the average signal intensity of *PIN4::PIN4:GFP* cells of three pre-scans, and an average value of each line was plotted for each time point. x-axis: relative signal intensity; y-axis: time (minute); Blue arrowheads: point of photo bleaching; T_Half_: Half-time, a calculated time which is necessary to obtain 50% of maximum recovery after photo bleaching, which is indicated by a gray line for *PIN4::PIN4:GFP* and a magenta line for *PIN4::PIN4:GFP EXO70A3ox;* Error bars: standard deviation. **(D)** Median longitudinal optical section of root tips (left panel) and maximum intensity projection (MIP) images (two right panels) of *DR5::GFP, DR5::GFP CRISPR-exo70A3-1* and *DR5::GFP EXO70A3ox*. **(E)** *DR5::GFP* signals at 3 hours after 90° turn. White arrows: the gravity vector. Yellow arrowheads: optically visible ends of basipetally extended GFP signals at the side of gravity vector under gravistimulation. See also Figure S3, S4 and S5.

While the function of *EXO70A3* in exocytosis suggests that the effect on PIN4 is exerted at the protein level, we wanted to test this assumption. For this, we measured PIN4 transcript level in *EXO70A3ox* plants. PIN4 transcript level was not elevated in root tips of these plants (Figure S3G), further suggesting regulation of PIN4 by EXO70A3 at the protein level. To directly test this, we conducted Fluorescence Recovery After Photobleaching (FRAP) experiments determining the exocytosis rate of PIN4. Providing further evidence for the involvement of *EXO70A3* in the exocytosis of PIN4, we found that the overexpression of *EXO70A3* results in the rapid recovery of PIN4:GFP signal on the plasma membrane after photo-bleaching compared to the wildtype background (Figure 4C).

Next we wanted to investigate the effect of EXO70A3 function on auxin transport. For this, we compared expression patterns of the auxin responsive *DR5::GFP* reporter gene between wildtype, *CRISPR-exo70A3-1* and *EXO70A3ox*. We found that there is an inverse relation between PIN4:GFP and DR5::GFP signal intensities in wildtype, *CRISPR-exo70A3-1* and *EXO70A3ox* backgrounds (Figure 4B, D). This suggests that *EXO70A3* modifies auxin levels in root tips through the regulation of PIN4 levels in Columella tissue. Taken together, these results indicate that EXO70A3 specifically regulates PIN4 presence in columella cells, presumably by a process related to PIN4-specific exocytosis, and thereby regulates auxin distribution in root tips.

### Perturbation of *EXO70A3* specifically delays root growth responses to gravistimuli

Since we had aimed to identify mechanisms that potentially alter RSA by modulating auxin pathway, we tested the effect of knockout and overexpression of *EXO70A3* on various auxin pathway-related phenotypes including other RSA traits, primary root length and lateral root density. We did not find evidence for an effect in the two knockout mutants of *EXO70A3* and the two *EXO70A3* ubiquitously overexpressing lines while testing primary root length, root growth response to IAA, lateral root density, and auxin-dependent shoot traits (Figure S5A-D).

In contrast, in line with the result that *EXO70A3* mutants visibly altered root growth orientation of seedlings in the presence of NPA (Figure 3A), an assay of dynamic root gravitropic responses in the absence of the chemical treatment revealed that roots of the *EXO70A3* mutant lines show significant delays in gravitropic responses compared to the wildtype (Figure 5A-D, Movie S1). We therefore conclude that *EXO70A3* specifically regulates ARD but not, at least not to a detectable extent, other auxin-dependent traits.

**Figure 5.**
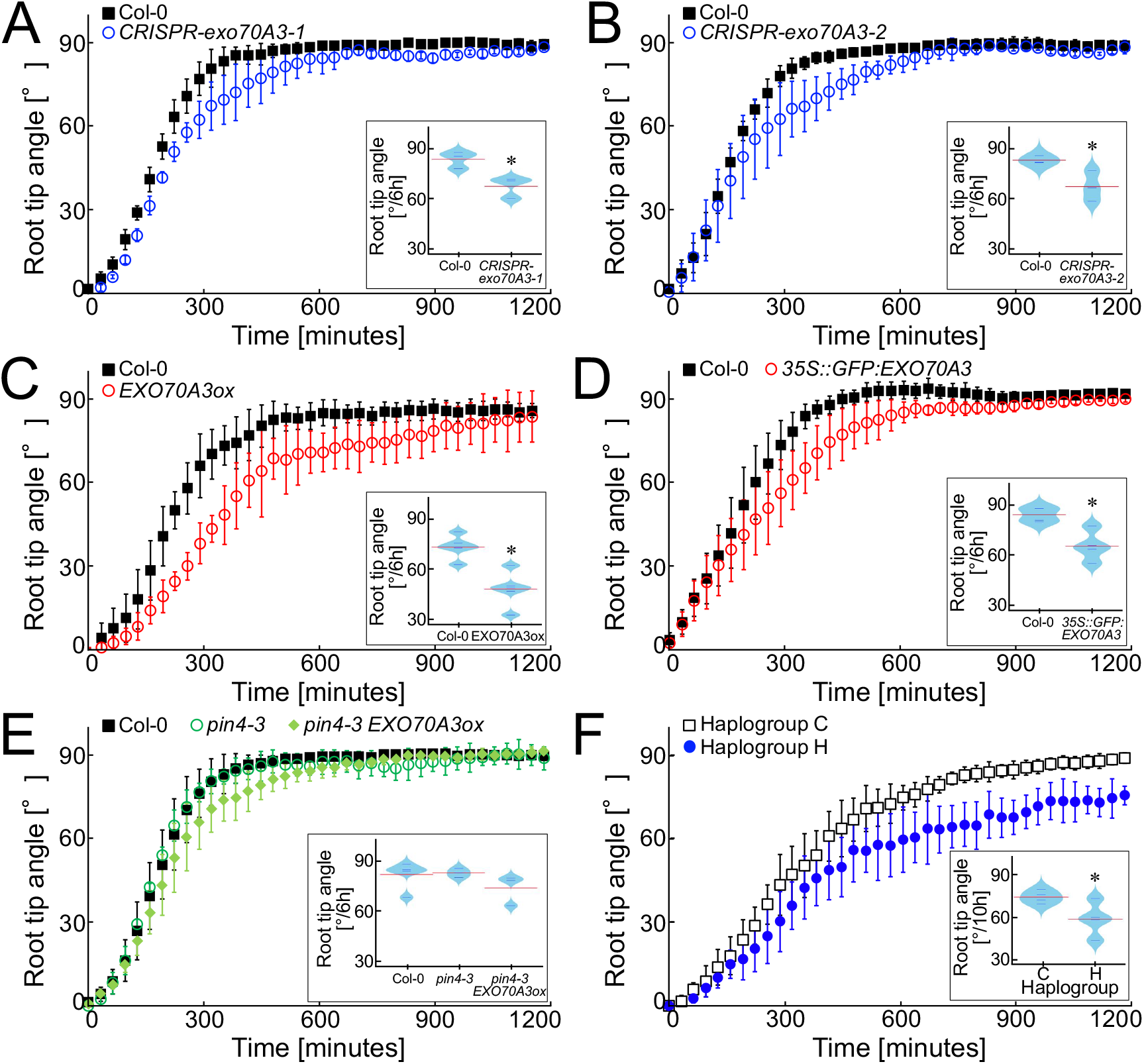
*EXO70A3* regulates rapidness of the gravitropic bending upon gravistimulus. **(A-F)** Re-orientation of root tips after 90° turning of plates measured in 32 minute intervals. *CRISPR-exo70A3-1* **(A)**, *CRISPR-exo70A3-2* **(B)**, *EXO70A3ox* **(C)**, *35S::GFP:EXO70A3* **(D)** and *pin4-3* and *pin4-3 EXO70A3ox* **(E)** were compared with Col-0. In the same way, four natural accessions from Haplogroup C and four natural accessions from Haplogroup H were compared **(F)**. x-axis: time [minutes] after plate turning; y-axis: average angle change of root tip; Error bars: standard deviation; Beanplots: total root tip angle changes of Col-0 and *EXO70A3ox* after 6 or 10 hours; Red lines: mean; short lines: individual data points; shape: density of the data distribution; Asterisks: significance evaluated by Welch t-test (*p*-value < 0.05, n ≥ 3) or significance evaluated by population structure corrected association test for angle changes at 10h after re-orientation (F, *p*-value < 0.05). See also Figure S3 and S5 and Movie S1 and S2.

Even in the root tip, auxin is involved in multiple processes that could have an effect on root growth orientation. We therefore investigated possible causes and first tested whether aberrant development of the root meristem could be the cause for phenotypes. This is not true since neither the cellular root architecture nor the numbers of gravity perceiving statocyte and statoliths in the columella cells are altered in plants of the *EXO70A3* knockout and overexpression lines (Figure S5A-D). Instead, and in line with the altered PIN4 distribution in these lines (Fig. 4A), the *EXO70A3* knockout and overexpression lines showed a perturbation of the auxin distribution pattern, as an asymmetric accumulation of DR5::GFP signal in the downward peripheral layer was observed under gravistimulus in WT plants but not, in most cases (8/11 and 14/20 roots), in *CRISPR-exo70A3-1* and *EXO70A3ox* plants, respectively (Figure 4E). Knockouts and overexpressors of *EXO70A3* showed similar phenotypes, indicating that deregulation of the *EXO70A3* function causes the phenotype. This is supported by the effect on DR5:GFP since both knockout and overexpression of EXO70A3 causes perturbation of auxin responses. Such similarities between overexpression and knockout (knockdown) mutants are frequently observed as shown in a systematic screen in yeast in which 23% of the observed morphological consequences upon gene overexpression were similar to the respective deletion mutant phenotypes (Sopko *et al*., 2006).

The prominent regulation of PIN4 by *EXO70A3* prompted us to investigate the role of *PIN4* in the control of root growth orientation. Much like the EXO70A3 overexpression and mutant lines, *pin4-3* knockout mutants did not show developmental phenotypes in shoot and root (Figure S5E). While the dynamic root gravitropic response was also similar to Col-0 (Figure 5E), the variation of root tip angles during the dynamic root gravitropic response, which is another index of root agravitropism (Yamamoto *et al*., 1998), was significantly larger than that of Col-0 (Figure S5F). The cross of *EXO70A3ox* and *pin4-3* showed this variance phenotype (Figure S5F) of the pin4-3 mutant but did not display the *EXO70A3ox* induced change in the average dynamic root gravitropic response (Figure 5E). This indicates that EXO70A3 is upstream of *PIN4*. Taken together, this suggests that much like in overexpressors of *EXO70A3* and *CRISPR-exo70A3* mutants, the control of gravitropism is exerted less precisely in *pin4-3* mutants, but that a more pronounced disturbance of the quick root gravitropism requires an abnormal localization of PIN4 rather than absence of PIN4. Moreover, additional genes downstream of *EXO70A3* might be needed to delay gravitropic responses.

Importantly, the effects of *EXO70A3* knockout and overexpression on ARD were consistent with the phenotypic difference observed between natural alleles of *EXO70A3* (haplogroup H showed a delay in responding to a gravistimulus (*EXO70A3* mutant type phenotype), population structure corrected association test for angle changes at 10h after re-orientation: *p*-value = 0.037, Figure 5F, Movie S2), suggesting that allelic variation of *EXO70A3* is involved in ARD processes. While the delay in dynamic ARD was consistent with that observed in *CRISPR-exo70A3-1* and *EXO70A3ox* plants, no differences in *EXO70A3* expression were observed between the haplogroups (Figure S3I), suggesting that allelic differences are due to post-transcriptionally effective or coding variation in the EXO70A3 protein itself. This notion is strengthened by the presence of non-synonymous amino acid substitutions in haplogroup H and its causality in NPA response variation (Figure 3B, S2B).

### *EXO70A3* can shift root system architectures from shallow to deep

We have shown that *EXO70A3* regulates RSA-related root traits on MS medium, which is not a root growth substrate in nature. To assess the role of *EXO70A3* activity in physiologically relevant conditions, we measured RSA of mature plants grown in soil. For this, we contemplated the space the roots occupy particularly in a soil volume. The RSA of *CRISPR-exo70A3-1* and *EXO70A3ox*, as well as that of *pin4-3*, showed a significantly higher extent of vertical expansion (decreased shallowness; ANOVA: *p*-value =1.1 × 10^-3^, post-hoc Tukey test: *p*-value < 5.0 × 10^-3^) in comparison to the RSA of Col-0 (Figure 6A-C). Likewise, the *EXO70A3* locus of haplogroup H was significantly associated with an increased vertical expansion of RSA compared to haplogroup C (population structure corrected association test: *p*-value = 8.8 × 10^-5^, Figure 6B, D). These data strongly suggest that allelic variation of *EXO70A3* determines natural variation of RSA configurations in soil, and depending on the *EXO70A3* allele conferring shallow rooting or deep rooting. Interestingly, while *pin4-3* did not show the pronounced delay of gravitropic bending, it did show deeper RSA. This might indicate that the heterogeneity of root angles during bending observed in *pin4-3* and the *EXO70A3* mutants, rather than the quickness of the bending response that is only observed in *EXO70A3* mutants leads to deeper rooting.

**Figure 6.**
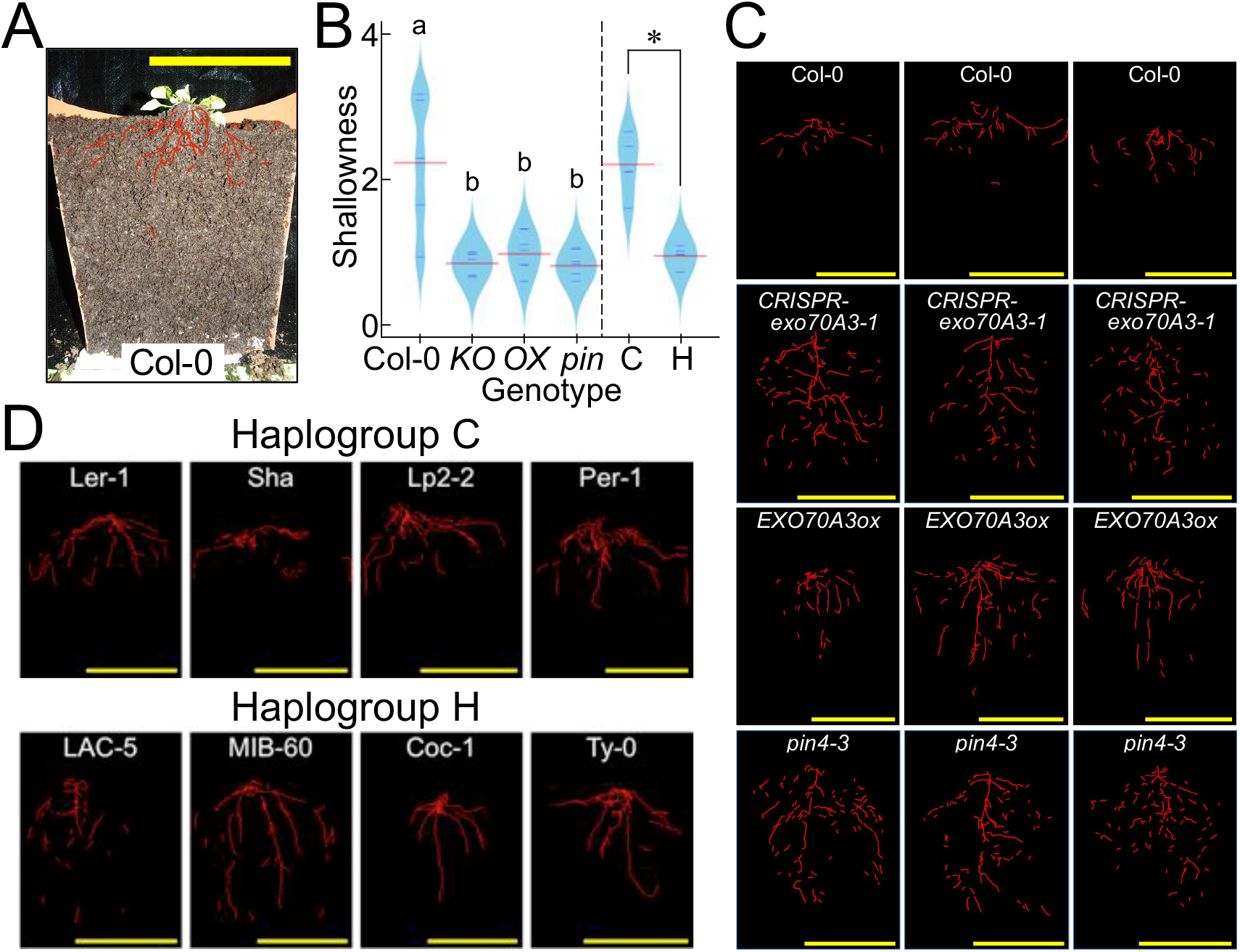
*EXO70A3* alleles impact root system architecture. Vertical-sections of representative root systems for various EXO70A3-related lines and natural accessions grown in soil for 31 days. Red lines: traced roots from the image; Yellow bars: 10 cm. **(A)** Representative images of original vertical-sections of a Col-0 plant. **(B)** Beanplot of the horizontal/vertical expansion (Shallowness) of root systems (31 DAG) in Col-0, *CRISPR-exo70A3, EXO70A3ox, pin4-3* and haplogroups C and H. x-axis: lines; y-axis: Shallowness; KO: *CRISPR-exo70A3-1;* OX: *EXO70A3ox; pin: pin4-3;* C: haplogroup C accessions; H: haplogorup H accessions; “a” and “b”: significance evaluated by ANOVA (*p*-value < 0.05, n = 5) for Col-0 and three mutant lines; Asterisk: significance evaluated by population structure-corrected association test (*p*-value < 0.05, n = 4) for Haplogroup C vs. H. **(C)** Vertical-sections of representative root systems for Col-0, *CRISPR-exo70A3-1, EXO70A3ox* and *pin4-3* (traced roots). **(D)** Vertical-sections of representative root systems for haplogroup C and H accessions (traced roots). See also Figure S7.

### Shallow RSA-type *EXO70A3* leads to increased fitness in drought conditions

Given the remarkable tuning of root traits associated with natural alleles of *EXO70A3*, we hypothesized that this locus is involved in local adaptation. To test this, we exploited data of previous genome-wide scans for signatures of selection (Horton *et al*., 2012) and climate adaptation (Hancock *et al*., 2011). Notably, we found a significant signature within the *EXO70A3* haplotype block (CLR (Kim *et al*., 2007), *p*-value < 0.01, Figure S2, S6A), suggesting a recent selective sweep at this locus. Importantly, we also observed that, in SNP-climate condition correlations that were caluculatd in Hacock *et al*., 2011, the negative correlation between the SNP in *EXO70A3* for which we detected the GWA signal with horizontality and the intensity of precipitation seasonality (WorldClim, http://www.worldclim.org/) was significantly strong (Figure S6B, included in top 5% SNPs). Since this parameter is a measure of the variability in weekly precipitation throughout the year, we reasoned that this correlation might reflect adaptation to temporal drought during the growth period of *A. thaliana*. To test this hypothesis, we assessed whether allelic variation in *EXO70A3* affects fitness under water-limited conditions. Using seed yield as a proxy for fitness, we observed that total seed numbers per plant in control conditions was statistically indistinguishable in *EXO70A3ox* (Welch t-test *p*-value = 0.107) and haplogroup H (ANOVA Haplotype *p*-value = 0.994) from Col-0 and haplogroup C, respectively (Table S3, Figure 7A). In drought conditions, however, it was significantly lower in *EXO70A3ox* (Welch t-test *p*-value = 0.008) and haplogroup H (ANOVA *p*-value = 1.9 × 10^-5^, population corrected association test between C and H haplogroups was marginally significant: *p*-value = 0.052) as compared to Col-0 and haplogroup C, respectively (Table S3, Figure 7B). Moreover, while both Col-0 and *EXO70A3ox* plants produced significantly fewer seeds in response to drought in comparison to well-watered control conditions (Table S3, two-way ANOVA drought treatment: *p*-value < 2.0 × 10^-16^), *EXO70A3ox* responded more strongly to the drought treatment, showing a larger reduction in seed number than Col-0 (Table S3, Figure 7C, two-way ANOVA genotype-treatment interaction: *p*-value = 3.4 × 10^-13^). Similarly, seed production of accessions in both haplogroup H and haplogroup C was significantly reduced in response to drought treatment (Table S3, two-way ANOVA drought treatment: *p*-value < 2.0 × 10^-16^), yet haplogroup H was associated with a greater reduction of seed number in response to drought (Table S3, Figure 7C, two-way ANOVA genotype-treatment interaction: *p*-value < 2.0 × 10^-16^, population structure corrected association test for the response was marginally significant: *p*-value = 0.093). Therefore, the fitness advantage conferred by haplotype C under water-limited conditions is consistent with its correlation with precipitation seasonality. In summary, our data suggest that the haplogroup C allele of *EXO70A3* might have contributed to the adaptation of *A. thaliana* to environments with temporally variable water availability by tuning the RSA to a horizontally-expanded formation.

**Figure 7.**
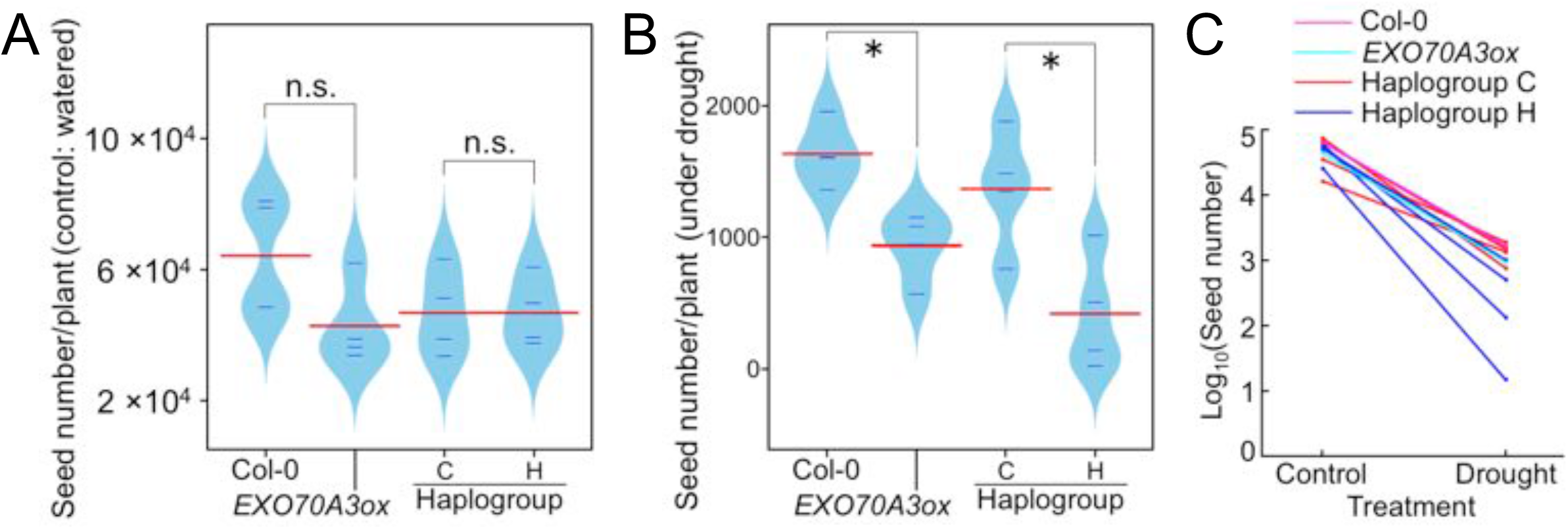
*EXO70A3* alleles associate with fitness in drought conditions. **(A, B)** Beanplots showing distributions of the average seed numbers of Col-0, *EXO70A3ox*, and haplogroups C and H in control (well-watered) (A) and drought conditions (B). x-axis: lines; y-axis: average seed number of four plants grown in one pot (Col-0 and *EXO70A3ox*) or each accession in four blocks of the experimental design (haplogroups C and H). Asterisk: significance evaluated by ANOVA (*p*-value < 0.05); n.s.: not siginificant. **(C)** Reaction norms for seed numbers under control (wet) and drought conditions. x-axis: treatment; y-axis: average seed numbers of Col-0 (magenta line), *EXO70A3ox* (cyan line), and haplogroups C (red lines) and H (blue lines). See also Figure S6 and S7 and Table S3.

## DISCUSSION

### Natural variation of auxin transport dependent early root growth traits

Auxin signaling is a major regulatory pathway for root developmental processes such as cellular proliferation and differentiation (Dello Ioio *et al*., 2008; Ishida *et al*., 2010), cellular expansion (Muday *et al*., 2012), root growth direction (Band *et al*., 2012), and lateral root formation (Lavenus *et al*., 2013); these traits are critical for determining root system architecture. Despite the prominent role of the auxin pathway in regulating root growth and recent progress in revealing genetic bases of natural variation of root growth (Gifford *et al*., 2013; Meijon *et al*., 2014; Slovak *et al*., 2014; Satbhai *et al*., 2018), it remained unclear whether and how the auxin pathway itself is subject to natural variation. Here we demonstrated that there is significant natural variation in response to perturbation of auxin transport for root growth orientation control, one trait regulated by auxin-controlled processes. While the change of root length in different accessions upon auxin transport perturbation was rather homogenous, the change of root growth orientation displayed a much higher variability (Figure 1B, S1). It is therefore tempting to speculate that the auxin pathway-related components that govern root growth orientation are subject to natural variation to a higher extent than those that govern root growth rate. It follows that auxin-dependent regulatory mechanisms of root elongation and root growth orientation are independent. This idea is consistent with recent work that has shown that the change of root growth direction in response to gravity stimuli is mediated by rapid changes in auxin fluxes, while these rapid changes do not affect auxin dependent root apical meristem zonation (Mahonen *et al*., 2014).

### Sequence variation at the *EXO70A3* locus determines how roots respond to gravity

Using GWA mapping and subsequent genetic and molecular analyses, we determined that *EXO70A3* is the causal locus that underlies a significant proportion of the response in terms of root growth orientation to auxin transport inhibition. We base this conclusion on multiple lines of evidence. First, this gene is in a haplotype block centered around the marker SNP which showed the highest signal of association (Figure S2), and according to the 1001 genome sequencing project (1001genomes.org), the high response of haplogroup H to NPA is linked with non-synonymous amino acid changes that were exclusively found in the *EXO70A3* coding region of this group (Figure S2). Indeed, alleles containing this non-synonymous polymorphism were causal for the variable response of natural accessions to NPA (Figure 3B). In addition, the expression pattern of *EX070A3* in the root is specific to tissues with key functions in root growth direction control (Figure 2B, S3A-E). Moreover, natural alleles as well as abnormal distribution of EXO70A3 affect root orientation responses (Figure 5). Furthermore, *EXO70A3* is functionally linked to auxin transport regulation that is relevant to root orientation control at the cellular and molecular level. Overexpression of *EXO70A3* results in overaccumulation of PIN4 in columella cells and *exo70a3* knockout eliminated PIN4:GFP signal from columella cells, altering auxin signal localization patterns during root gravitropism (Figure 4A-D). Since the mRNA expression level of PIN4 in *EXO70A3ox* is indistinguishable from that of Col-0 wildtype (Figure S3H), EXO70A3 is an exocytosis factor (Synek *et al*., 2006), and *EXO70A3* level corresponds with PIN4-GFP signal level, it is highly likely that EXO70A3 regulates the abundance and localization of PIN4 at the plasma membrane. Consistent with this model, FRAP analysis demonstrated the involvement of *EXO70A3* in PIN4 exocytosis (Figure 4C). In sum, these results suggest that EXO70A3 regulates plasma membrane localization of PIN4 by exocytosis and thereby determines root gravitropic responses (Figure S7). The non-synonymous amino acid changes in different natural *EXO70A3* alleles would affect this process and eventually cause the natural variation observed in the modulation of the gravitropic response by NPA (Figure 1A). A detailed RNA-Seq analysis in root cells showed that the highest expression of *EXO70A3* in the data set was detected under cycloheximide treatment (Li *et al*., 2017, Figure S3E), suggesting that *EXO70A3* is tightly regulated by protein degradation. This might indicate that the regulation of related components including auxin signaling factors and auxin transporters such as PINs at protein level might play important roles in the rapid response of roots to gravistimuli.

### The auxin pathway can be specifically modulated by a component of the exocytosis pathway

We have identified a mechanism which controls the auxin pathway and establishes variation in RSA configurations via modulation of the auxin transport machinery. Interestingly, despite the well-known contributions of auxin efflux carriers and endocytosis factors to auxin transport, we did not identify any of the known auxin transporters and endocytosis factors. Instead, we identified a component of the exocytosis pathway. In comparison with intensely studied auxin carriers and endocytic regulators for auxin transport (Kleine-Vehn and Friml, 2008), exocytic components have not been regarded as of primary importance (Kleine-Vehn *et al*., 2011) although the involvement of several general exocytosis factors in the transport of PIN proteins has been recently reported (Feraru *et al*., 2012; Hazak *et al*., 2010; Naramoto *et al*., 2014; Tanaka *et al*., 2014). In particular, the involvement of the *EXOCYST* family has not been well investigated despite the remarkable radiation of family members of EXOCYST-subunit encoding genes in landplants (Cvrckova *et al*., 2012). Of these, only the role of the ubiquitous *A. thaliana* exocytosis factor, *EXOCYST70A1*, in general cellular recycling had been shown to involve regulation of the PIN1 and 2 auxin transporters, thereby affecting numerous traits including shoot and root phenotypes (Drdova *et al*., 2013). Moreover, the importance of EXOCYSTs in auxin transport and signaling is still debated (Cole *et al*., 2014). Our data show a clear and specific role of an EXOCYST complex component, EXO70A3 on auxin-dependent growth through the regulation of PIN4 localization. The specific function of *EXO70A3* detailed here is consistent with the model that specific EXOCYST complexes containing a particular EXOCYST70 isoform can have distinct tissue- or process-specific roles in land plants. While the genetic perturbation of many auxin pathway genes leads to severe and often highly pleiotropic effects on multiple traits (Friml *et al*., 2002; Tsugeki *et al*., 2009), *EXO70A3* represents an uncharted layer of the regulation of auxin signaling system that modulates the swiftness of auxin-dependent control of the root gravitropism by conferring the flexibility of auxin transport through PIN4 without significantly disrupting the auxin pathway itself in root tips.

### *EXO70A3* controls root system architecture, providing adaptive value for drought conditions

Our data strongly suggest that EXO70A3-dependend variation in the control of root growth direction can result in either a shallow or a deep RSA (Figure 6). A major part of the function of *EXO70A3* is exerted by its downstream target PIN4 as lack of *PIN4* in the *pin4-3* mutant leads to a deeper RSA in a same way as dysregulation of *EXO70A3* does. As *pin4-3* plants showed the increased variation of root tip angles while it did not show the delay of gravitropic root bending, it is indicated that downward growth of roots in the soil does not simply reflect the rapidness of bending upon gravity change on agar plates. There are various other stimuli that regulate root growth directions in the soil in addition to gravistimulus. Functions of *EXO70A3* in response to these stimuli in collaboration with that to gravistimulus might fully explain RSA formation in the soil.

The distribution of the *EXO70A3* alleles is highly associated with precipitation seasonality, the variability in weekly precipitation throughout the year (Figure S6B). Importantly, plants with different RSA configurations conferred by variation of the *EXO70A3* locus seem to display different fitness in environments with variable rainfall patterns (Figure S7). In particular, our data indicate a beneficial effect of a shallow, more horizontally distributed RSA in *A. thaliana* under drought conditions. Since surface soil water is considered to be important in environments with sparse rainfall (Lauenroth *et al*., 2014; Nippert and Knapp, 2007), a shallow root system might be an adaptation for efficiently capture the water in the short time that it is available. This idea is consistent with findings that, in water-limited ecosystems, there are significant positive correlations between precipitation seasonality and rooting shallowness in annual species (Schenk and Jackson, 2002) and between highly-branched (rather than deep) root systems and drought-tolerance of grass species (Hartnett *et al*., 2013). However, it has been recently reported that *DEEPER ROOTING 1*, an auxin inducible gene of *Oryza sativa*, contributes to drought tolerance through vertically deeper RSA formation (Uga *et al*., 2013). In contrast to *A. thaliana*, which often grows in areas of highly variable rainfall, rice is cultivated at relatively high temperatures on irrigated soil. Therefore, the differences in RSA observed between these species might reflect contrasting strategies for adaptation to environments with different water-limitation constraints.

### GWA mapping can uncover novel players with adaptive relevance

Despite the fact that the auxin pathway is one of the most highly investigated signaling pathways in plants, we found no significant associations with loci in proximity of known auxin pathway components. While false negatives can be expected due to our approach (in particular the limited number of accessions as well as the stringent population structure control and statistical threshold), this still suggests that the common alleles mapped by GWA studies using phenotypic variation in natural populations are typically not the large-effect genes that are frequently mapped by mutant approaches. This is in agreement with other root GWA studies in which unbiased approaches were taken to map genes. For instance, despite the large body of work for nitrogen responses of roots based on mutant approaches, recent GWA studies of natural variation in nitrogen-level dependent root growth responses (Gifford *et al*., 2013) exclusively identified genes which had not been attributed to a function in this process based on past studies. The same pattern holds true for root growth regulation at the cellular scale (Meijon *et al*., 2014) and the organ level (Lachowiec *et al*., 2015; Slovak *et al*., 2014). Finally, a recent GWAS study showed that genes known for their involvement in the abscisic acid hormonal pathway were not enriched in the large number of genes associated with root growth responses to the hormone abscisic acid (Ristova *et al*., 2018). A reason for the absence of known genes could be that alterations of the expression or function of genes that are critically important for fundamental processes in development cause drastic effects with pleiotropic consequences, causing detrimental effects for the plants. Such alleles are likely not commonly maintained in a natural population. Therefore, with regard to understanding the genetic components that determine phenotypes (the genotype to phenotype challenge) and are of adaptive relevance, mapping of these natural alleles seems to be a highly promising approach. Overall, our findings not only demonstrate a specific role of exocytosis in the modulation of auxin-dependent developmental processes, but also underscore the importance of processes related to auxin pathway for the natural variation of RSA. The investigation of other exocytosis components promises to elucidate precise and dynamic control mechanisms of auxin dependent plant growth processes.

## EXPERIMENTAL PROCEDURES

### Plant materials and growth conditions

All plants for image acquisition on plates were grown on vertical 1 × MS agar plates (1 % sucrose, 0.8 % agar, pH = 5.7) in 12 cm × 12 cm square plates with or without chemicals under long day conditions (16 hours light) at 21°C. The seeds of the 215 natural accessions of *Arabidopsis thaliana* (Table S1) were a gift from Dr. Magnus Nordborg’s laboratory (GMI, Vienna, Austria). *EXO70A3ox* (SALK_075426) were purchased from Nottingham Arabidopsis Stock Center (Leicestershire, UK). Two CRISPR knockout lines (*CRISPR-exo70A3-1* and *CRISPR-exo70A3-2*) were created following a previously published method (Mao *et al*., 2016). *PIN1-GFP, PIN2-GFP, PIN3-GFP, PIN4-GFP, PIN7-GFP, pin4-3* and *DR5::GFP* were a gift from Dr. Jürgen Kleine-Vehn (University of Natural Resources and Life Sciences, Vienna, Austria). Crosses between *CRISPR-exo70A3-1*, *EXO70A3ox* and other mutants were made by manually pollinating emasculated pistils of those marker lines with pollen of these *EXO70A3* mutants. For allelic complementation lines, protein coding genetic sequences of *EXO70A3* were amplified from various natural accessions and fused with *EXO70A3* promoter sequence from Col-0, then transformed to *CRISPR-exo70A3-1*. Plants for crosses, seed amplification, and phenotyping of aerial parts were grown on soil at 21°C, 16 hours light/16°C, 8 hours dark conditions.

### Quantitative phenotyping of NPA treated roots and GWA study

The 215 natural accessions were grown on 1 × MS agar plates containing 10 μM NPA under long day conditions (16 hours light) at 21°C. 24 plants were seeded for each accession. Starting at 2 days after germination (DAG), root images were acquired by scanners (EPSON Perfection V600 Photo, Seiko Epson CO., Nagano, Japan) each 24 hours for 5 days (2 DAG – 6 DAG). Root image analyses and phenotype quantification were performed using the BRAT software (Slovak *et al*., 2014). GWA studies were conducted using the average trait value for each trait of 215 natural accessions by an accelerated mixed model (EMMAX (Kang *et al*., 2010)) followed by EMMA (Kang *et al*., 2008) for the most significant 200 associations as implemented in the web-based GWA mapping application GWAPP (Seren *et al*., 2012, conducted on December 15, 2014).

### Data mining and genetic analysis

The eFP browser (Winter *et al*., 2007) was used as a general reference for gene expression profiles of candidate genes. SNP data obtained from the *Arabidopsis thaliana* 1001 genome project (Schmitz *et al*., 2013, www.1001genomes.org) was used to analyze polymorphism among natural accessions. The analysis regarding signals of selection was conducted using the Selection Browser of the Regional Mapping Project (Horton *et al*., 2012). In haplogroup analysis, grouping of the 215 natural accessions of *A. thaliana* was performed according to haplotypes around the SNPs in a 40KB window around the SNPs of interest. These SNPs were used as the input for fastPHASE (Scheet and Stephens, 2006) version 1.4.0 and visualized using R script.

### Plant observations

For root phenotyping of *CRISPR-exo70A3-1, CRISPR-exo70A3-2, EXO70A3ox, 35S::GFP:EXO70A3, pin4-3* and *pin4-3 EXO70A3ox* young seedlings, seedlings were grown on 1 × MS agar plates. For root angle changes of these mutants or allelic complementation lines in response to NPA, plants grown for five days after germination were transferred to 1 × MS agar plates with or without 0.5 μM or 0.1 μM NPA, respectively, and images were acquired by scanners (EPSON Perfection V600 Photo) one day after the transfer and analyzed using Fiji (Schindelin *et al*., 2012). For each genotype, 10 or more plants were used. For the observation of statocytes, plants were grown on 1 × MS agar plates for five days after germination and stained in Lugol’s solution for two minutes and transferred into a chloral hydrate solution. Images were acquired using an AXIO image.M1 or an AXIO observer Z1 (Carl Zeiss AG, Oberkochen, Germany) and a CMOS camera (SPOT Idea 1.3 Mp Color Digital Camera, SPOT imaging, Sterling Heights, MI, USA). 5 - 10 plants per genotype were observed in an experiment. For analysis of above-ground organs of mature plants, images were acquired using a Nikon D80 (Nikon Co., Tokyo, Japan) digital camera. 4 - 9 plants were observed for each line. For analyses of fluorescent protein containing lines, crossbred lines and control lines for these crossbreds were grown on 1 × MS agar plates with or without 1 μM NPA for six days after germination. GFP signals in plant roots were observed under a confocal microscope Zeiss LSM700 (Carl Zeiss) with a 10× objective (EC plan Neofluar 10×0.3 M27), a 20× objective (Plan Apochromat 20×0.8 M27) or a 40× objective (Plan-Apochromat 40×/0.95 Korr M27, Zeiss). FRAP (Fluorescent recovery after photo bleaching) was conducted using *PIN4::PIN4:GFP* and *PIN4::PIN4:GFP EXO70A3ox* lines using a Visitron Spining Disc microscope, which is based on Nikon Eclipse Ti E inverted microscope (Nikon Instech Co., Ltd., Tokyo, Japan) equipped with CSU-W1 spinning disc unit (Yokogawa Electric Corporation, Tokyo, Japan), with a 60× water objective (Plan Apo VC 60×/1.2 WI water, Nikon) and Immersol WTM (Carl Zeiss). At least ten plants per genotype were observed for an experiment. To compare traits between haplogroups, we used eight accessions (haplogroup C: Ler-1, Shakdara (Sha), Lp2-2, and Per-1; haplogroup H: LAC-5, MIB-60, Coc-1, and Ty-0). All experiments were repeated at least twice.

### Root system architecture (RSA) assay

RSA in soil was studied using three plants from each genotype and in a blind fashion, so that the plant genotypes were not known to the researcher conducting the test. Each plant was grown on soil for 30 days in a pot (*ϕ* 20 cm, 17 cm height) at 21°C, 16 hours light/16°C, 8 hours dark conditions. The whole pot was then sectioned in two halves at the center of the plant and an RSA image was acquired by a Nikon D80 camera. Using Fiji, the roots on the image were marked (overlay) and subsequently the coordinates of all root pixels were determined. The Shallowness (standard deviation of all root pixels in X axis direction in a picture divided by that in Y axis direction) was calculated using these data as an indicator of shallow or deep RSA.

### Seed number assay

As a proxy of fitness in drought conditions, we measured seed number/plant for 8 natural accessions, Col-0 and *EXO70A3ox*. We used four plants from each genotype and four sets of the whole drought experiment. The experiment was conducted blindly so that the plant genotypes were not known to the researcher conducting the test. Four plants of a line were grown on soil for more than 60 days in a pot (*ϕ* 20 cm, 17 cm height) at 60% humidity, 21°C, 16 hours light/16°C, 8 hours dark conditions until seed production was completed. Four plants of two different genotypes were grown within one pot. In drought conditions, the water content of pots was decreased from 60% (Day After Germination: DAG5) to 30% (DAG 35). After that, the water content was kept at 25 %. For the water-sufficient control, the same experiment was repeated while water level was kept at 65% through the whole experimental period. Mature siliques from each plant were harvested and the number of seeds in five siliques was counted to determine the average number of seeds per silique for each plant. Subsequently, the total silique number of each plant was counted and the total seed number was calculated using the average seed number per silique and the total silique number. An average value of four plants of each line in a pot was calculated, rounded and used in statistical analyses. Significance of the difference in drought responses was calculated using Poisson regression models.

### Quantitative reverse transcription PCR (qrt-PCR)

For qrt-PCR, plants were grown on 1 × MS agar plates under normal growth conditions with or without 10 μM NPA or 100 nM IAA for 7 days after germination. Tissues were collection by excision with fine scissors. For root tips, 2.5 mm root tips were excised from primary roots. 10 plants for the whole plant, 10 plants for the shoot, 20 plants for the root, and more than 150 plants for the root tip samples were used. Samples were immediately frozen in liquid nitrogen, ground, and total RNA was extracted using the RNeasy Plant Mini kit (QIAGEN GmbH, Hilden, Germany). qrt-PCR reaction was prepared using 2x SensiMix™ SYBR & Fluorescein Kit (PEQLAB LLC, Wilmington, DE, USA) and PCR was conducted with a combination of iCycler and IQ5 (Bio-Rad Laboratories, Inc., Hercules, CA, USA). All transcript levels were normalized to the tubulin β gene (AT5G62690) or 18S ribosomal RNA. All experiments were repeated at least twice. Genes quantified include *EXO70A3* (AT5G52350), AT5G52340, AT5G52360, PIN1 (AT1G73590), PIN2 (AT5G57090), PIN3 (AT1G70940), PIN4 (AT2G01420), and PIN7 (AT1G23080). For the comparison of *EXO70A3* expression levels among natural accessions, we used eight accessions (haplogroup C: Ler-1, Shakdara (Sha), Lp2-2, and Per-1; haplogroup H: LAC-5, MIB-60, Coc-1, and Ty-0).

## Supporting information

All supplemental information

## ACKNOWLEDGEMENTS

The authors thank U. Seren, B. Vilhjalmsson, E. Sasaki, D. Meng, A. Korte, and M. Nordborg for assistance with the GWA and genetic analysis, and access to software. We are grateful to T. Tsuchimatsu, P. Zhang, and Matthew Horton for valuable discussions and T. Greb, P. Benfey, H. Tsukagoshi, J. Van Norman, and members of Busch laboratory for critical reading of the manuscript and T. Friese and M. Watson for manuscript editing. We thank V. Nizhynska, P. Korte, and M. Nordborg (Gregor Mendel Institute, Vienna, Austria) for donating seeds of natural accessions and M. Ruiz Rosquete and J. Kleine-Vehn (University of Natural Resources and Life Sciences, Vienna, Austria) for donating the GFP marker lines. We are thankful to the IMBA/IMP/GMI Workshop, in particular to M. Ziegler, D. Kummerer, and M. Colombini for their special efforts for the root system architecture assay. We thank VSCF for the creation of CRISPR lines. This research was supported by funds from the Austrian Academy of Sciences through the Gregor Mendel Institute, a grant from the Austrian Science Fund (FWF P27163-B22), and start-up funds from the Salk Institute for Biological Studies. T.O. and W.B. conceived the experiments. T.O. organized, performed, and analyzed all experiments. C.G. and W.B. wrote software for image acquisition and image analysis. D.F. developed software for the haplotype analysis and conducted population structure based statistical analyses. M.M. participated in phenotyping of plants. R.S. conducted the root phenotyping under control conditions. W.B. and S.S. participated in the root system architecture assay. T.O., D. F., and W.B. wrote the manuscript.

