## Supplementary material for "EXOCYST70A3 controls root system depth in Arabidopsis via the dynamic modulation of auxin transport": All supplemental information

**Figure & Table captions for Supplementary materials:**

**Table S1. Root traits of *A. thaliana* accessions on NPA and MS media, Related to Figure 1 and 2**

ID: Identifier of *A. thaliana* natural accessions as used in GWAPP (Seren et al., 2012); Accession name: common accessions name; Origin: country of collection; 10 µM NPA treatment: traits acquired from images of plants (6 DAG) grown on media (1 × MS media, 1% sucrose, 0.8% agar, 16 hours light conditions, 21 °C) containing 10 µM of auxin transport inhibitor NPA; Control: traits acquired from images of plants (5 DAG) grown on media (1 × MS media, 1% sucrose, 0.8% agar, 16 hours light conditions, 21 C°). Control data from (Slovak et al., 2014). Haplogroup: classification based on haplotypes around the *EXOCYST70A3* locus. C: Common-sensitivity to NPA; H: High-sensitivity to NPA.

**Table S2. Gene models in proximity to the significantly associated locus for Horizontality under NPA treatment, Related to Figure 1 and 2.**

Tair10 gene models in proximity (3 KB distance) to the significant SNP (passing the Bonferroni corrected threshold for approximately 215,000 tests, *p*-value < $10^{-7}$).

**Table. S3. Seed production under control (sufficient water) and drought conditions, Related to Figure 7**

Haplogroup: classification based on haplotypes around the *EXOCYST70A3* locus. C: Common-sensitivity to NPA; H: High-sensitivity to NPA.; Treatment: water conditions during plant growth. Control: 65% water/soil.; Drought: 25% water/soil.; Replicate: experimental replicates; GWAS ID: Identifier of *A. thaliana* natural accessions as used in GWAPP (*26*).

**Figure S1. Trait correlations of untreated and NPA treated roots of 215 *A. thaliana* natural accessions, Related to Figure 1**

(**A**) Histograms of data distribution for traits of NPA treated plants. x-axis: phenotype value; y-axis: Frequency. (**B**) z-scores of each root trait were compared between control and NPA treatments. x-axis: control; y-axis: 10 µM NPA treatment. *r* indicates Pearson's correlation coefficient between control and NPA treatment in relation to the phenotype. *p*-value shows the significance of these correlations.

**Figure S2. Analysis of the locus around the GWA-significant SNP, Related to Figure 1, 2 and 3**

(**A**) Two haplogroups that divide 215 natural accessions according to 40k bp sequences around the SNP that was found to be significantly associated with Horizontality by GWA mapping (Table S2). Haplotype structures were analyzed by fastPHASE (Scheet and Stephens, 2006) and visualized by blocks with different colors using 250K SNPs data from the Regional Mapping Project (Horton et al., 2012) with the dendrogram indicating the haplogroups. Trait (Horizontality, white circles) values and the nucleotides of the accessions at the SNP position (green and orange bars) are indicated. Each peripheral node in the top dendrogram shows a natural accession of *A. thaliana*. Black horizontal lines show the *EXO70A3* gene. Accessions between green dashed lines belong in haplogroup H and other accessions in haplogroup C.

(**B**) Gene model and polymorphisms among 30 natural accessions for the genomic region surrounding the significant GWA SNP. Red line: location of the significant GWA SNP. Top of the pannel: a magnification of the Manhattan plot for this genomic region. x-axis: chromosomal position; y-axis: significance (negative logarithmic of *p*-value); Black dashed line: the threshold of significance (Bonferroni-corrected *p*-value); Red dot: top SNP; Blue dots: other SNPs in the region. Middle of the panel: a gene model of this locus. arrows: gene; boxes: exon. Bottom of the panel: SNPs and amino acid substitutions in this locus in 30 accessions. Depicted are representative accessions of haplogroups C and H. Data from 1001 genome project (Schmitz et al., 2013, 1001genomes.org). Gray boxes: genomic locus equivalent to Col-0; Gray lines: SNP in non-coding region; Cyan lines: synonymous SNP; Magenta lines: non-synonymous SNP which is exclusively for Haplogroup H; Yellow lines: other non-synonymous SNPs. (**C**) Gene models of natural accessions that were used in *EXO70A3* allelic complementation analysis. Orange arrow: genomic sequence of Col-0 *EXO70A3* coding region; Light green boxes: exon; Gray bar: intergenic region; Black bars: genomic locus of various natural accessions that are equivalent to Col-0 *EXO70A3*; Gray lines: SNP in non-coding region; Cyan lines: synonymous SNP; Magenta lines: non-synonymous SNP which is exclusively for Haplogroup H; Yellow lines: other non-synonymous SNPs.

**Figure S3. Gene expression patterns of *EXO70A3* and neighboring genes, Related to Figure 2, 4 and 5**

(**A-D**) Tissue-, cell-type specific, and auxin related treatment-dependent expression pattern of *EXO70A3* gene from publicly available microarray datasets visualized by the Arabidopsis eFP browser ((Winter et al., 2007), http://bar.utoronto.ca/efp/cgi-bin/efpWeb.cgi). Signals less than or equal to 20 (the background value of AtGEN Express (Goda et al., 2008; Schmid et al., 2005)) were masked with a grey color. Signals above this threshold were color coded in relation to their average levels in the respective datasets (see color legend for each data set). (A) AtGen Express data (organ level). (B) Cell-type specific RootMap (cellular level). (C, D) IAA and NPA treatments (whole seedlings). (**E**) *EXO70A3* expression profile in various root cells and tissues which was quantified by RNA-Seq (data collected from Li *et al.*, 2017). x-axis: cells/tissues; y-axis: transcript level (Fragments per kilo base per million mapped reads: FPKM). (**F-I**) qRT-PCR results of expression levels of *EXO70A3* and related genes. Transcript levels of proximal genes of *EXO70A3*, AT5G52340 (F) and AT5G52360 (G), in various organs of Col-0 and *EXO70A3ox*. x-axis: sample; y-axis: relative expression level (normalized to Col-0 Whole); Error bars: standard deviation. (H) Gene expression levels of five *PIN* and *EXO70A3* genes in root tip of *EXO70A3ox*. x-axis: gene; y-axis: transcript level (normalized to Col-0); error bars: standard deviation. Asterisk: significance evaluated by Welch t-test ($p$-value = $2.2 \times 10^{-3}$, n = 6). (I) Expression levels of *EXO70A3* in root tips of 4 natural accessions of the haplogroups C and H, Col-0, and *EXO70A3ox* lines. Expression levels normalized to Col-0. x-axis: sample; y-axis: normalized expression level; Error bars: standard deviation; Asterisk: significance evaluated by Student t-test ($p$-value = 0.014). Haplogroups C and H were not distinctive (ANOVA > 0.05).

**Figure S4. PIN:GFP localization in *EXO70A3ox/PIN::PIN:GFP* lines, Related to Figure 4**

Confocal images of representative roots grown on MS media. Green: GFP signal; Red: cell wall stained with propidium iodide (PI); Yellow bars: 100 μm. (**A**) Median longitudinal optical sections of root tips (5 DAG) of *PIN1::PIN1:GFP*, *PIN2::PIN2:GFP*, *PIN3::PIN3:GFP*, and *PIN7::PIN7:GFP* lines and crosses with the *EXO70A3ox* line are shown. (**B**) Confocal images of representative lateral roots (15 DAG) of

*PIN4::PIN4-GFP* and a cross with *EXO70A3ox* line. Upper panels: overview of low resolution images of emerging lateral roots. Boxes indicate the position of high-resolution images depicted below. Lower panel: median longitudinal optical sections of lateral root tips.

**Figure S5. Phenotypic analysis of *EXO70A3*-related lines, Related to Figure 4 and 5**

**(A-E)** Various auxin-related traits including Statocytes and statolith, IAA responses in primary roots and hypocotyl, lateral root density, shoot gravitropism, rosette development, phyllotaxy, floral stem height and inflorescence structure in *EXO70A3*-related lines. *CRISPR-exo70A3-1* (A), *CRISPR-exo70A3-2* (B), *EXO70A3ox* (C), *35S::GFP:EXO70A3* (D) and *pin4-3* (E) were compared with Col-0. Statocytes and statolith: representative root tips of plants (5 DAG) grown on MS plates as visualized by Lugol staining. Black bars: 20 µm. IAA response in primary root: Plants were treated with 0.1, 1, 10, or 100 nM IAA on 1 × MS plates and pictured at 5 DAG. x-axis: treatments; y-axis: primary root length; Error bars: standard deviation; "a" and "b": two significantly different groups (evaluated by post-hoc Tukey test after ANOVA, $p$-value < 0.05). Hypocotyl elongation in response to IAA: Plants were treated with 10 or 100 nM IAA on 1 × MS plates and pictured at 5 DAG. x-axis: treatments; y-axis: hypocotyl length; Error bars: standard deviation; "a" and "b": two significantly different groups (evaluated by post-hoc Tukey test after ANOVA, $p$-value < 0.05). Lateral root density: representative lateral root structures of plants. Plants were grown on 1 × MS plates until 5 DAG and equal size 5 plants were transferred to fresh 1 × MS plate for further growth, then pictured at 20 DAG. Yellow bars: 5 cm; x-axis: average lateral root density [number/cm primary root]; y-axis: genotype; Error bars: standard deviation. Shoot gravitropism: inflorescence angle changes during shoot gravitropism response of 40 DAG plants. x-axis: time after rotation [h]; y-axis: average angle change [º]; Error bars: standard deviation. Rosette development, phyllotaxy, floral stem height and floral structure: Mutant plants were grown with Col-0 plants on soil side by side under long day conditions (16 hours light, 21 °C/8 hours dark, 16 °C) and pictured at 15 and 30 DAG (rosette development and phyllotaxy), 45 DAG (inflorescence structure) and 50 DAG (floral stem height). Yellow bars: 5 cm; x-axis: genotype; y-axis: phenotypes (indicated in charts); Error bars: standard deviation; WT: Col-0; Mutant: mutant line which is indicated in the title of each figure. NPA response (E): Col-0 and *pin4-3* plants were grown on 1 × MS plates until 5 DAG and equal size plants were transferred to chemical treatment plates (mock and 0.5 µM NPA), then pictured (1 day after transfer). Beanplot: x-axis: treatment; y-axis: root tip angle [º]; Red lines: mean; short lines: individual data points; shape: density of the data distribution. (**F**) Variances of root tip angles during dynamic root re-orientation of *pin4-3* and *pin4-3 EXO70A3ox*. Significance of the difference of variances between Col-0 and mutant was analyzed by Fisher test and color coded. x-axis: time [hours], y-axis: standard deviation of root tip angles. (**F**) Average angles and variances of root tips during dynamic root re-orientation of *EXO70A3*-related lines and *pin4-3*-related lines. x-axis shows genotype. Top panel: y-axis shows average root tip angle (6 hour after plate re-orientation), black point: genotype which showed significant phenotype in comparison with Col-0; Middle panel: y-axis shows root tip angle variation (10 hour after plate re-orientation), black point: genotype which showed significant phenotype in comparison with Col-0; Bottom panel: Rose plots to show root tip angle distribution (10 hour after plate re-orientation). black bar: occupancy (%) of roots in the range of direction in all roots (distance from center of circles shows the occupancy level); *pin4-3 EXO70A3ox* showed *pin4-3*-type phenotypes (no delay of root bending and the larger variation of root tip angles) but not *EXO70A3* mutants-type phenotypes (the delay of root bending and the larger variation of root tip angles), indicating that

*EXO70A3* is upstream of *PIN4*.

**Figure S6. Exploration of signals of selection in *EXO70A3* locus, Related to Figure 7**

**(A)** Selection signals around the significantly associated SNP as shown in the Selection Browser (Horton et al., 2012). Orange line: the position of the significant GWAS SNP; Red arrow: the closest significant selection signal ($p$-value < 0.01). (**B**) Distribution of associations of SNPs with precipitation seasonality that were reported in GWAS for climate adaptation (Hancock et al., 2011). x-axis: significance of association (Spearman's correlation coefficient: ρ). y-axis: frequency. Area between the blue lines shows the range which contains 95% of SNPs. Red line: Correlation of the significant SNP identified by our GWA mapping (chromosome 5: 21256465) which is located outside of the 95% range.

**Figure S7. Hypothetical multilevel model of *EXO70A3* function and effects on variations in traits, Related to Figure 6 and 7**

Model of EXO70A3 function based on the presented data. Lower left: Through the vesicle tethering function of EXO70A3 which is specific to vesicles that contain PIN4, PIN4 is delivered to the plasma membrane. Upper left: This occurs asymmetrically upon gravity stimulus in a subset of columella cells that are localized at the side of the root that points in the direction of gravity, leading to an auxin flux to the root side located towards the gravity direction and therefore establishes an asymmetric auxin distribution. Allelic variation of *EXO70A3* leads to altered EXO70A3 activity and determines the threshold and dynamics of this auxin flux resulting in distinct root growth direction control properties and lateral root density patterns among *A. thaliana* natural accessions, potentially involved in adaptation to various environments including water or nutrient deprived areas.

**movie S1. Dynamic root gravitropism responses of Col-0, *CRISPR-exo70A3-1* and EXO70A3ox after 90 degree re-orientation of roots, Related to Figure 5**

Plants grown for 5 days after germination on vertical 1 × MS agar plate under long day conditions (16 hour light) at 21 °C were transferred to another 1 × MS agar plate and incubated for 1 hours under the same conditions. Then the plate was turned 90 degrees and time dependent re-orientation of root tips to gravity was captured every 4 minutes using scanners for 20 hours. The movie is a time-lapse of these images and is a mosaic assembled from 3 representative individuals for each genotype. Upper row: Col-0 wildtype plants; Middle row: *EXO70A3ox* plants; lower row: *CRISPR-exo70A3-1* plants.

**movie S2. Dynamic root gravitropism responses of different *EXO70A3* allele accessions (haplogroups C and H) after 90 degree re-orientation of roots, Related to Figure 5**

Plants grown for 5 days after germination on vertical 1 × MS agar plate under long day conditions (16 hour light) at 21 °C were transferred to another 1 × MS agar plate and incubated for 1 hours under the same conditions. Then the plate was turned 90 degrees and time dependent re-orientation of root tips to gravity was captured every 4 minutes using scanners for 20 hours. The movie is a time-lapse of these images and is a mosaic assembled from 1 representative individual for each accession.

**Table S1. Root traits of *A. thaliana* accessions on NPA and MS media[a], Related to Figure 1 and 2**

ID: Identifier of *A. thaliana* natural accessions as used in GWAPP (Seren et al., 2012); Accession name: common accessions name; Origin: country of collection; 10 µM NPA treatment: traits acquired from images of plants (6 DAG) grown on media (1 × MS media, 1% sucrose, 0.8% agar, 16 hours light conditions, 21 °C) containing 10 µM of auxin transport inhibitor NPA; Control: traits acquired from images of plants (5 DAG) grown on media (1 × MS media, 1% sucrose, 0.8% agar, 16 hours light conditions, 21 °C). Control data from (Slovak et al., 2014). Haplogroup: classification based on haplotypes around the *EXOCYST70A3* locus. C: Common-sensitivity to NPA; H: High-sensitivity to NPA.

| | | | 10 µM NPA treatment | | | | Control | | | | Haplogroup |
|---|---|---|---|---|---|---|---|---|---|---|---|
| | | | Length traits | Angle traits | | Width traits | Length traits | Angle traits | | Width traits | |
| **GWAS_ID** | **Accession Name** | **Origin** | Total length | Horizontality | Skewness[b] | Average root width | Total length | Horizontality | Skewness[b] | Average root width | (*EXO70A3* locus) |
| 86 | CUR-8 | France | 240.00 | 81.82 | 57.20 | 7.69 | 713.69 | 116.04 | -4.49 | 4.07 | C |
| 96 | LAC-5 | France | 394.50 | 309.69 | 57.66 | 8.02 | 619.23 | 70.45 | 2.37 | 4.21 | H |
| 149 | LDV-58 | France | 125.00 | 106.37 | 58.12 | 8.26 | 650.11 | 160.30 | -6.63 | 4.79 | C |
| 204 | MIB-60 | France | 259.00 | 418.65 | 57.98 | 8.03 | ND | ND | ND | ND | H |
| 224 | MIB-86 | France | 263.00 | 101.16 | 58.09 | 7.32 | ND | ND | ND | ND | H |
| 236 | MOG-11 | France | 141.00 | 85.50 | 58.10 | 8.54 | ND | ND | ND | ND | C |
| 260 | PAR-5 | France | 241.00 | 159.01 | 58.17 | 8.13 | ND | ND | ND | ND | H |
| 266 | RAN | France | 237.00 | 196.73 | 58.63 | 8.12 | ND | ND | ND | ND | C |
| 394 | Vou-5 | France | 219.50 | 71.89 | 57.78 | 7.08 | ND | ND | ND | ND | C |
| 461 | EM-183 | UK | 123.00 | 49.82 | 57.68 | 7.47 | 484.78 | 55.10 | -3.98 | 3.94 | C |
| 936 | FOR-5 | USA | 72.00 | 90.24 | 57.25 | NA | 505.25 | 45.33 | -2.99 | 3.98 | C |
| 1874 | MNF-Pot-80 | USA | 302.00 | 179.09 | 57.65 | 7.95 | 677.08 | 59.58 | 2.24 | 3.94 | C |
| 2057 | Map-42 | USA | 261.00 | 120.12 | 57.71 | 7.86 | 733.46 | 74.27 | -0.75 | 4.32 | C |
| 2171 | Paw-26 | USA | 303.00 | 159.59 | 58.26 | 7.67 | ND | ND | ND | ND | C |
| 2290 | Ste-3 | USA | 231.50 | 135.80 | 57.50 | 7.27 | ND | ND | ND | ND | C |
| 2320 | Wilcox-4 | USA | 309.50 | 55.07 | 57.84 | 6.75 | ND | ND | ND | ND | C |
| 5719 | Bur-0 | Ireland | 294.00 | 66.06 | 56.76 | 8.08 | 963.52 | 307.15 | -9.02 | 4.90 | C |
| 5723 | Chr-1 | UK | 17.50 | 53.20 | 57.25 | NA | 305.51 | 52.61 | 3.71 | 4.04 | C |
| 5729 | Coc-1 | UK | 194.00 | 368.71 | 57.77 | 7.76 | 497.95 | 33.21 | -1.24 | 4.32 | H |
| 5731 | Crl-1 | UK | 280.00 | 68.90 | 57.71 | 7.27 | 650.44 | 78.34 | -3.74 | 4.10 | C |
| 5736 | Ema-1 | UK | 324.00 | 237.27 | 45.74 | 7.93 | 736.29 | 56.40 | -1.32 | 4.44 | C |
| 5742 | Frd-1 | UK | 215.50 | 130.35 | 57.45 | 7.70 | 643.36 | 83.76 | 0.24 | 3.67 | H |
| 5745 | Hil-1 | UK | 246.00 | 86.17 | 48.23 | 7.10 | 703.56 | 164.80 | -6.06 | 4.35 | C |
| 5751 | Kyl-1 | UK | 123.00 | 91.49 | 58.20 | NA | 632.51 | 53.46 | 0.28 | 4.18 | C |
| 5752 | Lan-1 | UK | 258.00 | 154.01 | 57.88 | 7.67 | 701.96 | 138.87 | -6.76 | 4.30 | C |
| 5837 | Bor-1 | Czech | 341.00 | 155.98 | 56.95 | 7.68 | 736.86 | 95.06 | 3.63 | 4.02 | C |
| 6008 | Duk | Czech | 263.00 | 114.94 | 47.58 | 7.34 | 691.98 | 72.97 | -4.03 | 4.02 | C |
| 6074 | Ör-1 | Sweden | 167.00 | 129.89 | 57.83 | 7.58 | 524.12 | 81.34 | -1.54 | 3.79 | C |
| 6243 | Tottarp-2 | Sweden | 233.50 | 124.53 | 57.47 | 7.31 | ND | ND | ND | ND | C |
| 6730 | CIBC-5 | UK | 210.00 | 56.77 | 57.21 | 8.12 | 576.78 | 150.21 | -12.35 | 4.42 | C |
| 6897 | Ag-0 | France | 249.00 | 130.12 | 57.52 | 7.59 | ND | ND | ND | ND | C |
| 6898 | An-1 | Belgium | 275.50 | 121.24 | 51.06 | 7.83 | 887.99 | 151.42 | -3.33 | 4.03 | C |
| 6899 | Bay-0 | Germany | 304.00 | 104.38 | 55.95 | 6.95 | 777.79 | 74.45 | -4.93 | 4.12 | C |
| 6903 | Bor-4 | Czech | 296.00 | 252.80 | 56.97 | 7.89 | 788.25 | 78.98 | -2.61 | 4.26 | C |
| 6904 | Br-0 | Czech | 191.00 | 71.94 | 57.28 | 8.86 | 786.50 | 52.92 | -1.45 | 4.24 | C |
| 6906 | C24 | Portugal | 208.00 | 109.57 | 57.09 | 8.19 | 614.46 | 83.09 | -3.70 | 4.14 | C |
| 6907 | CIBC-17 | UK | 257.50 | 343.27 | 57.19 | 8.62 | 672.07 | 125.16 | -4.63 | 4.39 | C |
| 6909 | Col-0 | USA | 340.50 | 160.68 | 57.63 | 6.99 | 741.51 | 48.11 | -1.72 | 3.76 | C |
| 6910 | Ct-1 | Itary | 230.00 | 134.62 | 57.70 | 6.96 | 683.83 | 70.82 | -5.15 | 4.01 | C |
| 6911 | Cvi-0 | Cape Verde | 202.00 | 193.77 | 57.90 | 7.51 | 860.06 | 352.44 | -6.75 | 4.74 | C |
| 6915 | Ei-2 | Germany | 311.50 | 186.85 | 57.47 | NA | 721.11 | 40.90 | -3.16 | 4.09 | C |
| 6916 | Est-1 | Russia | 342.00 | 342.86 | 49.21 | 7.69 | 805.90 | 161.72 | -4.46 | 4.23 | C |
| 6919 | Ga-0 | Germany | 359.00 | 235.26 | 57.20 | 7.84 | 744.24 | 251.23 | -13.66 | 4.01 | C |
| 6920 | Got-22 | Germany | 265.00 | 275.36 | 57.58 | 8.14 | 664.33 | 121.37 | -6.01 | 3.75 | H |
| 6922 | Gu-0 | Germany | 297.00 | 183.39 | 57.27 | 8.01 | 559.92 | 56.60 | -2.30 | 4.12 | C |
| 6923 | HR-10 | UK | 13.00 | 2.00 | 57.25 | NA | 614.31 | 61.44 | -1.97 | 4.35 | C |
| 6926 | Kin-0 | USA | 21.00 | 20.34 | 57.25 | NA | 504.88 | 50.99 | 0.72 | 4.08 | H |
| 6928 | Kno-18 | USA | 230.50 | 118.28 | 57.50 | 7.44 | 804.14 | 157.42 | -3.22 | 4.23 | C |
| 6929 | Kondara | Tajikistan | 57.00 | 20.54 | 57.28 | 8.56 | 662.68 | 97.95 | -5.15 | 4.20 | C |
| 6930 | Kz-1 | Kazakhstan | 351.00 | 107.75 | 57.44 | 7.19 | 862.87 | 172.36 | -6.40 | 4.43 | C |
| 6931 | Kz-9 | Kazakhstan | 209.00 | 1059.35 | 57.43 | 7.98 | 757.69 | 108.73 | -3.32 | 4.44 | C |
| 6932 | Ler-1 | Germany | 165.50 | 92.13 | 57.41 | 8.40 | 793.83 | 192.32 | -9.62 | 4.01 | C |
| 6933 | LL-0 | Spain | 275.00 | 117.04 | 57.35 | 7.60 | 502.61 | 81.00 | -4.30 | 3.73 | C |
| 6936 | Lz-0 | France | 375.00 | 106.63 | 57.23 | 7.62 | ND | ND | ND | ND | C |
| 6937 | Mrk-0 | Germany | 329.50 | 190.76 | 57.67 | 7.88 | 855.12 | 86.75 | 0.27 | 4.15 | C |
| 6938 | Ms-0 | Russia | 213.00 | 121.80 | 57.24 | 8.05 | ND | ND | ND | ND | C |
| 6939 | Mt-0 | Libya | 256.50 | 89.84 | 57.28 | 8.66 | ND | ND | ND | ND | C |
| 6940 | Mz-0 | Germany | 253.00 | 186.07 | 57.25 | 8.41 | ND | ND | ND | ND | C |
| 6942 | Nd-1 | Switzerland | 342.00 | 206.23 | 57.32 | 8.63 | ND | ND | ND | ND | C |

| | | | | | | | | | | | |
|---|---|---|---|---|---|---|---|---|---|---|---|
| 6943 | NFA-10 | UK | 179.00 | 95.88 | 57.27 | 8.59 | ND | ND | ND | ND | C |
| 6944 | NFA-8 | UK | 148.00 | 136.46 | 58.09 | 7.26 | ND | ND | ND | ND | C |
| 6945 | Nok-3 | Netherlands | 229.00 | 209.43 | 57.79 | 7.70 | 449.52 | 35.37 | 1.58 | 3.97 | C |
| 6946 | Oy-0 | Norway | 324.00 | 139.61 | 57.25 | 8.31 | ND | ND | ND | ND | H |
| 6951 | Pu2-23 | Czech | 212.50 | 127.99 | 56.85 | 7.72 | ND | ND | ND | ND | C |
| 6956 | Pu2-7 | Czech | 195.00 | 124.97 | 56.88 | 7.78 | ND | ND | ND | ND | C |
| 6958 | Ra-0 | France | 433.50 | 173.81 | 56.81 | 8.07 | ND | ND | ND | ND | C |
| 6959 | Ren-1 | France | 281.00 | 279.23 | 56.78 | 8.09 | ND | ND | ND | ND | H |
| 6960 | Ren-11 | France | 368.00 | 506.50 | 56.78 | 8.30 | ND | ND | ND | ND | H |
| 6961 | Se-0 | Spain | 267.00 | 114.71 | 57.47 | 6.65 | ND | ND | ND | ND | C |
| 6962 | Sha | Tajikistan | 277.00 | 98.35 | 57.49 | 7.54 | ND | ND | ND | ND | C |
| 6963 | Sorbo | Tajikistan | 148.00 | 119.33 | 57.45 | 8.15 | ND | ND | ND | ND | C |
| 6966 | Sq-1 | UK | 203.00 | 121.44 | 57.42 | 7.92 | ND | ND | ND | ND | C |
| 6967 | Sq-8 | UK | 327.00 | 221.05 | 57.50 | 7.25 | ND | ND | ND | ND | C |
| 6968 | Tamm-2 | Finland | 242.00 | 141.16 | 57.81 | 7.53 | 710.58 | 104.49 | -4.28 | 4.20 | C |
| 6969 | Tamm-27 | Finland | 242.00 | 133.31 | 57.81 | 7.80 | 804.65 | 91.11 | 2.56 | 4.10 | C |
| 6970 | Ts-1 | Spain | 268.00 | 83.22 | 57.50 | 6.77 | ND | ND | ND | ND | C |
| 6971 | Ts-5 | Spain | 241.50 | 116.50 | 57.86 | 6.69 | 613.48 | 107.25 | -4.83 | 3.88 | C |
| 6972 | Tsu-1 | Japan | 453.00 | 207.46 | 57.49 | 8.15 | ND | ND | ND | ND | C |
| 6973 | Ull2-3 | Sweden | 270.00 | 171.43 | 57.77 | 8.18 | ND | ND | ND | ND | C |
| 6975 | Uod-1 | Austria | 260.50 | 153.57 | 57.74 | 7.32 | ND | ND | ND | ND | C |
| 6976 | Uod-7 | Austria | 338.00 | 141.75 | 57.80 | 7.05 | 745.19 | 44.78 | 1.76 | 3.89 | C |
| 6977 | Van-0 | Canada | 202.50 | 85.93 | 57.76 | 7.62 | ND | ND | ND | ND | C |
| 6978 | Wa-1 | Poland | 260.00 | 164.83 | 57.61 | 8.46 | ND | ND | ND | ND | C |
| 6979 | Wei-0 | Switzerland | 279.50 | 286.86 | 57.73 | 7.40 | ND | ND | ND | ND | C |
| 6980 | Ws-0 | Russia | 376.00 | 235.58 | 57.75 | 8.15 | ND | ND | ND | ND | C |
| 6981 | Ws-2 | Russia | 197.00 | 69.04 | 57.80 | 7.29 | 623.98 | 78.34 | -3.89 | 3.72 | C |
| 6982 | Wt-5 | Germany | 338.00 | 271.95 | 57.76 | 7.27 | ND | ND | ND | ND | C |
| 6984 | Zdr-1 | Czech | 256.00 | 93.36 | 57.40 | 6.91 | ND | ND | ND | ND | C |
| 6985 | Zdr-6 | Czech | 192.00 | 147.94 | 57.40 | 7.40 | ND | ND | ND | ND | C |
| 6987 | Ak-1 | Germany | 227.00 | 91.75 | 52.46 | 7.23 | 524.12 | 100.35 | -5.47 | 3.97 | C |
| 6990 | Amel-1 | Netherlands | 428.00 | 167.37 | 56.98 | 7.85 | 900.22 | 137.70 | 0.82 | 4.41 | C |
| 6992 | Ang-0 | Belgium | 251.00 | 151.11 | 51.41 | 6.93 | 637.17 | 103.64 | 0.80 | 3.79 | C |
| 6994 | Ann-1 | France | 212.00 | 158.95 | 57.48 | 8.06 | ND | ND | ND | ND | C |
| 7000 | Aa-0 | Germany | 344.00 | 214.82 | 51.10 | 7.95 | 700.92 | 71.99 | -1.64 | 4.32 | C |
| 7002 | Baa-1 | Netherlands | 253.00 | 606.71 | 50.56 | 8.18 | 463.09 | 38.30 | 4.65 | 4.20 | C |
| 7004 | Bs-2 | Switzerland | 310.50 | 193.10 | 56.08 | 7.61 | 862.03 | 100.65 | -4.66 | 3.87 | C |
| 7014 | Ba-1 | UK | 184.00 | 153.83 | 57.04 | 8.70 | 535.61 | 63.22 | -2.68 | 4.13 | C |
| 7015 | Bla-1 | Spain | 201.00 | 100.80 | 57.25 | NA | 745.21 | 60.33 | -2.28 | 4.01 | C |
| 7026 | Boot-1 | UK | 291.00 | 221.85 | 56.41 | 8.23 | 677.96 | 63.44 | -1.07 | 4.16 | H |
| 7031 | Bsch-0 | Germany | 218.00 | 181.73 | 55.94 | 7.80 | 784.06 | 89.08 | -2.97 | 4.13 | C |
| 7062 | Ca-0 | Germany | 343.00 | 368.35 | 56.04 | 8.02 | 888.75 | 113.22 | -2.99 | 4.34 | C |
| 7071 | Chat-1 | France | 320.00 | 181.70 | 55.97 | 7.54 | 633.74 | 63.08 | -0.29 | 3.98 | C |
| 7081 | Co | Portugal | 476.00 | 382.83 | 57.66 | 7.69 | 1066.93 | 144.97 | -5.62 | 4.58 | C |
| 7092 | Com-1 | France | 260.50 | 101.00 | 57.86 | 7.00 | 613.26 | 133.03 | -6.70 | 4.25 | C |
| 7094 | Da-0 | Germany | 208.50 | 146.59 | 57.60 | 6.99 | 655.74 | 67.80 | -5.74 | 3.81 | C |
| 7098 | Di-1 | France | 291.50 | 128.08 | 47.43 | 7.09 | 742.22 | 180.49 | -7.00 | 4.24 | C |
| 7102 | Do-0 | Germany | 282.00 | 94.90 | 47.30 | 6.90 | 717.56 | 53.51 | -1.09 | 4.22 | C |
| 7123 | Ep-0 | Germany | 227.00 | 102.05 | 50.30 | 7.29 | 543.55 | 62.24 | -4.67 | 3.58 | C |
| 7143 | Gel-1 | Netherlands | 360.00 | 143.63 | 50.49 | 7.40 | 776.15 | 61.73 | 2.50 | 4.24 | C |
| 7147 | Gie-0 | Germany | 310.00 | 277.21 | 57.28 | 7.88 | 840.00 | 190.18 | -4.95 | 3.85 | C |
| 7163 | Ha-0 | Germany | 274.00 | 166.59 | 49.15 | 7.17 | 718.88 | 100.83 | -6.25 | 4.46 | C |
| 7166 | Hey-1 | Netherlands | 232.00 | 102.46 | 54.66 | 6.86 | 790.97 | 77.57 | -1.83 | 4.17 | C |
| 7172 | Hl-3 | Germany | 310.00 | 241.92 | 48.05 | 7.78 | 679.64 | 85.74 | -4.57 | 4.70 | C |
| 7176 | Is-1 | Germany | 249.50 | 141.84 | 57.97 | 7.05 | ND | ND | ND | ND | C |
| 7178 | Jm-1 | Czech | 348.00 | 159.75 | 58.24 | 8.05 | 857.07 | 69.41 | -1.78 | 4.68 | C |
| 7181 | Je-0 | Germany | 215.00 | 127.80 | 57.25 | 8.79 | 593.43 | 60.74 | 0.81 | 4.52 | C |
| 7192 | Kil-0 | UK | 284.50 | 231.59 | 58.08 | 8.19 | 890.60 | 193.84 | -5.87 | 4.45 | C |
| 7199 | Kl-5 | Germany | 313.50 | 164.14 | 58.02 | 7.35 | 728.28 | 230.72 | -11.62 | 4.23 | C |
| 7201 | Kr-0 | Germany | 347.00 | 125.13 | 58.20 | 7.12 | 762.80 | 101.52 | -3.12 | 4.32 | C |
| 7205 | Krot-2 | Germany | 285.00 | 81.88 | 58.07 | 7.21 | 772.04 | 81.66 | -1.52 | 4.23 | C |
| 7210 | La-1 | Poland | 334.00 | 141.09 | 57.42 | 7.81 | 877.06 | 69.03 | 2.37 | 4.34 | C |
| 7224 | Li-3 | Germany | 528.00 | 226.80 | 57.64 | 8.00 | 1007.75 | 88.92 | -2.48 | 4.25 | C |
| 7231 | Li-7 | Germany | 436.00 | 337.58 | 57.67 | 7.65 | 743.78 | 89.73 | -1.82 | 3.89 | C |
| 7242 | Lo-2 | Germany | 275.00 | 172.12 | 58.10 | 7.70 | 806.71 | 79.19 | -2.33 | 4.03 | C |
| 7244 | Mnz-0 | Germany | 287.00 | 192.29 | 58.00 | 7.72 | ND | ND | ND | ND | C |
| 7246 | Ma-2 | Germany | 265.50 | 413.39 | 58.01 | 7.76 | ND | ND | ND | ND | C |
| 7255 | Mh-0 | Poland | 287.00 | 181.61 | 58.05 | 7.82 | ND | ND | ND | ND | C |

| | | | | | | | | | | | |
|---|---|---|---|---|---|---|---|---|---|---|---|
| 7262 | Nw-4 | Germany | 312.00 | 130.95 | 58.02 | 7.84 | ND | ND | ND | ND | C |
| 7268 | Np-0 | Germany | 281.00 | 211.16 | 58.04 | 7.45 | ND | ND | ND | ND | H |
| 7275 | No-0 | Germany | 254.50 | 90.96 | 57.99 | 7.13 | ND | ND | ND | ND | C |
| 7276 | Ob-0 | Germany | 261.00 | 153.66 | 58.08 | 7.11 | ND | ND | ND | ND | C |
| 7280 | Old-1 | Germany | 316.00 | 118.56 | 58.22 | 7.80 | ND | ND | ND | ND | C |
| 7282 | Or-0 | Germany | 260.50 | 108.39 | 58.27 | 7.54 | ND | ND | ND | ND | C |
| 7283 | Ors-1 | Romania | 253.00 | 111.04 | 57.82 | 7.44 | 621.58 | 87.11 | -3.08 | 3.87 | C |
| 7287 | Ove-0 | Germany | 382.00 | 193.04 | 58.28 | 8.06 | ND | ND | ND | ND | C |
| 7291 | Pa-2 | Itary | 218.00 | 118.39 | 58.21 | 7.57 | ND | ND | ND | ND | C |
| 7297 | Pf-0 | Germany | 274.50 | 97.28 | 58.19 | 6.95 | ND | ND | ND | ND | C |
| 7299 | Pi-2 | Austria | 274.00 | 94.06 | 58.79 | 7.13 | ND | ND | ND | ND | C |
| 7300 | Pla-0 | Spain | 354.00 | 133.19 | 56.83 | 7.78 | ND | ND | ND | ND | C |
| 7306 | Pog-0 | Canada | 306.50 | 113.51 | 58.81 | 7.93 | ND | ND | ND | ND | C |
| 7307 | Pn-0 | France | 206.00 | 76.51 | 58.47 | 8.39 | ND | ND | ND | ND | C |
| 7309 | Po-1 | Germany | 284.50 | 133.28 | 56.79 | 7.93 | ND | ND | ND | ND | C |
| 7310 | Pr-0 | Germany | 227.00 | 117.43 | 58.75 | 7.50 | ND | ND | ND | ND | C |
| 7316 | Rhen-1 | Netherlands | 308.00 | 150.75 | 57.81 | 7.80 | 790.89 | 126.45 | -3.18 | 3.98 | C |
| 7317 | Ri-0 | Canada | 69.00 | 89.59 | 58.16 | 8.23 | ND | ND | ND | ND | H |
| 7320 | Rou-0 | France | 218.00 | 119.92 | 58.74 | 7.70 | ND | ND | ND | ND | C |
| 7330 | Sapporo-0 | Japan | 239.00 | 158.57 | 57.48 | 7.28 | ND | ND | ND | ND | C |
| 7331 | Sh-0 | Germany | 349.00 | 137.93 | 57.56 | 6.88 | ND | ND | ND | ND | C |
| 7337 | Si-0 | Germany | 293.00 | 175.70 | 57.50 | 7.30 | ND | ND | ND | ND | C |
| 7344 | Sg-1 | Germany | 273.00 | 246.39 | 57.55 | 7.68 | ND | ND | ND | ND | C |
| 7351 | Ty-0 | UK | 233.00 | 305.17 | 57.83 | 8.20 | ND | ND | ND | ND | H |
| 7352 | Te-0 | Finland | 165.00 | 185.87 | 57.48 | 7.74 | ND | ND | ND | ND | C |
| 7353 | Tha-1 | Netherlands | 341.50 | 258.24 | 57.47 | 7.56 | ND | ND | ND | ND | C |
| 7372 | Tscha-1 | Austria | 331.50 | 240.36 | 57.49 | 7.91 | ND | ND | ND | ND | C |
| 7378 | Uk-1 | Germany | 117.00 | 99.45 | 57.83 | 8.23 | ND | ND | ND | ND | C |
| 7382 | Utrecht | Netherlands | 378.00 | 123.48 | 57.79 | 6.94 | ND | ND | ND | ND | C |
| 7384 | Ven-1 | Netherlands | 303.00 | 213.22 | 57.88 | 7.99 | 835.70 | 188.58 | -7.71 | 4.02 | C |
| 7404 | Wc-1 | Germany | 304.50 | 132.72 | 57.79 | 7.55 | ND | ND | ND | ND | C |
| 7418 | Zu-1 | Switzerland | 292.00 | 84.97 | 57.42 | 7.36 | ND | ND | ND | ND | C |
| 7424 | Jl-3 | Czech | 171.50 | 136.25 | 47.83 | 7.90 | 650.54 | 86.47 | -2.79 | 4.48 | C |
| 7438 | N13 | Russia | 144.00 | 104.89 | 57.81 | 8.06 | 485.42 | 81.23 | -0.69 | 4.20 | C |
| 7477 | WAR | USA | 235.50 | 79.63 | 57.54 | 7.49 | ND | ND | ND | ND | C |
| 7514 | RRS-7 | USA | 295.00 | 149.70 | 57.82 | 7.12 | 816.22 | 97.62 | 3.95 | 3.66 | C |
| 7519 | Ömö2-3 | Sweden | 194.50 | 128.78 | 57.80 | 7.26 | 625.71 | 102.13 | -5.20 | 3.81 | C |
| 7520 | Lp2-2 | Czech | 291.00 | 73.45 | 57.44 | 7.23 | ND | ND | ND | ND | C |
| 7521 | Lp2-6 | Czech | 266.00 | 340.15 | 57.69 | NA | 724.12 | 76.71 | 0.46 | 4.03 | C |
| 7522 | Mr-0 | Itary | 33.00 | 52.16 | 57.94 | 9.97 | 642.79 | 163.14 | -5.75 | 4.71 | C |
| 7523 | Pna-17 | USA | 346.00 | 182.87 | 57.84 | 7.42 | 742.23 | 95.92 | 5.45 | 4.02 | C |
| 7525 | Rmx-A180 | USA | 251.00 | 210.12 | 57.55 | 7.58 | ND | ND | ND | ND | C |
| 8214 | Gy-0 | France | 255.50 | 117.42 | 50.38 | 7.25 | 593.18 | 89.12 | -2.28 | 4.34 | C |
| 8215 | Fei-0 | Portugal | 261.00 | 88.84 | 57.13 | 7.77 | 564.26 | 114.18 | -6.46 | 4.06 | C |
| 8233 | Dem-4 | USA | 269.00 | 139.98 | 57.46 | 7.74 | 772.48 | 45.52 | 2.36 | 4.04 | C |
| 8236 | Hsm | Czech | 348.00 | 98.51 | 47.98 | 6.78 | 855.57 | 59.02 | 0.96 | 4.25 | C |
| 8240 | Kulturen-1 | Sweden | 274.00 | 128.55 | 57.71 | 7.68 | 627.40 | 53.31 | 3.18 | 4.01 | C |
| 8241 | Liarum | Sweden | 205.50 | 190.61 | 57.90 | 8.10 | 628.58 | 50.29 | -1.61 | 4.24 | C |
| 8245 | Seattle-0 | USA | 177.50 | 161.26 | 57.47 | 7.72 | ND | ND | ND | ND | H |
| 8249 | Vimmerby | Sweden | 229.00 | 95.79 | 57.75 | 7.59 | 639.78 | 84.40 | -0.60 | 4.11 | C |
| 8256 | Bå1-2 | Sweden | 386.50 | 298.62 | 50.62 | 7.68 | 677.80 | 94.26 | -1.56 | 4.22 | C |
| 8258 | Bå4-1 | Sweden | 299.00 | 146.86 | 51.55 | 7.27 | 705.96 | 54.41 | -1.12 | 4.21 | C |
| 8259 | Bå5-1 | Sweden | 308.00 | 125.62 | 51.27 | 7.76 | 719.20 | 86.78 | 1.18 | 4.15 | C |
| 8265 | Blh-1 | Czech | 252.50 | 152.99 | 57.54 | 7.55 | ND | ND | ND | ND | C |
| 8270 | Bs-1 | Switzerland | 283.00 | 227.77 | 56.32 | 7.87 | 724.54 | 144.87 | -4.06 | 4.45 | C |
| 8271 | Bu-0 | Germany | 370.00 | 92.72 | 56.01 | 7.26 | 1038.26 | 125.54 | -1.07 | 4.58 | C |
| 8284 | DraII-1 | Czech | 329.00 | 101.77 | 57.18 | 8.24 | 833.79 | 53.06 | 0.66 | 4.13 | C |
| 8285 | DraIII-1 | Czech | 357.00 | 313.69 | 48.06 | 7.84 | 821.53 | 158.18 | -6.86 | 4.10 | C |
| 8290 | En-1 | Germany | 294.00 | 142.48 | 45.33 | 7.28 | 685.56 | 101.88 | -4.81 | 4.03 | C |
| 8296 | Gd-1 | Germany | 262.00 | 116.39 | 50.69 | 7.44 | 753.09 | 92.17 | -2.70 | 4.10 | C |
| 8300 | Gr-1 | Austria | 279.00 | 91.50 | 49.17 | 6.97 | 764.09 | 103.12 | 4.89 | 4.11 | C |
| 8304 | Hi-0 | Netherlands | 271.00 | 107.89 | 48.06 | 6.86 | 776.44 | 80.94 | -2.88 | 4.35 | C |
| 8310 | Hs-0 | Germany | 393.00 | 160.15 | 48.02 | 7.47 | 917.83 | 79.21 | -0.48 | 3.97 | C |
| 8311 | In-0 | Austria | 324.00 | 383.22 | 57.29 | 8.33 | 827.97 | 89.10 | -4.67 | 4.29 | C |
| 8312 | Is-0 | Germany | 304.50 | 124.84 | 47.63 | 6.97 | 904.86 | 216.96 | -10.92 | 4.38 | C |
| 8313 | Jm-0 | Czech | 318.00 | 389.45 | 47.77 | 7.42 | 826.24 | 314.00 | -12.98 | 4.16 | C |
| 8314 | Ka-0 | Austria | 392.00 | 344.09 | 58.11 | 7.89 | 792.84 | 116.00 | -0.23 | 4.32 | C |
| 8323 | Lc-0 | UK | 221.50 | 147.55 | 57.69 | 7.67 | 408.41 | 52.06 | -1.10 | 3.90 | C |

| | | | | | | | | | | | |
|---|---|---|---|---|---|---|---|---|---|---|---|
| 8325 | Lip-0 | Poland | 350.00 | 201.17 | 57.63 | 7.69 | 620.03 | 71.78 | -0.36 | 3.88 | C |
| 8329 | Lm-2 | France | 356.50 | 257.40 | 57.68 | 7.24 | 759.93 | 105.32 | -4.73 | 4.06 | C |
| 8334 | Lu-1 | Sweden | 189.50 | 109.07 | 57.23 | 8.04 | ND | ND | ND | ND | C |
| 8337 | Mir-0 | Itary | 101.00 | 42.00 | 58.26 | 7.37 | ND | ND | ND | ND | C |
| 8343 | Na-1 | France | 203.00 | 128.17 | 57.99 | 6.87 | ND | ND | ND | ND | C |
| 8348 | Nw-0 | Germany | 258.00 | 92.03 | 58.03 | 6.60 | ND | ND | ND | ND | C |
| 8353 | Pa-1 | Itary | 302.00 | 173.34 | 57.76 | 7.22 | 526.96 | 129.49 | -6.07 | 4.03 | C |
| 8354 | Per-1 | Russia | 193.00 | 55.96 | 58.26 | 7.83 | ND | ND | ND | ND | C |
| 8366 | Rd-0 | Germany | 244.50 | 173.24 | 58.73 | 7.86 | ND | ND | ND | ND | C |
| 8374 | Rsch-4 | Russia | 391.50 | 193.14 | 57.50 | 7.47 | ND | ND | ND | ND | C |
| 8378 | Sap-0 | Czech | 315.00 | 120.73 | 57.64 | 7.01 | ND | ND | ND | ND | C |
| 8387 | St-0 | Sweden | 272.00 | 83.10 | 57.54 | 6.81 | ND | ND | ND | ND | C |
| 8388 | Stw-0 | Russia | 226.00 | 252.34 | 57.54 | 6.72 | ND | ND | ND | ND | C |
| 8395 | Tu-0 | Itary | 441.50 | 127.77 | 57.49 | 6.89 | ND | ND | ND | ND | C |
| 8420 | Kelsterbach-4 | Germany | 370.00 | 144.82 | 57.97 | 7.43 | 676.23 | 113.79 | -4.57 | 4.00 | C |
| 9104 | Lag1-6 | Georgia | 253.50 | 117.59 | 57.73 | 7.29 | 688.64 | 94.81 | 1.15 | 3.95 | C |
| 9165 | Truk-5 | Ukraine | 167.00 | 306.85 | 57.48 | 7.63 | ND | ND | ND | ND | C |
| 9302 | Edinburgh-5 | UK | 188.00 | 129.58 | 48.90 | 7.02 | 621.72 | 110.45 | -4.47 | 4.07 | C |
| 9308 | Ullapool-3 | UK | 237.00 | 128.81 | 57.76 | 7.55 | ND | ND | ND | ND | C |
| 100000 | Wil-1-Dean-Lab | Lithuania | 285.50 | 108.26 | 57.85 | 6.66 | ND | ND | ND | ND | C |
| Average | | | 267.00 | 136.25 | 57.50 | 7.67 | 709.73 | 102.51 | -2.77 | 4.15 | – |
| Standard deviation | | | 80.47 | 106.41 | 3.06 | 0.51 | 130.08 | 55.83 | 3.66 | 0.25 | – |

[a]ND indicates that no data is available.

[b]Average value (an average of five values measured for 2 DAG - 6 DAG, respectively) was used.

**Table S2. Gene models in proximity of the significantly associated locus for Horizontality under NPA treatment, Related to Figure 1, 2, 3 and 4.**

TAIR10 gene models in proximity (3 KB distance) of the significant SNP (passing the Bonferroni corrected threshold for approximately 215,000 tests, *p*-value < $10^{-7}$).

| Chromosome | SNP position | GWAS *p*-value | Effect size [%][a] | Associated traits | Associated gene | | | | |
|---|---|---|---|---|---|---|---|---|---|
| | | | | | Gene start | Gene end | Distance between SNP and gene body | AGI code | Gene annotation |
| 5 | 21256465 | $4.65 \times 10^{-8}$ | 13.58 | Horizontality | 21250802 | 21253939 | 2526 | AT5G52340 | exocyst subunit exo70 family protein A2 |
| | | | | | 21254911 | 21257618 | 0 (on gene) | AT5G52350 | exocyst subunit exo70 family protein A3 |
| | | | | | 21257834 | 21257915 | 1369 | AT5G52355 | pre-tRNA |
| | | | | | 21257934 | 21259342 | 1469 | AT5G52360 | actin depolymerizing factor 10 |

[a] Calculated as previously reported (Kerdaffrec *et al.* 2016, eLife)

#### Table. S3. Seed production under control (sufficient water) and drought conditions, Related to Figure 7

Haplogroup: classification based on haplotypes around the *EXOCYST70A3* locus. C: Common-sensitivity to NPA; H: High-sensitivity to NPA.; Treatment: water conditions during plant growth. Control: 65% water/soil.; Drought: 25% water/soil.; Replicate: experimental replicates; GWAS ID: Identifier of *A. thaliana* natural accessions as used in GWAPP (Seren et al. 2012).

| General name | Haplogroup | Treatment | Replicate | Seed number/plant | Flowering time (days after germination)[a,b] | GWAS ID |
|---|---|---|---|---|---|---|
| Col-0 | - (C) | Control | 1 | 59954 | 23 | 6909 |
| Col-0 | - (C) | Control | 1 | 44209 | 22 | 6909 |
| Col-0 | - (C) | Control | 1 | 48739 | 23 | 6909 |
| Col-0 | - (C) | Control | 1 | 41328 | 23 | 6909 |
| Col-0 | - (C) | Control | 2 | 41088 | 23 | 6909 |
| Col-0 | - (C) | Control | 2 | 59202 | 23 | 6909 |
| Col-0 | - (C) | Control | 2 | 49795 | 23 | 6909 |
| Col-0 | - (C) | Control | 2 | 44522 | 23 | 6909 |
| Col-0 | - (C) | Control | 3 | 79380 | 24 | 6909 |
| Col-0 | - (C) | Control | 3 | 67973 | 23 | 6909 |
| Col-0 | - (C) | Control | 3 | 95171 | 24 | 6909 |
| Col-0 | - (C) | Control | 3 | 73260 | 24 | 6909 |
| Col-0 | - (C) | Control | 4 | 90249 | 23 | 6909 |
| Col-0 | - (C) | Control | 4 | 72053 | 23 | 6909 |
| Col-0 | - (C) | Control | 4 | 65993 | 22 | 6909 |
| Col-0 | - (C) | Control | 4 | 95371 | 22 | 6909 |
| Col-0 | - (C) | Drought | 1 | 3112 | 20 | 6909 |
| Col-0 | - (C) | Drought | 1 | 3213 | 20 | 6909 |
| Col-0 | - (C) | Drought | 1 | 0 | 20 | 6909 |
| Col-0 | - (C) | Drought | 1 | 1504 | 20 | 6909 |

| | | | | | | |
|---|---|---|---|---|---|---|
| Col-0 | - (C) | Drought | 2 | 1505 | 20 | 6909 |
| Col-0 | - (C) | Drought | 2 | 1086 | 20 | 6909 |
| Col-0 | - (C) | Drought | 2 | 1152 | 20 | 6909 |
| Col-0 | - (C) | Drought | 2 | 1691 | 20 | 6909 |
| Col-0 | - (C) | Drought | 3 | 1142 | 25 | 6909 |
| Col-0 | - (C) | Drought | 3 | 1288 | 25 | 6909 |
| Col-0 | - (C) | Drought | 3 | 1588 | 25 | 6909 |
| Col-0 | - (C) | Drought | 3 | 2401 | 25 | 6909 |
| Col-0 | - (C) | Drought | 4 | 1914 | 25 | 6909 |
| Col-0 | - (C) | Drought | 4 | 1815 | 25 | 6909 |
| Col-0 | - (C) | Drought | 4 | 1858 | 25 | 6909 |
| Col-0 | - (C) | Drought | 4 | 836 | 25 | 6909 |
| *EXO70A3ox* | - | Control | 1 | 46763 | 22 | - |
| *EXO70A3ox* | - | Control | 1 | 14531 | 22 | - |
| *EXO70A3ox* | - | Control | 1 | 37094 | 22 | - |
| *EXO70A3ox* | - | Control | 1 | 37296 | 22 | - |
| *EXO70A3ox* | - | Control | 2 | 28951 | 22 | - |
| *EXO70A3ox* | - | Control | 2 | 38528 | 22 | - |
| *EXO70A3ox* | - | Control | 2 | 35370 | 22 | - |
| *EXO70A3ox* | - | Control | 2 | 42682 | 22 | - |
| *EXO70A3ox* | - | Control | 3 | 59494 | 22 | - |
| *EXO70A3ox* | - | Control | 3 | 65354 | 22 | - |
| *EXO70A3ox* | - | Control | 3 | 44308 | 22 | - |
| *EXO70A3ox* | - | Control | 3 | 78958 | 22 | - |
| *EXO70A3ox* | - | Control | 4 | 30367 | 22 | - |
| *EXO70A3ox* | - | Control | 4 | 35279 | 22 | - |
| *EXO70A3ox* | - | Control | 4 | 50978 | 22 | - |
| *EXO70A3ox* | - | Drought | 1 | 1277 | 20 | - |
| *EXO70A3ox* | - | Drought | 1 | 1501 | 20 | - |
| *EXO70A3ox* | - | Drought | 1 | 1048 | 20 | - |
| *EXO70A3ox* | - | Drought | 1 | 495 | 20 | - |

| | | | | | | |
|---|---|---|---|---|---|---|
| *EXO70A3ox* | - | Drought | 2 | 685 | 20 | - |
| *EXO70A3ox* | - | Drought | 2 | 1764 | 20 | - |
| *EXO70A3ox* | - | Drought | 2 | 338 | 25 | - |
| *EXO70A3ox* | - | Drought | 2 | 1820 | 20 | - |
| *EXO70A3ox* | - | Drought | 3 | 1512 | 25 | - |
| *EXO70A3ox* | - | Drought | 3 | 739 | 25 | - |
| *EXO70A3ox* | - | Drought | 3 | 1104 | 25 | - |
| *EXO70A3ox* | - | Drought | 3 | 409 | 25 | - |
| *EXO70A3ox* | - | Drought | 4 | 0 | NF | - |
| *EXO70A3ox* | - | Drought | 4 | 691 | 25 | - |
| *EXO70A3ox* | - | Drought | 4 | 1294 | 25 | - |
| *EXO70A3ox* | - | Drought | 4 | 272 | 25 | - |
| Ler-1 | C | Control | 1 | 15576 | 23 | 6932 |
| Ler-1 | C | Control | 1 | 12186 | 23 | 6932 |
| Ler-1 | C | Control | 1 | 10182 | 24 | 6932 |
| Ler-1 | C | Control | 1 | 14526 | 24 | 6932 |
| Ler-1 | C | Control | 2 | 13498 | 23 | 6932 |
| Ler-1 | C | Control | 2 | 9977 | 23 | 6932 |
| Ler-1 | C | Control | 2 | 8471 | 24 | 6932 |
| Ler-1 | C | Control | 2 | 15375 | 23 | 6932 |
| Ler-1 | C | Control | 3 | 27440 | 23 | 6932 |
| Ler-1 | C | Control | 3 | 11290 | 23 | 6932 |
| Ler-1 | C | Control | 3 | 24710 | 23 | 6932 |
| Ler-1 | C | Control | 3 | 25591 | 23 | 6932 |
| Ler-1 | C | Control | 4 | 16051 | 23 | 6932 |
| Ler-1 | C | Control | 4 | 14780 | 23 | 6932 |
| Ler-1 | C | Control | 4 | 17081 | 23 | 6932 |
| Ler-1 | C | Control | 4 | 22333 | 23 | 6932 |
| Ler-1 | C | Drought | 1 | 1428 | 20 | 6932 |
| Ler-1 | C | Drought | 1 | 995 | 20 | 6932 |
| Ler-1 | C | Drought | 1 | 1417 | 20 | 6932 |

| | | | | | | |
|---|---|---|---|---|---|---|
| Ler-1 | C | Drought | 1 | 1895 | 20 | 6932 |
| Ler-1 | C | Drought | 2 | 2288 | 20 | 6932 |
| Ler-1 | C | Drought | 2 | 1903 | 20 | 6932 |
| Ler-1 | C | Drought | 2 | 1832 | 20 | 6932 |
| Ler-1 | C | Drought | 2 | 836 | 20 | 6932 |
| Ler-1 | C | Drought | 3 | 1357 | 25 | 6932 |
| Ler-1 | C | Drought | 3 | 1180 | 25 | 6932 |
| Ler-1 | C | Drought | 3 | 806 | 25 | 6932 |
| Ler-1 | C | Drought | 3 | 884 | 25 | 6932 |
| Ler-1 | C | Drought | 4 | 1050 | 25 | 6932 |
| Ler-1 | C | Drought | 4 | 1215 | 25 | 6932 |
| Ler-1 | C | Drought | 4 | 1273 | 25 | 6932 |
| Ler-1 | C | Drought | 4 | 1158 | 25 | 6932 |
| Shakdara | C | Control | 1 | 28800 | 22 | 6962 |
| Shakdara | C | Control | 1 | 18290 | 24 | 6962 |
| Shakdara | C | Control | 1 | 21802 | 22 | 6962 |
| Shakdara | C | Control | 1 | 18274 | 24 | 6962 |
| Shakdara | C | Control | 2 | 31040 | 22 | 6962 |
| Shakdara | C | Control | 2 | 44968 | 22 | 6962 |
| Shakdara | C | Control | 2 | 19286 | 22 | 6962 |
| Shakdara | C | Control | 2 | 23959 | 22 | 6962 |
| Shakdara | C | Control | 3 | 71903 | 22 | 6962 |
| Shakdara | C | Control | 3 | 37200 | 22 | 6962 |
| Shakdara | C | Control | 3 | 34002 | 22 | 6962 |
| Shakdara | C | Control | 3 | 54017 | 22 | 6962 |
| Shakdara | C | Control | 4 | 35088 | 23 | 6962 |
| Shakdara | C | Control | 4 | 41748 | 19 | 6962 |
| Shakdara | C | Control | 4 | 39138 | 24 | 6962 |
| Shakdara | C | Control | 4 | 37598 | 23 | 6962 |
| Shakdara | C | Drought | 1 | 1648 | 20 | 6962 |
| Shakdara | C | Drought | 1 | 1902 | 20 | 6962 |

| | | | | | | |
|---|---|---|---|---|---|---|
| Shakdara | C | Drought | 1 | 1604 | 20 | 6962 |
| Shakdara | C | Drought | 1 | 1710 | 20 | 6962 |
| Shakdara | C | Drought | 2 | 2247 | 20 | 6962 |
| Shakdara | C | Drought | 2 | 2142 | 20 | 6962 |
| Shakdara | C | Drought | 2 | 1797 | 25 | 6962 |
| Shakdara | C | Drought | 2 | 1852 | 20 | 6962 |
| Shakdara | C | Drought | 3 | 3038 | 20 | 6962 |
| Shakdara | C | Drought | 3 | 2453 | 20 | 6962 |
| Shakdara | C | Drought | 3 | 2012 | 20 | 6962 |
| Shakdara | C | Drought | 3 | 1815 | 20 | 6962 |
| Shakdara | C | Drought | 4 | 2041 | 25 | 6962 |
| Shakdara | C | Drought | 4 | 1402 | 25 | 6962 |
| Shakdara | C | Drought | 4 | 1479 | 25 | 6962 |
| Shakdara | C | Drought | 4 | 945 | 20 | 6962 |
| Lp2-2 | C | Control | 1 | 59564 | 27 | 7520 |
| Lp2-2 | C | Control | 1 | 57462 | 27 | 7520 |
| Lp2-2 | C | Control | 1 | 52994 | 28 | 7520 |
| Lp2-2 | C | Control | 1 | 59428 | 27 | 7520 |
| Lp2-2 | C | Control | 2 | 58531 | 26 | 7520 |
| Lp2-2 | C | Control | 2 | 40511 | 27 | 7520 |
| Lp2-2 | C | Control | 2 | 49896 | 27 | 7520 |
| Lp2-2 | C | Control | 2 | 73265 | 27 | 7520 |
| Lp2-2 | C | Control | 3 | 102778 | 26 | 7520 |
| Lp2-2 | C | Control | 3 | 109754 | 26 | 7520 |
| Lp2-2 | C | Control | 3 | 82600 | 26 | 7520 |
| Lp2-2 | C | Control | 3 | 112543 | 26 | 7520 |
| Lp2-2 | C | Control | 4 | 72530 | 26 | 7520 |
| Lp2-2 | C | Control | 4 | 72409 | 26 | 7520 |
| Lp2-2 | C | Control | 4 | 69163 | 26 | 7520 |
| Lp2-2 | C | Control | 4 | 89082 | 26 | 7520 |
| Lp2-2 | C | Drought | 1 | 884 | 25 | 7520 |

| | | | | | | |
|---|---|---|---|---|---|---|
| Lp2-2 | C | Drought | 1 | 815 | 20 | 7520 |
| Lp2-2 | C | Drought | 1 | 1068 | 20 | 7520 |
| Lp2-2 | C | Drought | 1 | 988 | 20 | 7520 |
| Lp2-2 | C | Drought | 2 | 561 | 25 | 7520 |
| Lp2-2 | C | Drought | 2 | 856 | 25 | 7520 |
| Lp2-2 | C | Drought | 2 | 1241 | 25 | 7520 |
| Lp2-2 | C | Drought | 2 | 96 | 20 | 7520 |
| Lp2-2 | C | Drought | 3 | 167 | 35 | 7520 |
| Lp2-2 | C | Drought | 3 | 0 | 35 | 7520 |
| Lp2-2 | C | Drought | 3 | 0 | 35 | 7520 |
| Lp2-2 | C | Drought | 3 | 5418 | 40 | 7520 |
| Lp2-2 | C | Drought | 4 | 0 | 30 | 7520 |
| Lp2-2 | C | Drought | 4 | 0 | 35 | 7520 |
| Lp2-2 | C | Drought | 4 | 0 | 35 | 7520 |
| Lp2-2 | C | Drought | 4 | 0 | 30 | 7520 |
| Per-1 | C | Control | 1 | 41658 | 26 | 8354 |
| Per-1 | C | Control | 1 | 44868 | 24 | 8354 |
| Per-1 | C | Control | 1 | 27103 | 25 | 8354 |
| Per-1 | C | Control | 1 | 57143 | 23 | 8354 |
| Per-1 | C | Control | 2 | 61402 | 24 | 8354 |
| Per-1 | C | Control | 2 | 48125 | 24 | 8354 |
| Per-1 | C | Control | 2 | 56719 | 25 | 8354 |
| Per-1 | C | Control | 2 | 68521 | 26 | 8354 |
| Per-1 | C | Control | 3 | 61320 | 23 | 8354 |
| Per-1 | C | Control | 3 | 79596 | 25 | 8354 |
| Per-1 | C | Control | 3 | 87498 | 23 | 8354 |
| Per-1 | C | Control | 3 | 90758 | 24 | 8354 |
| Per-1 | C | Control | 4 | 70278 | 24 | 8354 |
| Per-1 | C | Control | 4 | 67210 | 24 | 8354 |
| Per-1 | C | Control | 4 | 61904 | 23 | 8354 |
| Per-1 | C | Control | 4 | 93404 | 23 | 8354 |

| | | | | | | |
|---|---|---|---|---|---|---|
| Per-1 | C | Drought | 1 | 1024 | 25 | 8354 |
| Per-1 | C | Drought | 1 | 1217 | 25 | 8354 |
| Per-1 | C | Drought | 1 | 946 | 20 | 8354 |
| Per-1 | C | Drought | 1 | 2897 | 20 | 8354 |
| Per-1 | C | Drought | 2 | 1224 | 20 | 8354 |
| Per-1 | C | Drought | 2 | 1535 | 25 | 8354 |
| Per-1 | C | Drought | 2 | 680 | 20 | 8354 |
| Per-1 | C | Drought | 2 | 1075 | 20 | 8354 |
| Per-1 | C | Drought | 3 | 98 | 35 | 8354 |
| Per-1 | C | Drought | 3 | 5683 | 40 | 8354 |
| Per-1 | C | Drought | 3 | 42 | 35 | 8354 |
| Per-1 | C | Drought | 3 | 264 | 35 | 8354 |
| Per-1 | C | Drought | 4 | 209 | 35 | 8354 |
| Per-1 | C | Drought | 4 | 114 | 35 | 8354 |
| Per-1 | C | Drought | 4 | 57 | 35 | 8354 |
| Per-1 | C | Drought | 4 | 6710 | 35 | 8354 |
| LAC-5 | H | Control | 1 | 60237 | 31 | 96 |
| LAC-5 | H | Control | 1 | 37245 | 31 | 96 |
| LAC-5 | H | Control | 1 | 60522 | 31 | 96 |
| LAC-5 | H | Control | 2 | 53159 | 31 | 96 |
| LAC-5 | H | Control | 2 | 61534 | 32 | 96 |
| LAC-5 | H | Control | 2 | 49903 | 32 | 96 |
| LAC-5 | H | Control | 2 | 44078 | 31 | 96 |
| LAC-5 | H | Control | 3 | 62516 | 32 | 96 |
| LAC-5 | H | Control | 3 | 72228 | 31 | 96 |
| LAC-5 | H | Control | 3 | 56917 | 31 | 96 |
| LAC-5 | H | Control | 3 | 49544 | 31 | 96 |
| LAC-5 | H | Control | 4 | 68069 | 31 | 96 |
| LAC-5 | H | Control | 4 | 53850 | 31 | 96 |
| LAC-5 | H | Control | 4 | 62850 | 31 | 96 |
| LAC-5 | H | Control | 4 | 82702 | 31 | 96 |

| | | | | | | |
|---|---|---|---|---|---|---|
| LAC-5 | H | Drought | 1 | 0 | 30 | 96 |
| LAC-5 | H | Drought | 1 | 0 | 30 | 96 |
| LAC-5 | H | Drought | 1 | 155 | 25 | 96 |
| LAC-5 | H | Drought | 1 | 96 | 30 | 96 |
| LAC-5 | H | Drought | 2 | 550 | 35 | 96 |
| LAC-5 | H | Drought | 2 | 0 | 30 | 96 |
| LAC-5 | H | Drought | 2 | 965 | 30 | 96 |
| LAC-5 | H | Drought | 2 | 370 | 35 | 96 |
| LAC-5 | H | Drought | 3 | 0 | 35 | 96 |
| LAC-5 | H | Drought | 3 | 0 | 35 | 96 |
| LAC-5 | H | Drought | 3 | 0 | 35 | 96 |
| LAC-5 | H | Drought | 3 | 0 | 35 | 96 |
| LAC-5 | H | Drought | 4 | 0 | NF | 96 |
| LAC-5 | H | Drought | 4 | 0 | NF | 96 |
| LAC-5 | H | Drought | 4 | 0 | NF | 96 |
| LAC-5 | H | Drought | 4 | 0 | NF | 96 |
| MIB-60 | H | Control | 1 | 32158 | 28 | 204 |
| MIB-60 | H | Control | 1 | 54078 | 28 | 204 |
| MIB-60 | H | Control | 1 | 31898 | 27 | 204 |
| MIB-60 | H | Control | 1 | 79740 | 27 | 204 |
| MIB-60 | H | Control | 2 | 45114 | 27 | 204 |
| MIB-60 | H | Control | 2 | 23998 | 27 | 204 |
| MIB-60 | H | Control | 2 | 23712 | 28 | 204 |
| MIB-60 | H | Control | 2 | 37364 | 28 | 204 |
| MIB-60 | H | Control | 3 | 54144 | 25 | 204 |
| MIB-60 | H | Control | 3 | 52656 | 27 | 204 |
| MIB-60 | H | Control | 3 | 76350 | 25 | 204 |
| MIB-60 | H | Control | 3 | 49329 | 27 | 204 |
| MIB-60 | H | Control | 4 | 59582 | 27 | 204 |
| MIB-60 | H | Control | 4 | 68096 | 28 | 204 |
| MIB-60 | H | Control | 4 | 51968 | 28 | 204 |

| | | | | | | |
|---|---|---|---|---|---|---|
| MIB-60 | H | Control | 4 | 64630 | 27 | 204 |
| MIB-60 | H | Drought | 1 | 993 | 20 | 204 |
| MIB-60 | H | Drought | 1 | 1443 | 20 | 204 |
| MIB-60 | H | Drought | 1 | 186 | 20 | 204 |
| MIB-60 | H | Drought | 1 | 349 | 20 | 204 |
| MIB-60 | H | Drought | 2 | 1637 | 20 | 204 |
| MIB-60 | H | Drought | 2 | 2195 | 20 | 204 |
| MIB-60 | H | Drought | 2 | 1246 | 20 | 204 |
| MIB-60 | H | Drought | 2 | 2041 | 20 | 204 |
| MIB-60 | H | Drought | 3 | 1490 | 30 | 204 |
| MIB-60 | H | Drought | 3 | 680 | 35 | 204 |
| MIB-60 | H | Drought | 3 | 283 | 35 | 204 |
| MIB-60 | H | Drought | 3 | 746 | 35 | 204 |
| MIB-60 | H | Drought | 4 | 1178 | 35 | 204 |
| MIB-60 | H | Drought | 4 | 612 | 35 | 204 |
| MIB-60 | H | Drought | 4 | 706 | 35 | 204 |
| MIB-60 | H | Drought | 4 | 408 | 35 | 204 |
| Coc-1 | H | Control | 1 | 49172 | 27 | 5729 |
| Coc-1 | H | Control | 1 | 26831 | 29 | 5729 |
| Coc-1 | H | Control | 1 | 42148 | 27 | 5729 |
| Coc-1 | H | Control | 1 | 29068 | 27 | 5729 |
| Coc-1 | H | Control | 2 | 40326 | 25 | 5729 |
| Coc-1 | H | Control | 2 | 53946 | 27 | 5729 |
| Coc-1 | H | Control | 2 | 38021 | 26 | 5729 |
| Coc-1 | H | Control | 2 | 52345 | 28 | 5729 |
| Coc-1 | H | Control | 3 | 52573 | 28 | 5729 |
| Coc-1 | H | Control | 3 | 52920 | 28 | 5729 |
| Coc-1 | H | Control | 3 | 53323 | 29 | 5729 |
| Coc-1 | H | Control | 3 | 72626 | 28 | 5729 |
| Coc-1 | H | Control | 4 | 78730 | 26 | 5729 |
| Coc-1 | H | Control | 4 | 64813 | 29 | 5729 |

| | | | | | | |
|---|---|---|---|---|---|---|
| Coc-1 | H | Control | 4 | 59413 | 27 | 5729 |
| Coc-1 | H | Control | 4 | 106848 | 27 | 5729 |
| Coc-1 | H | Drought | 1 | 887 | 30 | 5729 |
| Coc-1 | H | Drought | 1 | 538 | 25 | 5729 |
| Coc-1 | H | Drought | 1 | 637 | 25 | 5729 |
| Coc-1 | H | Drought | 1 | 371 | 30 | 5729 |
| Coc-1 | H | Drought | 2 | 488 | 25 | 5729 |
| Coc-1 | H | Drought | 2 | 0 | NF | 5729 |
| Coc-1 | H | Drought | 2 | 606 | 25 | 5729 |
| Coc-1 | H | Drought | 2 | 412 | 30 | 5729 |
| Coc-1 | H | Drought | 3 | 1176 | 25 | 5729 |
| Coc-1 | H | Drought | 3 | 605 | 25 | 5729 |
| Coc-1 | H | Drought | 3 | 636 | 20 | 5729 |
| Coc-1 | H | Drought | 3 | 1700 | 20 | 5729 |
| Coc-1 | H | Drought | 4 | 0 | NF | 5729 |
| Coc-1 | H | Drought | 4 | 0 | 45 | 5729 |
| Coc-1 | H | Drought | 4 | 0 | NF | 5729 |
| Coc-1 | H | Drought | 4 | 0 | 45 | 5729 |
| Ty-0 | H | Control | 1 | 18914 | 36 | 7351 |
| Ty-0 | H | Control | 1 | 17897 | 36 | 7351 |
| Ty-0 | H | Control | 1 | 21862 | 36 | 7351 |
| Ty-0 | H | Control | 1 | 15853 | 36 | 7351 |
| Ty-0 | H | Control | 2 | 14245 | 36 | 7351 |
| Ty-0 | H | Control | 2 | 18391 | 36 | 7351 |
| Ty-0 | H | Control | 2 | 23727 | 36 | 7351 |
| Ty-0 | H | Control | 2 | 21835 | 36 | 7351 |
| Ty-0 | H | Control | 3 | 30996 | 37 | 7351 |
| Ty-0 | H | Control | 3 | 17035 | 37 | 7351 |
| Ty-0 | H | Control | 3 | 16989 | 37 | 7351 |
| Ty-0 | H | Control | 3 | 25668 | 36 | 7351 |
| Ty-0 | H | Control | 4 | 43280 | 37 | 7351 |

| | | | | | | |
|---|---|---|---|---|---|---|
| Ty-0 | H | Control | 4 | 34630 | 37 | 7351 |
| Ty-0 | H | Control | 4 | 25637 | 37 | 7351 |
| Ty-0 | H | Control | 4 | 47174 | 37 | 7351 |
| Ty-0 | H | Drought | 1 | 0 | NF | 7351 |
| Ty-0 | H | Drought | 1 | 0 | NF | 7351 |
| Ty-0 | H | Drought | 1 | 0 | NF | 7351 |
| Ty-0 | H | Drought | 1 | 0 | NF | 7351 |
| Ty-0 | H | Drought | 2 | 0 | NF | 7351 |
| Ty-0 | H | Drought | 2 | 0 | NF | 7351 |
| Ty-0 | H | Drought | 2 | 0 | NF | 7351 |
| Ty-0 | H | Drought | 2 | 0 | NF | 7351 |
| Ty-0 | H | Drought | 3 | 78 | 30 | 7351 |
| Ty-0 | H | Drought | 3 | 130 | 30 | 7351 |
| Ty-0 | H | Drought | 3 | 0 | 30 | 7351 |
| Ty-0 | H | Drought | 3 | 26 | 30 | 7351 |
| Ty-0 | H | Drought | 4 | 0 | 30 | 7351 |
| Ty-0 | H | Drought | 4 | 0 | 30 | 7351 |
| Ty-0 | H | Drought | 4 | 0 | 30 | 7351 |
| Ty-0 | H | Drought | 4 | 0 | 30 | 7351 |

[a] NF: no flowering.

[b] Flowering was observed every five days in the drought experiment.

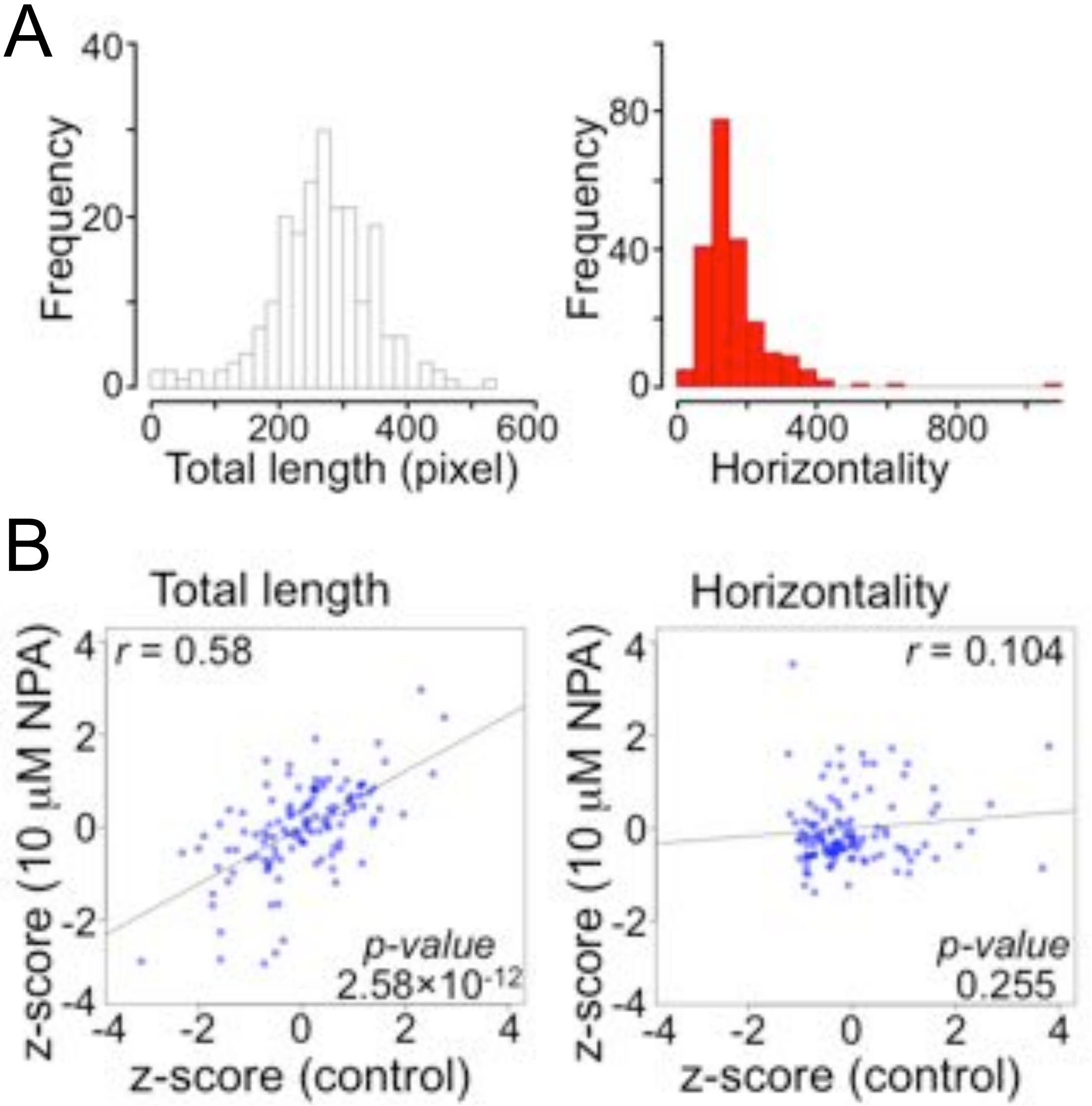


Figure S1 Ogura et al. 2018

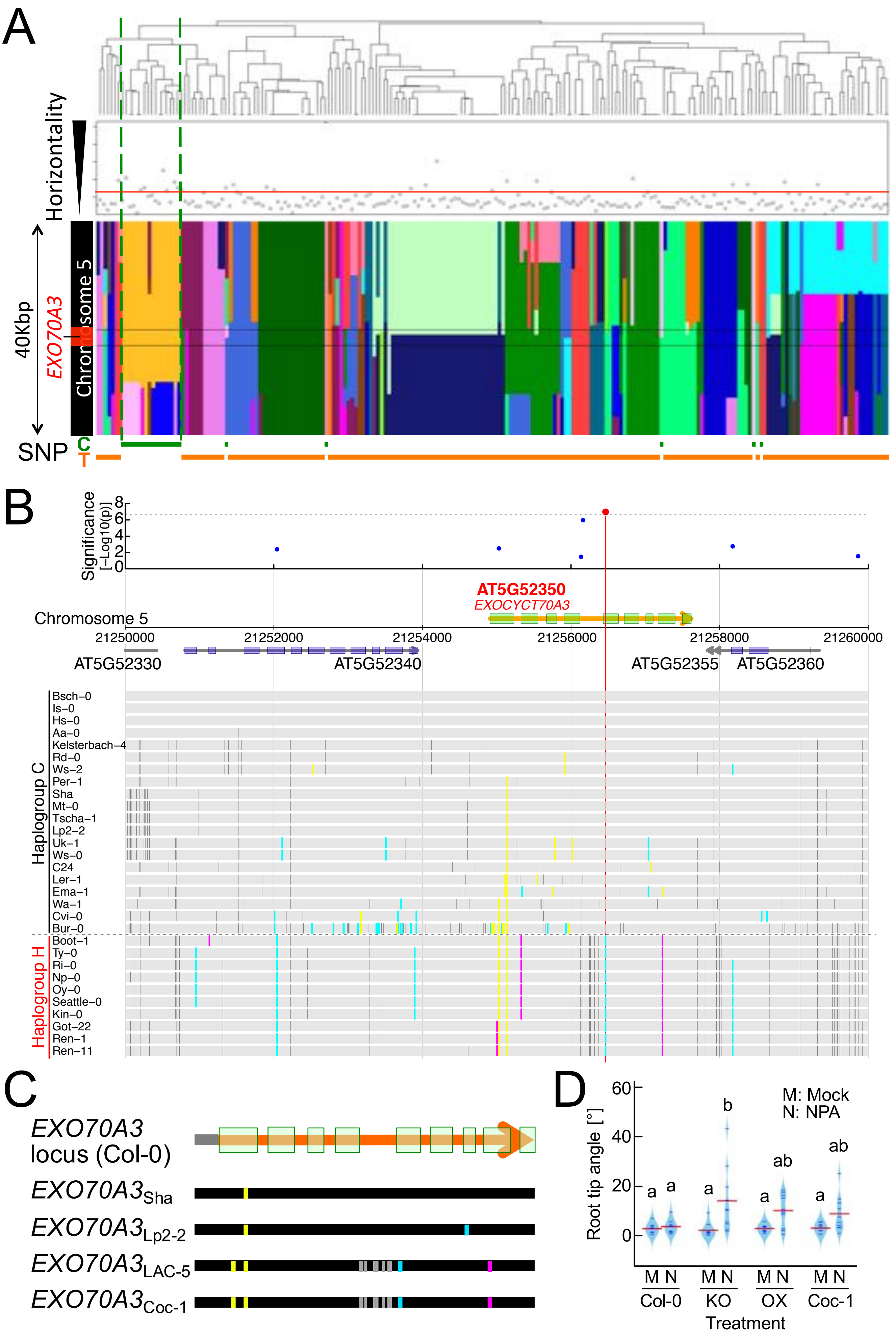


Figure S2 Ogura et al. 2018

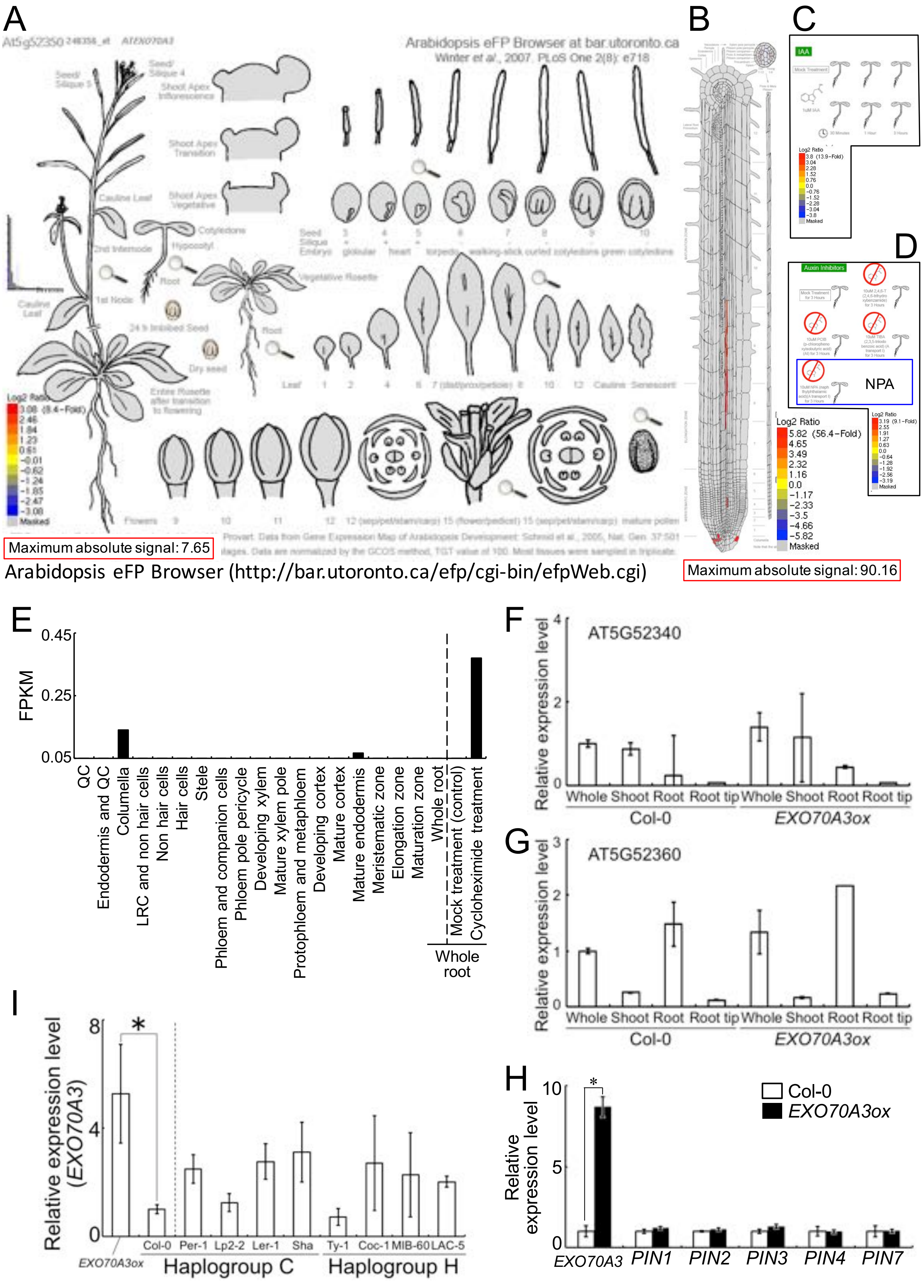


Figure S3 Ogura et al. 2018

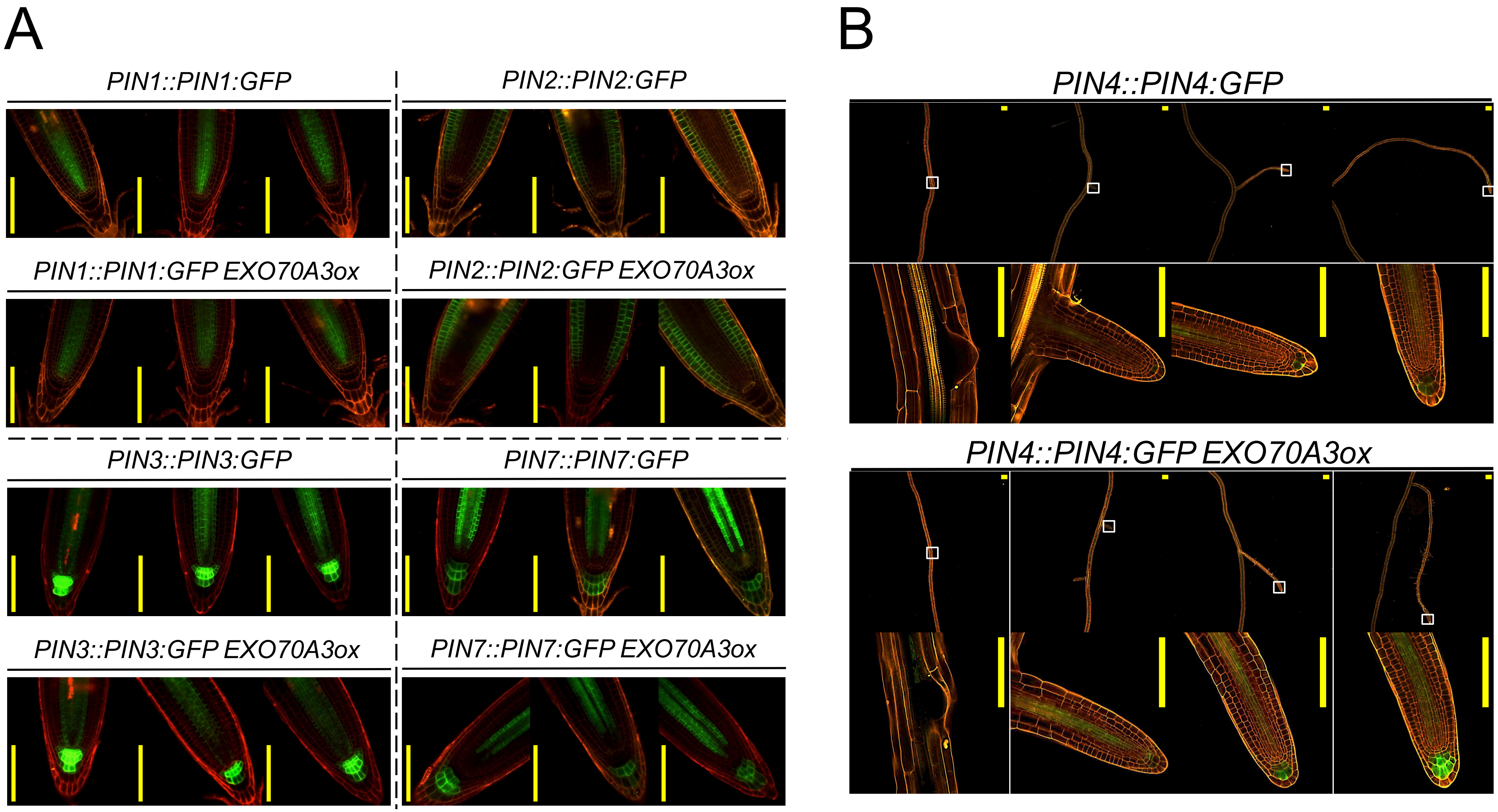


Figure S4 Ogura et al. 2018

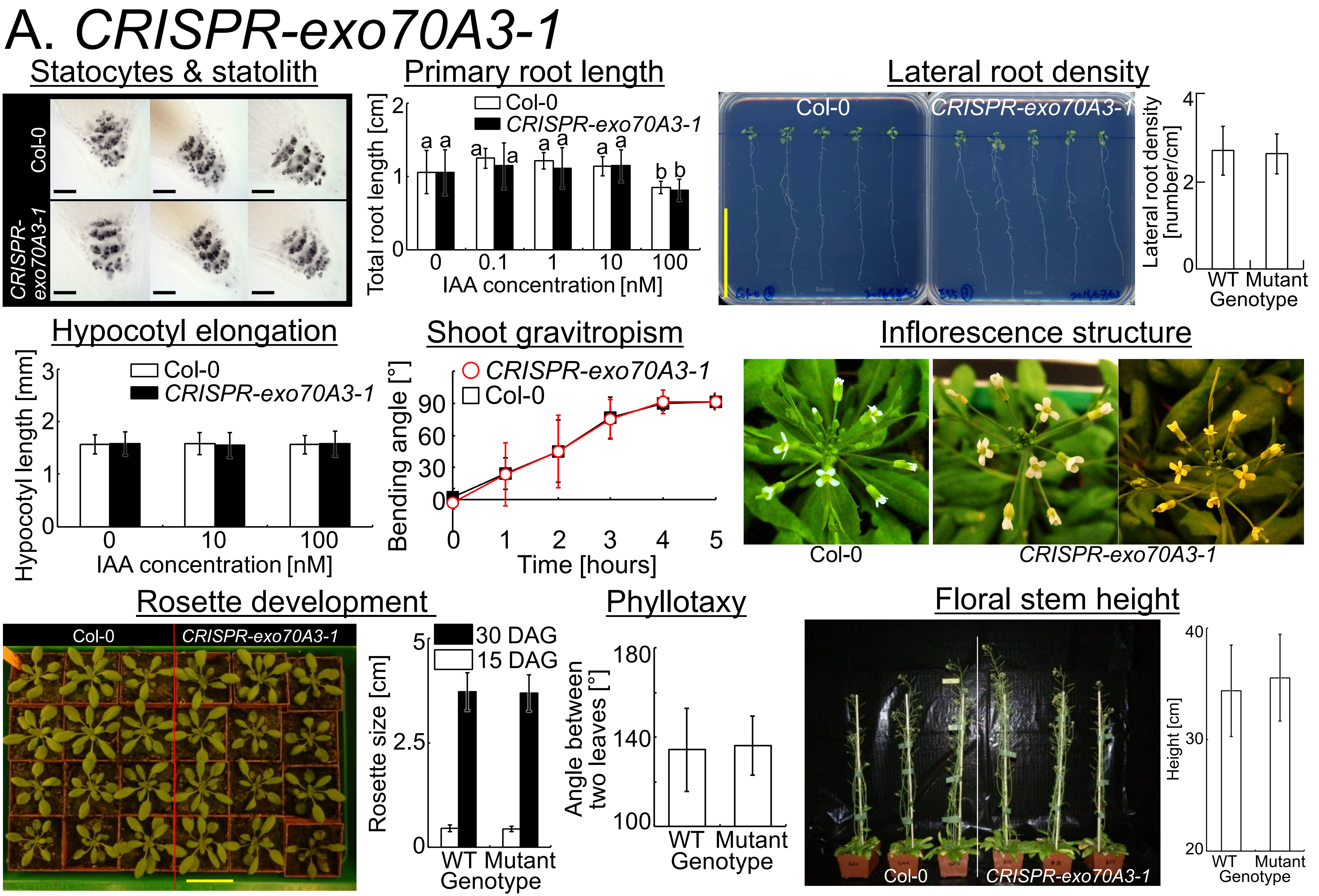


Figure S5 Ogura et al. 2018

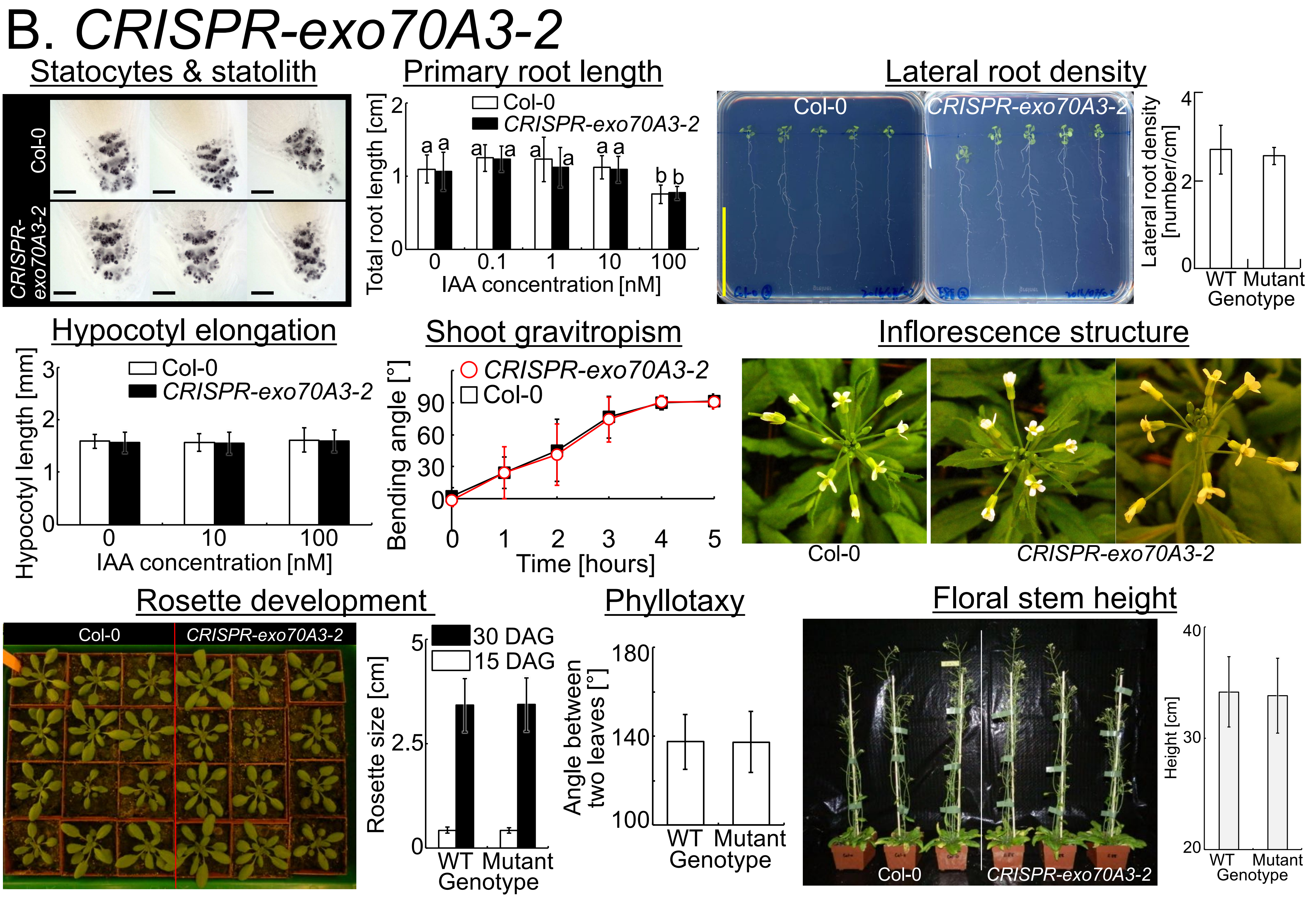


Figure S5 Ogura et al. 2018 (continued)

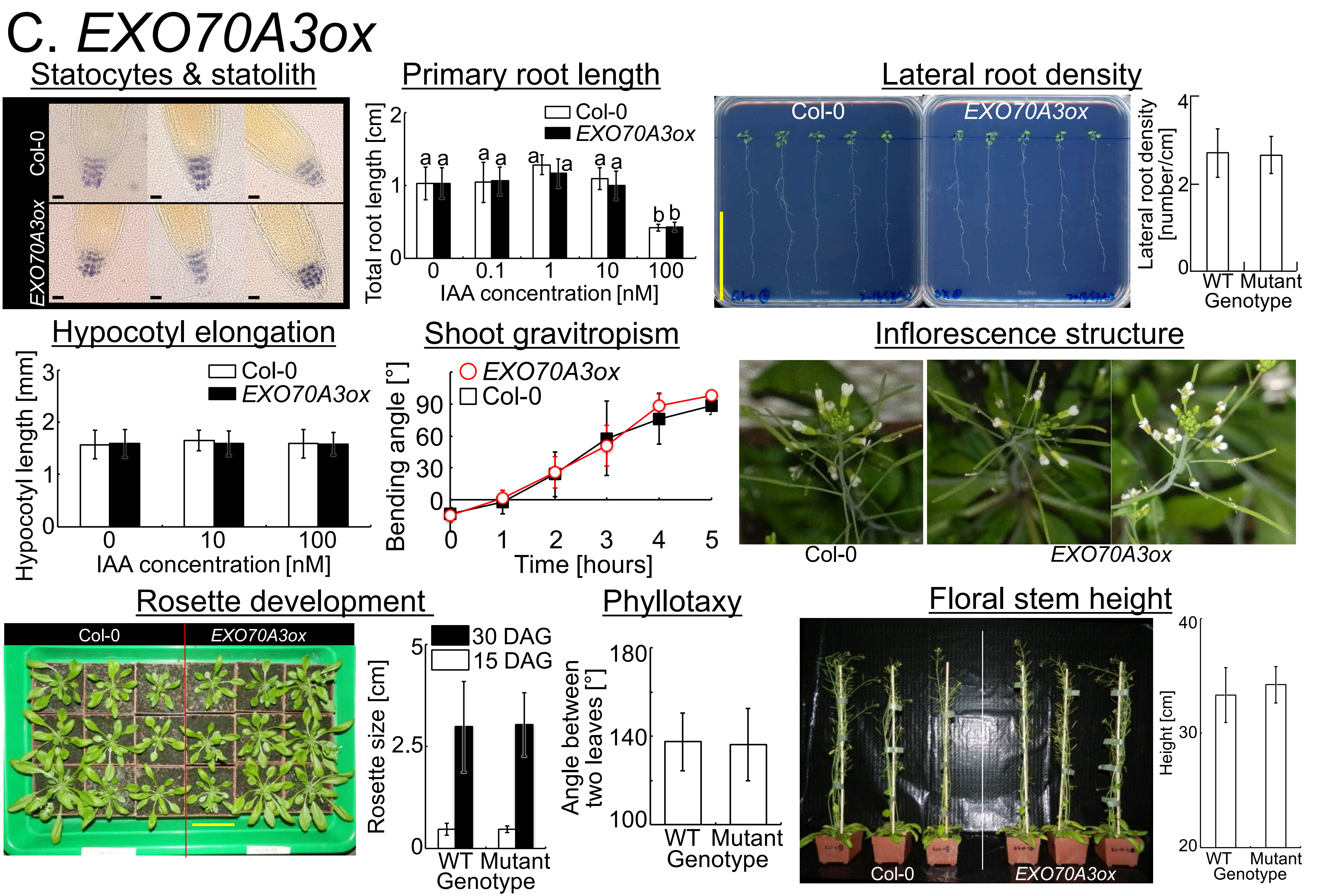


Figure S5 Ogura et al. 2018 (continued)

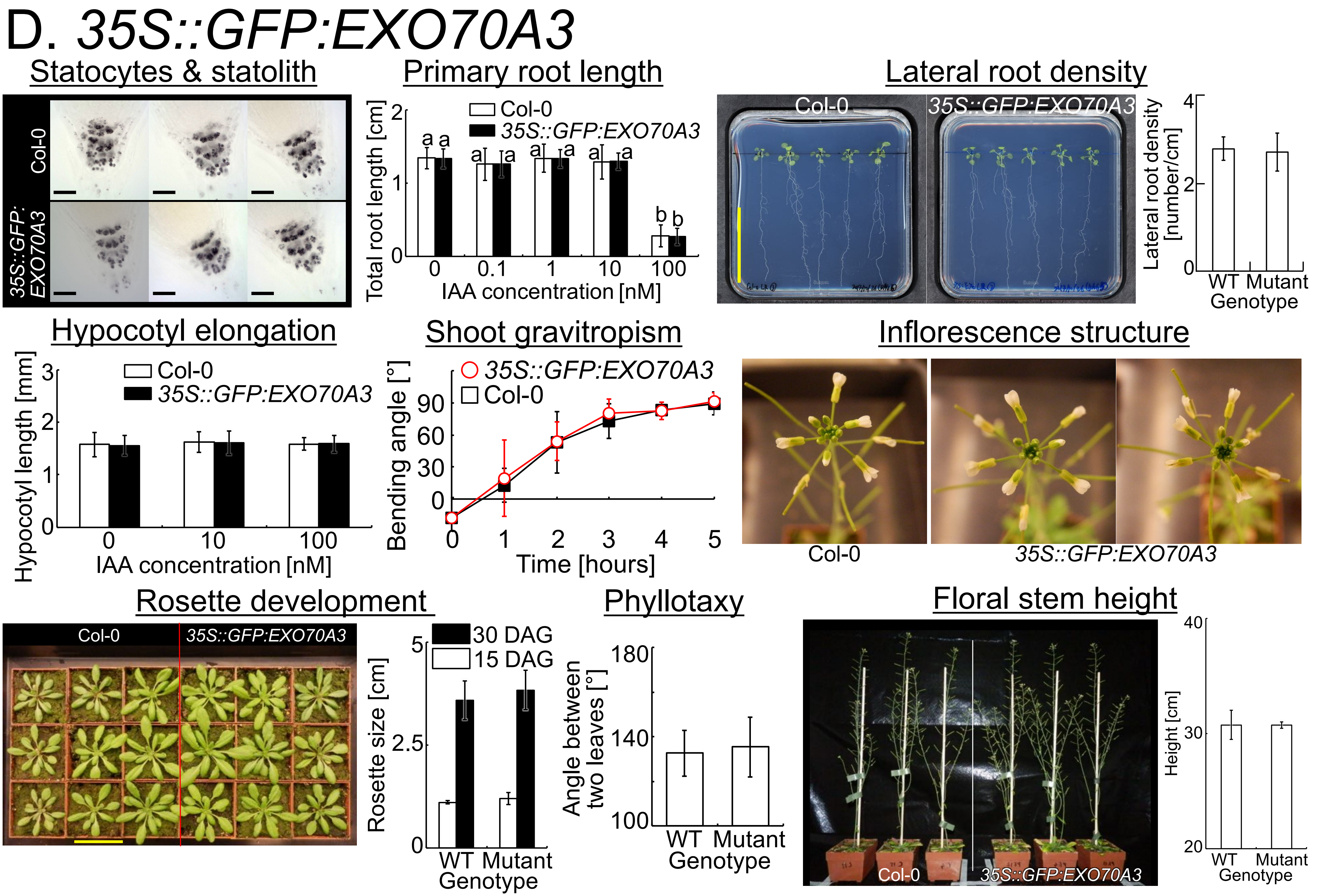


Figure S5 Ogura et al. 2018 (continued)

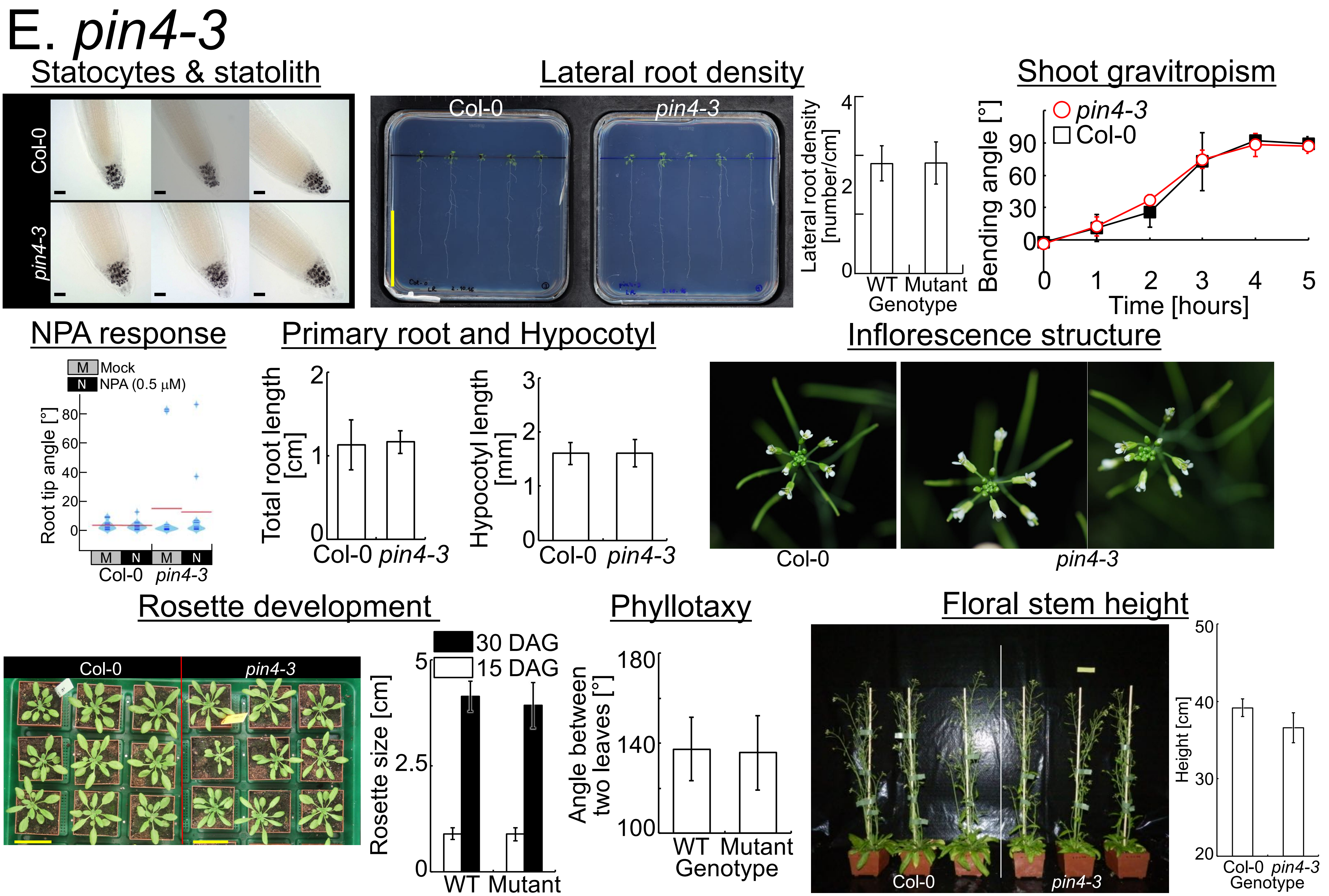


Figure S5 Ogura et al. 2018 (continued)

### F. Variances in root tip angles

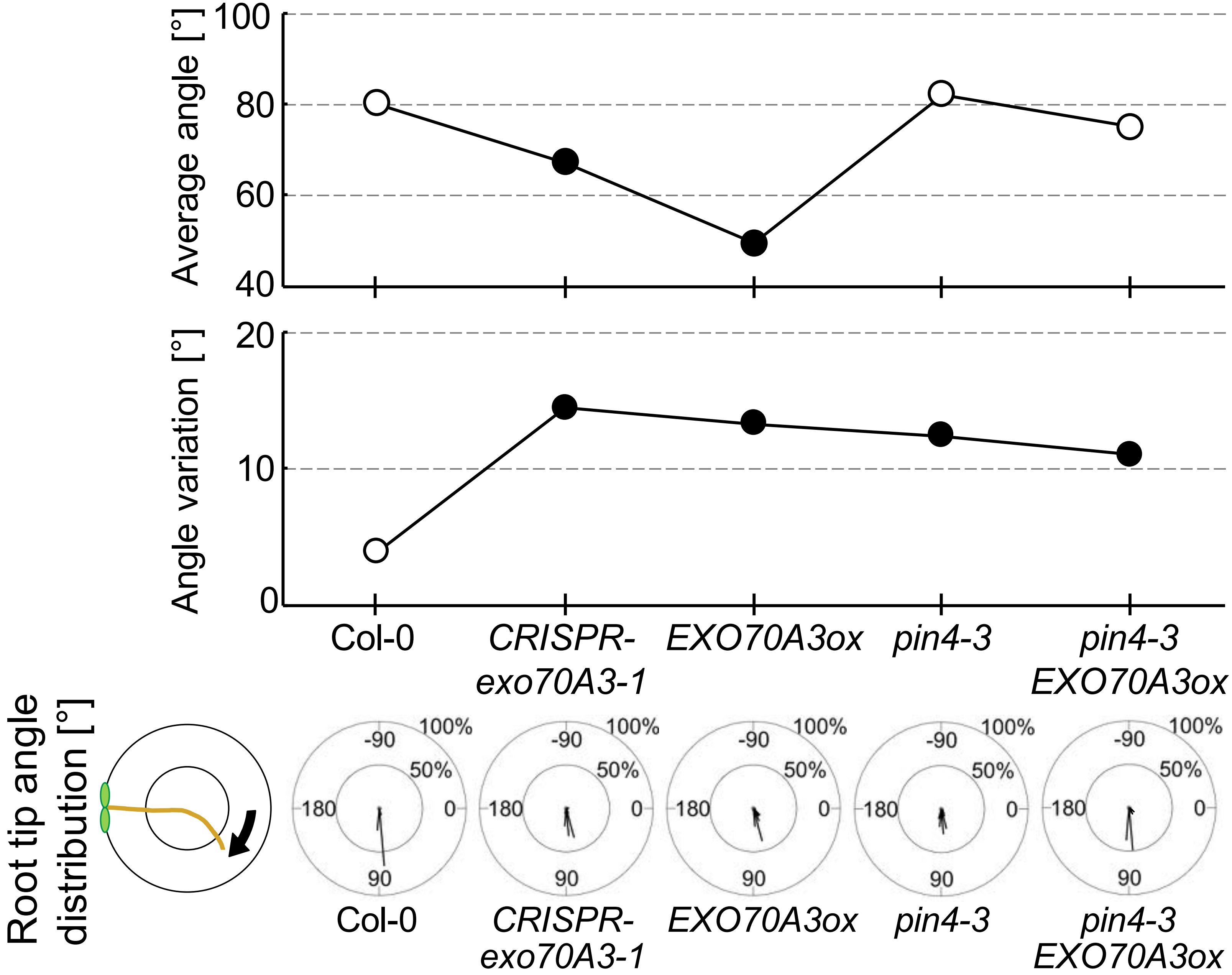


Figure S5 Ogura et al. 2018 (continued)

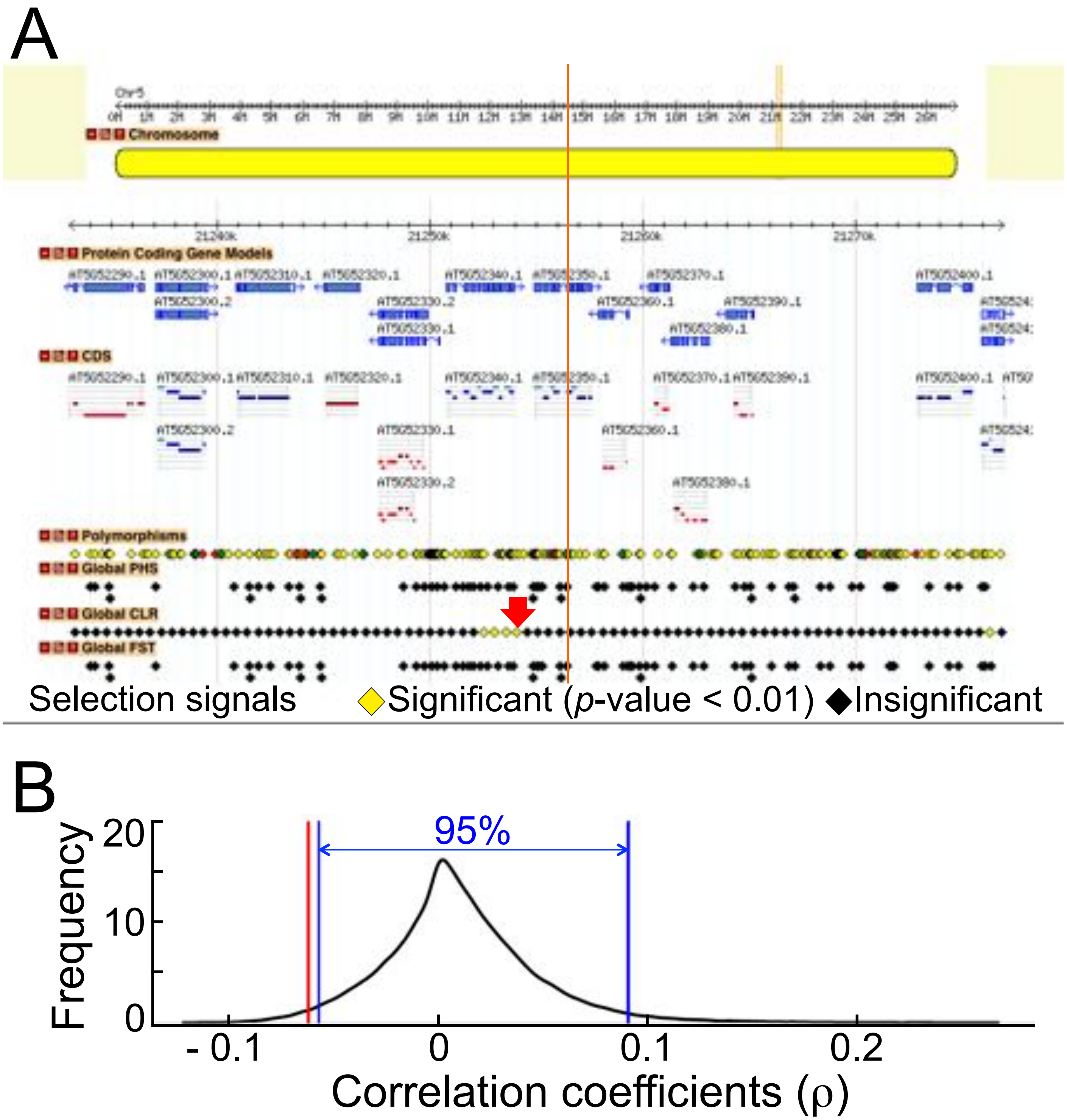


Figure S6 Ogura et al. 2018

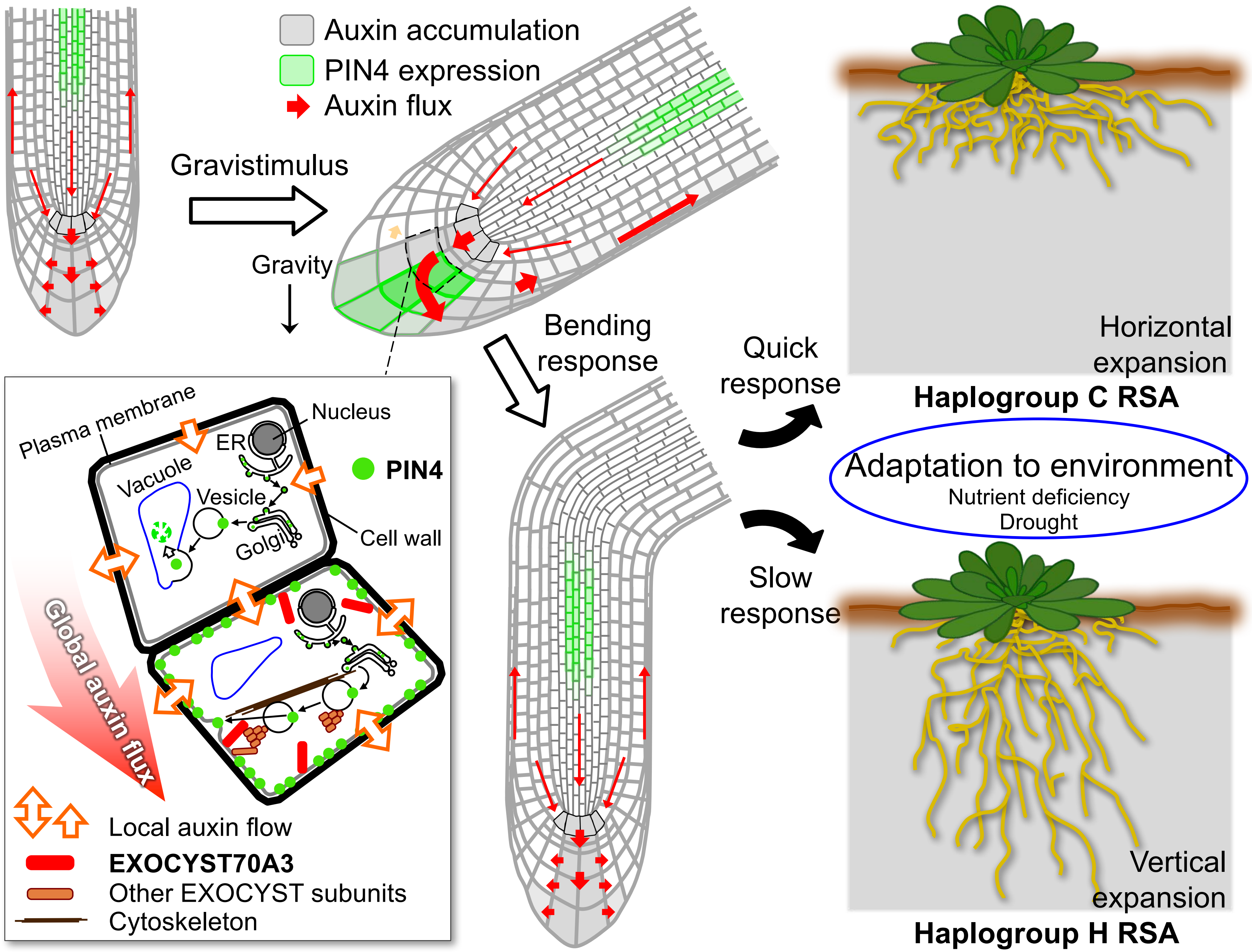


Figure S7 Ogura et al. 2018

**Supplemental Information**

**Supplemental Experimental Procedures**

**Plant materials and growth conditions**

All plants for the image acquisition on plates were grown on vertical 1 × MS agar plates (1 % sucrose, 0.8 (phenotyping for GWA mapping) or 1.0 % (other experiments) agar, pH = 5.7, phenotyping for GWA mapping: 35 ml/plate, other experiments: 53 ml/plate) in 12 cm × 12 cm square plates (Greiner Bio One International AG, Kremsmünster, Austria) that were sealed with Leucopore tape (1.25 cm width, Duchefa Biochemie B.V., Haarlem, The Netherlands), with or without chemicals under long day conditions (16 hours light) at 21°C. OSRAM Fluora L36W/77 (OSRAM Licht AG, Munich, Germany) was used as the light source. Indole-3-acetic acid (IAA), *N*-1-naphthylphthalamic acid (NPA), and propidium iodide were purchased from SIGMA-ALDRICH Co. LLC. (Saint Louis, MO, USA). Sucrose was obtained from AppliChem GmbH (Darmstad, Germany). Plant Agar and Murashige & Skoog medium incl. MES buffer were purchased from Duchefa Biochemie B.V. 6.5 cm × 6.5 cm × 5.2 cm plastic pots for plant growth on soil (Blumentopf Typ2) were obtained from Alpaco Lück GmbH & Co. KG (Meinerzhagen, Germany).

The seeds of the 215 natural accessions of *Arabidopsis thaliana* (Extended Data Table 1) were a gift from Dr. Magnus Nordborg's laboratory (GMI, Vienna, Austria). *CRISPR-exo70A3-1* and *CRISPR-exo70A3-2* were prepared following previously published method (Mao *et al.*, 2016) and genotypes were confirmed by sequencing. *EXO70A3ox* (SALK_075426) was purchased from Nottingham Arabidopsis Stock Center (Leicestershire, UK). *35S::GFP:EXO70A3* was a gift from Dr. Viktor Žárský (IEB AS CR, Prague, Czech Republic). *PIN1::PIN1:GFP*, *PIN2::PIN2:GFP*, *PIN3::PIN3:GFP*, *PIN4::PIN4:GFP*, *PIN7::PIN7:GFP*, *DR5::GFP* and *pin4-3* were a gift from Dr. Jürgen Kleine-Vehn (University of Natural Resources and Life Sciences, Vienna, Austria). Crosses between *EXO70A3*-related mutants and other mutants were made by manually pollinating emasculated pistils of those marker lines with *EXO70A3ox* pollen. For allelic complementation analysis, *EXO70A3* promoter sequence from Col-0 and EXO70A3 protein coding genomic sequence from five natural accessions (Col-0, Shakdara (Sha), Lp2-2, LAC-5 and Coc-1) were PCR amplified and cloned into pGreen0029 binary vector (Hellens *et al.* 2000) using In-Fusion Cloning system (Takara Bio Europe, Saint-Germain-en-Laye, France) to obtain 5 constructs that have fused *EXO70A3* promoter sequence from Col-0 with different *EXO70A3* protein coding genomic sequence. The five constructs and pGreen0029 itself as a vector control were used for floral dipping transformation to create transgenic lines. For each construct, at least three individual transformed lines were established, and a line which showed median phenotype in these lines in terms of NPA response was selected for further analysis as a representative line of the allele. Plants for crosses, seed amplification and phenotyping of aerial parts were grown on soil (PROFI SUBSTRAT, Einheitserde Werkverband e.V., Sinntal-Altengronau, Germany) with a bottom layer of Premium Perlite (Gramoflor GmbH & Co. KG, Vechta, Germany) at 21°C, 16 hours light/16°C, 8 hours dark conditions.

**Quantitative phenotyping of NPA treated roots for GWAS**

Seeds were sterilized by chlorine gas generated from 130 ml of 10% sodium hypochlorite and 3.9 ml of 37% hydrochloric acid and then stratified in sterile water for 3 days at 4°C in the dark. The 215 natural accessions were grown on 1 × MS agar plates containing 10 μM NPA under long day conditions (16 hours light) at 21°C. 24 plants were used for each accession. Starting at 2 days after germination (DAG), root images were acquired by scanners (EPSON

Perfection V600 Photo, Seiko Epson CO., Nagano, Japan) each 24 hours for 5 days (2 DAG – 6 DAG). Root image analyses and phenotype quantification were performed using the BRAT software (Slovak et al., 2014). In particular, from the traits quantified by BRAT, we used a length phenotype (Total length), a width trait (Average root width), and two derivatives of angle phenotypes of roots (Horizontality: Root horizontal index/ Root vertical index × Total length of the root, Skewness: the averages of angles of all lines, which were drawn between each pixel of the root and the root-hypocotyl junction, to the gravity vector). Horizontality represents both "the trend of root growth to horizontal direction" and "the size of root growth in horizontal direction". Simply stating, "Horizontality" shows the intensity of root growth in horizontal direction.

**Genome wide association studies (GWAS) and bioinformatics analyses**

GWAS were conducted using the average trait value for each trait of 215 natural accessions by an accelerated mixed model (EMMAX (Kang *et al.*, 2010)) followed by EMMA (Kang *et al.*, 2008) for the most significant 200 associations as implemented in the web-based GWAS application GWAPP (Seren *et al.*, 2012), conducted on December 15, 2014). For "horizontality" phenotype, box-cox transformation was conducted before GWAS to obtain better data distribution. All other statistical analyses in this study were performed using R language (http://www.R-project.org/). For the test of the correlation between phenotypes on control and NPA plates, z-scores for phenotype values of each accession were calculated according to the following formula: z = ("Individual phenotype value" – "Median of phenotype values")/"Standard deviation of phenotype values", and used for the analysis. For multiple tests, one-way, two-way or three-way factorial ANOVA (analysis of variance) was first conducted to assess the existence of distinct groups using a threshold $p$-value < 0.05. Then the post-hoc Tukey test was performed to find significantly different groups (threshold: $p$-value < 0.05). To test whether population structure does explain a significant portion of phenotypic variation observed between two haplogroups, the significance of phenotypic differences between haplogroups was assessed using the mixed model procedure as implemented in EMMAX (Kang *et al.*, 2010) with a kinship matrix estimated from the Regional Mapping Project data (Horton *et al.*, 2012). The analysis regarding signals of selection was conducted using the Selection Browser of the Regional Mapping Project (Horton *et al.*, 2012). The eFP browser (Winter *et al.*, 2007) was used as a general reference for gene expression profiles of candidate genes as well as a RNA-Seq data from Li *et al.* 2017 (Li *et al.*, 2017). The SALK genome browser created by the *Arabidopsis thaliana* 1001 genome project (Schmitz et al., 2013), signal.salk.edu/atg1001/3.0/gebrowser.php) was used to analyze polymorphism among natural accessions. Signals of selection was investigated using Selection Scan (Horton *et al.*, 2012). The search for an association between the SNP which we identified by GWAS using Horizontality and the climate condition was conducted by comparing correlations that were calculated between each SNP and 19 climate conditions (WorldClim, http://www.worldclim.org/) in Hancock *et al.*, 2011.

**Haplogroup analysis**

For grouping the 215 natural accessions of *A. thaliana* according to haplotypes around the single nucleotide polymorphisms (SNPs) of interest, SNPs in a 40KB window around the SNPs of interest were extracted from a pre-imputation version of the Regional Mapping Project panel described previously (Horton *et al.*, 2012). These SNPs were used as the input into fastPHASE (Scheet and Stephens, 2006) version 1.4.0, which was run using the default settings, with the exception of invoking the –Pzp option to output raw likelihood values. For each SNP in the analysis, the cluster membership with the highest likelihood was designated as the haplogroup for that SNP. In the resulting dendrograms, accessions were ordered using the distance matrix calculated from all SNPs in the window of interest.

**Plant observations**

All manual measurements and analyses of plant images were conducted using Fiji (Schindelin *et al.*, 2012). For root phenotyping of *CRISPR-exo70A3-1*, *CRISPR-exo70A3-2*, *EXO70A3ox*, *35S:GFP:EXO70A3* and *pin4-3* young seedlings, seedlings were grown on 1 × MS agar plates. For root angle changes of these mutants in response to NPA, we conducted transfer experiments instead of germinating the plants on NPA as for our GWAS screen, to directly and more precisely measure the effect of NPA on the direction of root growth. First seeds were sawn on normal 1 × MS agar plates and grown until 5 days after germination. Immediately after that, plants grown to a similar size were transferred to control (DMSO) or treatment plates that contained 0.1 μM NPA (allelic complementation) or 0.5 μM NPA (other experiments) and images were acquired one day after the transfer by scanners (EPSON Perfection V600 Photo). The angle of the root tip to the vector of gravity was measured from these images. For each genotype, 10 or more plants were used and the whole experiment was repeated at least twice. For lateral root density, plants grown on 1 × MS agar plates were transferred to fresh plates 5 days after germination (DAG) and images of plants were acquired 20 days after germination. Then numbers of lateral roots and the primary root lengths (cm) of plants were manually measured. The lateral root density (the number of lateral roots/cm) was calculated by dividing the lateral root numbers by the primary root lengths. To compare lateral root densities among natural accessions, we used eight accessions (haplogroup C: Ler-1, Shakdara (Sha), Lp2-2 and Per-1; haplogroup H: LAC-5, MIB-60, Coc-1 and Ty-0). For hypocotyl length and total root length measurements, various concentrations of IAA in DMSO solutions were added to 1 × MS agar plates (final IAA concentrations: 0.1, 1, 10 and 100 nM, DMSO concentration: 0.1% v/v). 0.1% DMSO plates were used as a mock treatment. Plants were grown on these plates for 5 days after germination and images were acquired. 10 or more plants of each genotype were used for each experiment and the experiment was repeated twice.

For the observation of statocytes, plants were grown on 1 × MS agar plates for 5 days after germination. They were stained in Lugol's solution (Applichem) for 2 minutes and transferred into a chloral hydrate solution (chloral hydrate 8 g, glycerol 1 ml, water 2 ml). Images were observed using an AXIO image.M1 or an AXIO observer Z1 (Carl Zeiss AG, Oberkochen, Germany) combined with a CMOS camera (SPOT Idea 1.3 Mp Color Digital Camera, SPOT imaging, Sterling Heights, MI, USA). More than 5 plants per genotype were observed in an experiment and the experiment was repeated 2 times.

For analysis of above-ground organs of mature plants, images were acquired using a Nikon D80 (Nikon Co., Tokyo, Japan) digital camera. Images were taken after 15 and 30 (phyllotaxis and rosette development), 45 (inflorescence) and 50 (height) days growth on soil for shoot traits comparison between Col-0 and *EXO70A3*–related mutant. At least 9 plants were observed for each line and the experiment was repeated twice.

For analyses of fluorescent protein containing lines, crossbred lines (*DR5::GFP EXO70A3ox*, *DR5::GFP CRISPR-exo70A3-2*, *PIN1::PIN1:GFP EXO70A3ox*, *PIN2::PIN2:GFP EXO70A3ox*, *PIN3::PIN3:GFP EXO70A3ox*, *PIN4::PIN4:GFP EXO70A3ox*, *PIN7::PIN7:PIN7:GFP EXO70A3ox* and *PIN4::PIN4:GFP CRISPR-exo70A3-1*) and control lines for these crossbreds (*DR5::GFP*, *PIN1::PIN1:GFP*, *PIN2::PIN2:GFP*, *PIN3::PIN3:GFP*, *PIN4::PIN4:GFP* and *PIN7::PIN7:GFP*, respectively) were grown on 1 × MS agar plates for 5 days after germination. GFP signals in plant roots were observed under a laser scanning confocal microscope LSM700 (Carl Zeiss) with a 10× objective (EC plan Neofluar 10×0.3 M27), a 20× objective (Plan Apochromat 20×0.8 M27) or a 40× objective (Plan-Apochromat 40×/0.95 Korr M27, Zeiss). GFP-fusion proteins and PI were excited by a 488-nm diode array and detected by collecting light with wavelengths below 555 nm and above 560 nm, respectively. Gravity responses of GFP signals were observed using *DR5::GFP EXO70A3ox*, *DR5::GFP CRISPR-exo70A3-1* and control *DR5::GFP* lines following the dynamic root gravitropism assay (see below) and images were acquired 3 hours after re-orientation of plates. Cell outlines were visualized by propidium iodide (red signals). At least 10 plants per genotype were observed for an experiment and the experiments were repeated at least twice. FRAP (Fluorescent recovery after photo bleaching) was conducted using *PIN4::PIN4:GFP* and *PIN4::PIN4:GFP EXO70A3ox* lines by Visitron Spining Disc microscope, which is based on Nikon Eclipse Ti E inverted microscope (Nikon Instech Co., Ltd., Tokyo, Japan) equipped with CSU-W1 spinning disc unit (Yokogawa Electric Corporation, Tokyo, Japan), with a 60× water objective (Plan Apo VC 60x/1.2 WI water, Nikon) and Immersol W$^{TM}$ (Carl Zeiss). Plants were grown for 5 DAG and whole seedlings were mounted on glass slides with distilled water. FRAP observation was done using EMCCD (Ixon Ultra 888, Andor Technology Ltd., Belfast, UK) as follows: laser 488 nm (20% for observation, 100 % for bleaching), emission filter: ET 525/50, exposure 500 ms, FRAP time 150 ms/pixel, interval 0.5 minutes, pre-count 3 times, total time 20 minutes and EMCCD gain 645 nm. Fluorescent intensity was manually measured using Fiji. A ratio of fluorescent intensity to a neighboring cell, which was not photobleached, is calculated for each intensity data and then normalized by an average ratio of all pre-counts of *PIN4::PIN4:GFP* and used for the analysis. At least 10 plants were used in an experiment and the experiment was repeated twice.

**Dynamic root and shoot gravitropism assays**

To observe the dynamic shoot gravitropic responses (negative gravitropism), *EXO70A3*-related mutants and control Col-0 plants were grown on soil for 40 days. The whole pots were turned by 90° and images of the time-dependent angle changes of floral stems were acquired every hour for 5 hours using a Nikon D80 camera. Quantification of the angles of floral stems relative to gravity on these images was conducted using Fiji. At least Four plants were used for each line, and the experiment was repeated twice.

For the dynamic root gravitropism assay (positive gravitropism), plants were grown on 1 × MS agar plates for 5 days after germination under normal growth conditions. Plants of the same size were transferred to new vertical 1 × MS agar plates and incubated for a further one hours under the same growth conditions. These plates were then turned 90 degrees (in a manner that plates remained vertical) and fixed on vertically oriented scanners (EPSON Perfection V600 Photo). Images were acquired every four minutes for 20 hours under the same growth conditions. Root re-orientation (root tip angle change to the vector of gravity on images representing 32 minute intervals) was measured using Fiji as Root tip angle. 10 or more roots were measured for each genotype and the experiment was repeated at least three times. To compare the dynamic root gravitropism among accessions, we used eight accessions (haplogroup C: Ler-1, Shakdara (Sha), Lp2-2 and Per-1; haplogroup H: LAC-5, MIB-60, Coc-1 and Ty-0).

**Root system architecture (RSA) assay**

To assess the RSA in soil, we used three plants from each genotype. The experiment was conducted blind so that the plant genotypes were not known to the researcher conducting and analyzing the experiment, except for *pin4-3* and *CRISPR-exo70A3-1*. Each plant was grown on soil for 30 days in a pot ($\phi$ 20 cm, 17 cm height, Fiskars Germany GmbH, Herford, Germany) at 21°C, 16 hours light/16°C, 8 hours dark conditions. The whole pot was then sectioned in two halves at the center of the plant using a band saw machine (STARTRITE 20RWS, Startrite, A.LT. Saws & Spares Ltd., Gillingham, UK) equipped with a specially modified band (10 mm BAND SAW, LENOX Newell Rubbermaid

Inc., East Longmeadow, MA, USA). The soil at the section was gradually removed with a paint brush, and an image was acquired by a Nikon D80 camera. Using Fiji, the roots on the image were marked (overlay) and subsequently the coordinates of all root pixels were determined. The Shallowness (standard deviation of all root pixels in X axis direction in a picture divided by that in Y axis direction) was calculated using these data as an indicator of shallow or deep RSA. Note that, while many recent studies used rhizotron (Devienne-Barret *et al.* 2006, Plant and Soil) and related systems, we could not use these for our analysis. Images acquired by the rhizotron system, GLO-Roots system (Rella´n-A´lvarez *et al.* 2015, eLife) and other related systems are optically appealing and probably useful to understand morphological patterning of roots. However, all these systems do not allow roots grow in 3-dimensions (only in 2-dimensions, sometimes with a rounded shape plane) due to their system requirements for root visualization, particularly in case of Arabidopsis roots. Since roots in nature do not grow in 2-dimensions, these systems have severe limitations in studying natural root system architectures. In fact, no Col-0 root system architectures grow vertically deeper in 3-dimensional growth space while the rhizotron and related systems always showed vertically expanded roots (they do not have the space to grow anywhere else but down in an space that is essentially 2-dimensional).

**Seed number assay**

To analyze fitness of different genotypes in drought conditions, we conducted an assay similar to that reported previously (Schmalenbach *et al.*, 2014). In particular, we used four plants from each genotype and four sets of the whole drought experiment, resulting in a total number of 16 plants per genotype. Each experiment contained five pots that were arranged in a block in a growth chamber. In each block, positions of five pots were randomized. The experiment was conducted blindly so that the plant genotypes were not known to the researcher conducting and analyzing the experiment. Each four plants of a line were grown on soil for more than 60 days in a pot ($\phi$ 20 cm, 17 cm height, Fiskars Germany GmbH, Herford, Germany) at 60% humidity, 21°C, 16 hours light/16°C, 8 hours dark conditions until the seed production was completed. Four plants of two different genotypes were grown within one pot. Before starting the experiment, the initial weight of each pot filled with humid soil was measured. Subsequently, the soil in pots without plants was dried up by heating to 95 °C for 8 days and the initial water contents (% g water/g soil) of the soil was determined. Then water content of each pot was calculated with the total soil weight and the initial dry soil weight. During plant growth, water contents of pots were decreased from 60% (DAG5) to 30% (DAG 35). After that, pots were watered every two to three days as necessary by spray from top to keep the water contents 25 %. As the water-sufficient control, same experiment was repeated while water level was kept 65% through the whole experimental period in this control. Mature siliques from each plant were harvested and the number of seeds in five siliques was counted to determine the average number of seeds per silique for each plant. Subsequently, the total silique number of each plant was counted and the total seed number was decided using the average seed number per silique and the total silique number. An average seed number of four plants of each line in a pot was calculated, rounded and used in statistical analyses.

The significance of effects from seed number experiments in terms of responses to drought was determined using Poisson regression models. For the Col-0 versus *EXO70A3ox* experiment, the following model was fit using the glm function in the stats package in R:

$$\text{Seed number} \sim \mu + \text{Treatment} + \text{Genotype} + \text{Treatment*Genotype}$$

Where Seed number is the number of seeds per plant, Treatment refers to the watering regime, Genotype is Col-0 or *EXO70A3ox*, and Treatment*Genotype is the interaction between these two variables. In this model, all terms are significant as shown below.

| Coefficients[a] | | | | | |
|---|---|---|---|---|---|
| | Estimate | Standard error | z value | Pr (>\|z\|) | |
| (Intercept) | 11.070813 | 0.001972 | 5613.143 | $< 2.0 \times 10^{-16}$ | *** |
| Treatment-Drought | -3.673558 | 0.012535 | -293.067 | $< 2.0 \times 10^{-16}$ | *** |
| GWAS_ID-*EXO70A3ox* | -0.406473 | 0.003119 | -130.304 | $< 2.0 \times 10^{-16}$ | *** |
| Treatment-Drought*GWAS_ID-*EXO70A3ox* | -0.151038 | 0.02075 | -7.279 | $3.36 \times 10^{-13}$ | *** |

[a]Significance codes: 0, '***', 0.001, '**', 0.01, '*', 0.05, '.', 0.1, ' ', 1

For the experiment comparing haplogroups C and H of *EXO70A3*, the following mixed-effect Poisson model was fit using the glmer function in the lme4 package for R:

Seed number ~ $\mu$ + Treatment + Haplogroup + Treatment*Haplogroup + 1|Accession + $\varepsilon$

Where seed number is the number of seeds per plant, Treatment is a fixed effect referring to the watering regime, Haplogroup is a fixed effect representing the H or C haplogroup, and Accession is a random effect fitted for each individual accession. Treatment*Haplogroup is the interaction between these two variables. In this model, the Treatment and Treatment*Haplogroup interaction were significant, while the marginal Haplogroup effect was not as shown below.

| Coefficients[a] | | | | | |
|---|---|---|---|---|---|
| | Estimate | Standard error | z value | Pr (>\|z\|) | |
| (Intercept) | 10.61369 | 0.23522 | 45.1 | $< 2.0 \times 10^{-16}$ | *** |
| Treatment-Drought | -3.53378 | 0.00686 | -515.1 | $< 2.0 \times 10^{-16}$ | *** |
| Haplogroup-H | 0.08876 | 0.33269 | 0.3 | 0.79 | *** |
| Treatment-Drought*Haplogroup-H | -1.19095 | 0.01409 | -84.5 | $< 2.0 \times 10^{-16}$ | *** |

[a]Significance codes: 0, '***', 0.001, '**', 0.01, '*', 0.05, '.', 0.1, ' ', 1

In order to apply a population structure correction to the comparison of these haplogroups, a Poisson regression model was fit using the glm function in the stats package in R:

Seed number ~ $\mu$ + Treatment + Accession + Treatment*Accession

Where Seed number is the number of seeds per plant, Treatment refers to the watering regime, Accession is the accession tested, and Treatment*Accession is the interaction between these two variables. For each accession, the coefficients for these effects were extracted from the model output and used to calculate the following "modeled" phenotypes as shown below: seed number (SN) in well-watered conditions, seed number in drought conditions, and response to drought.

| GWAS_ID | SN (well-watered) | Response to drought | SN (drought) | Haplogroup |
|---|---|---|---|---|
| 6932 | 9.692257 | -2.48811 | 7.204149 | H |
| 6962 | 10.45793 | -2.91863 | 7.539293 | H |
| 7520 | 11.19351 | -4.56579 | 6.627711 | H |
| 8354 | 11.06028 | -3.7566 | 7.303675 | H |
| 204 | 10.82579 | -3.90585 | 6.919931 | C |
| 5729 | 10.90723 | -4.68564 | 6.221584 | C |
| 7351 | 10.11186 | -7.43771 | 2.674149 | C |
| 96 | 10.96822 | -6.07412 | 4.894101 | C |

A population-structure corrected test of differences between the haplogroups H and C was then performed for each of these phenotypes as described in "Genome wide association studies (GWAS) and bioinformatics analyses" part on Supplemental Experimental Procedure page 2. The resulting *p*-values for these tests are shown below.

Control: 0.844

Drought: 0.11051

Response: 0.09257

The significance of the differences in absolute seed numbers produced per plant in drought between Col-0 and *EXOCYST70A3ox* and between haplogroups C and H was analyzed by ANOVA.

**Quantitative reverse transcription PCR (qRT-PCR)**

For qRT-PCR, primers were purchased from Eurofins MWG Operon Co. (Ebersberg, Germany). Plants were grown on 1 × MS agar plates under normal growth conditions with or without 100 nM IAA for 7 days after germination. Tissues were collection by excision with fine scissors. Whole shoots and roots were simply cut from whole plants. For root tips, 2.5 mm root tips were excised from primary roots. 5 plants for the whole plant, 5 plants for the shoot, 10 plants for the root and more than 100 plants for the root tip samples were used. Samples were immediately frozen in liquid nitrogen. Frozen plant materials were ground using a ball mill (MM301, Retch, Haan, Germany) with 5 mm glass beads and total RNA was extracted using RNeasy Plant Mini (QIAGEN GmbH, Hilden, Germany). RNA concentrations were quantified using Nanodrop (Thermo Fisher Scientific Inc., Waltham, MA USA). cDNA was synthesized using RevertAid Reverse Transcriptase (Thermo Fisher Scientific Inc.) and Oligo $(dT)_{18}$ primer (Thermo Fisher Scientific Inc.) according to the manufacturer's protocol. qrt-PCR reaction was prepared using 2x SensiMix™ SYBR & Fluorescein Kit (PEQLAB LLC, Wilmington, DE, USA) according to the manufacturer's instructions and PCR was conducted by a combination of iCycler and IQ5 (Bio-Rad Laboratories, Inc., Hercules, CA, USA). All transcript levels were normalized to the tubulin β gene (AT5G62690). For AT5G52340 and AT5G52360, gene expression levels were calculated as relative values to that of Col-0 whole plants. For the other genes, expression levels were compared as the relative values to that in Col-0 root tips. *EXO70A3* gene expression in IAA treated plants was calculated as the relative value to the transcript level in mock treated (1% DMSO) Col-0 root tips. All experiments were repeated at least twice. Genes quantified include *EXO70A3* (AT5G52350), AT5G52340, AT5G52360, PIN1 (AT1G73590), PIN2 (AT5G57090),

PIN3 (AT1G70940), PIN4 (AT2G01420) and PIN7 (AT1G23080). For the comparison of *EXO70A3* expression levels among natural accessions, we used eight accessions (haplogroup C: Ler-1, Shakdara (Sha), Lp2-2 and Per-1; haplogroup H: LAC-5, MIB-60, Coc-1 and Ty-0).
